# Dendritic normalisation improves learning in sparsely connected artificial neural networks

**DOI:** 10.1101/2020.01.14.906537

**Authors:** Alex D Bird, Hermann Cuntz

## Abstract

Inspired by the physiology of neuronal systems in the brain, artificial neural networks have become an invaluable tool for machine learning applications. However, their biological realism and theoretical tractability are limited, resulting in poorly understood parameters. We have recently shown that biological neuronal firing rates in response to distributed inputs are largely independent of size, meaning that neurons are typically responsive to the proportion, not the absolute number, of their inputs that are active. Here we introduce such a normalisation, where the strength of a neuron’s afferents is divided by their number, to various sparsely-connected artificial networks. The learning performance is dramatically increased, providing an improvement over other widely-used normalisations in sparse networks. The resulting machine learning tools are universally applicable and biologically inspired, rendering them better understood and more stable in our tests.

## 1 Introduction

Artificial neural networks have had a huge impact over the last couple of decades, enabling substantial advances in machine learning for fields such as image recognition^1^, translation^2^, and medical diagnosis^3^. The inspiration for these tools comes from biological neuronal networks, where learning arises from changes in the strength of synaptic connections between neurons through a number of different plasticity mechanisms^4–7^. The growth of neural networks away from the limitations of biological networks, for example the use of global backpropagation algorithms that have access to information unavailable to real synaptic connection^8^, has meant that state-of-the-art artificial intelligence algorithms differ fundamentally from the biological function of the brain. Nevertheless, a number of biophysical principles have been successfully reintroduced, using salient features of real neuronal networks to make advances in the field of artificial neural networks^9–14^. Here we show how the dendritic morphology of a neuron, which influences both its connectivity and excitability, produces an afferent weight normalisation that improves learning in such networks.

Real neurons receive synaptic contacts across an extensively branched dendritic tree. Dendrites are leaky core conductors, where afferent currents propagate along dendritic cables whilst leaking across the cell membrane^15^. Larger dendrites increase the number of potential connections a cell can receive, meaning that more afferent currents can contribute to depolarisation^16^. Conversely, larger cells typically have lower input resistances, due to the increased spatial extent and membrane surface area, meaning that larger synaptic currents are necessary to induce the same voltage response and so bring a neuron to threshold^17;18^. It has recently been shown theoretically by Cuntz et al (2019)^19^ that these two phenomena cancel each other exactly: the excitability of neurons receiving distributed excitatory synaptic inputs is largely invariant to changes in size and morphology. In addition, neurons possess several active mechanisms to help maintain firing-rate homeostasis through both synaptic plasticity regulating inputs^20;21^ and changes in membrane conductance regulating responses^22^. These results imply a consistent biophysical mechanism that contributes to stability in neuronal activity despite changes in scale and connectivity.

Changing connectivity has traditionally not played much of a role in feedforward artificial neural networks, which typically used fully-connected layers where each neuron can receive input from all cells in the preceding layer. Sparsely-connected layers have however long been used as alternatives in networks with a variety of different architectures^9^. Sparse connectivity more closely resembles the structure of real neuronal networks and a number of recent studies have demonstrated that larger, but sparsely-connected, layers can be more efficient than smaller fully-connected layers both in terms of total parameter numbers and training times^23–25^. The advantage in efficiency comes from the ability to entirely neglect synaptic connections that do not meaningfully contribute to the function of the network. Sparse networks are also less likely to be overfitted to their training data as sparse representations of inputs are forced to focus on essential features of the signal instead of less-informative noise.

To produce an appropriate sparse connectivity a number of regularisation techniques have been suggested; *L*^1^- and *L*^0^-regularisations^23;26^ both penalise (the latter more explicitly) the number of connections between neurons during training. Mocanu et al (2018)^24^, building on previous work^27–29^, have recently introduced an evolutionary algorithm to reshape sparse connectivity, with weak connections being successively excised and randomly replaced. This procedure causes sparse artificial networks to develop small-world and scale-free topologies similar to biological neuronal circuits^30^ and has comparable performance to fully-connected layers despite having many fewer parameters to optimise.

Normalisation is another feature that has previously been shown to enhance learning in neural networks. In particular Ioffe & Szegedy (2015)^31^ introduced batch normalisation, where the inputs over a given set of training data are normalised, and Salimans & Kingma (2016)^32^ introduced *L*^2^-normalisation, where afferent synaptic weights are normalised by their total magnitude. The latter is reminiscent of heterosynaptic plasticity, where afferent synapses across a neuron depress in response to potentiation at one contact in order to maintain homeostasis^21^. Both techniques have been applied in fully-connected networks and both work to keep neuronal activities in the region where they are most sensitive to changes in inputs. The normalisation that arises from the relationship between a real neuron’s morphology and connectivity sits in this context and provides a particularly powerful, and biologically realistic, way to normalise sparse inputs. Dividing the magnitude of individual synaptic weights by their number distributes activity across neurons whilst keeping each cell sensitive to changes in inputs; neurons therefore encode signals from the proportion of presynaptic partners that are active, providing a simple and broadly applicable technique to ensure faster convergence to optimal solutions.

## 2 Methods and Models

Detailed descriptions of the datasets and networks used are given in captions and included as **Supplementary File 1**. In all cases traditional stochastic gradient descent^34;35^ is used with a minibatch size of 10 and a learning rate *η* of 0.05. Of particular importance is the sparse evolutionary training (SET) algorithm introduced by Mocanu et al (2018)^24^. Briefly, sparse connections are initiated uniformly randomly with probability *ε* to form an Erdoős-Rényi random graph^36^. After each training epoch, a fraction *ζ* of the weakest contacts are excised and an equal number of new random connections are formed. New connections are distributed normally with mean 0 and standard deviation 1.

All code is written in Python 3.6 and is included as **Supplementary File 2**. The networks in **Figures 1** and **2** are coded from first principles using the standard Numpy package, and the networks in **Figure 3** make use of Keras with a TensorFlow backend (`keras.io`). The application of dendritic normalisation in Keras with TensorFlow allows for immediate inclusion in Keras-based deep learning models. The normalisation requires a custom layer, constraint, and optimiser.

**Figure 1:**
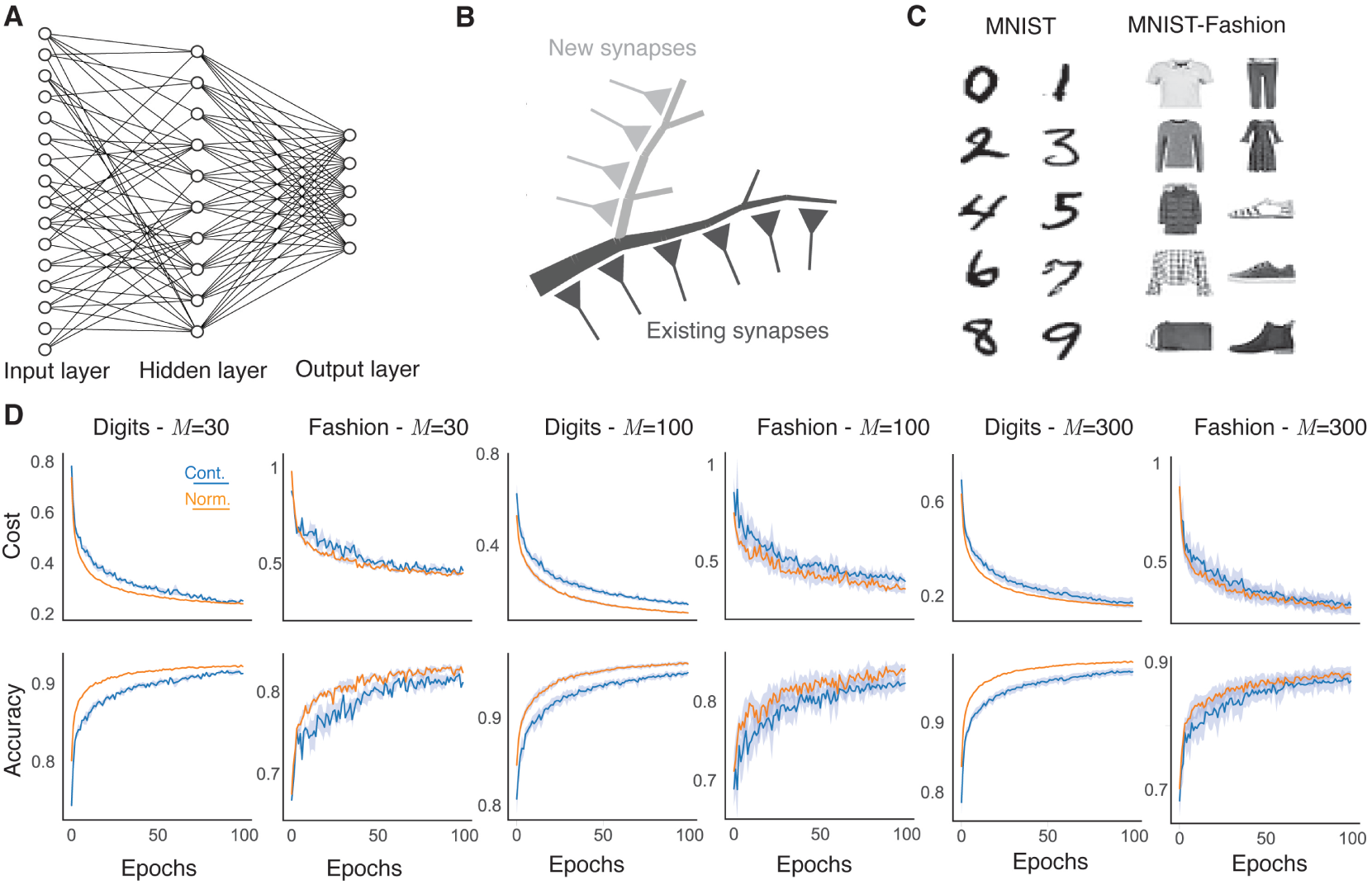
Dendritic normalisation improves learning in sparse artificial neural networks. **A**, Schematic of a sparsely-connected artificial neural network. Input units (left) correspond to pixels from the input. Hidden units (centre) receive connections from some, but not necessarily all, input units. Output units (right) produce a classification probability. **B**, Schematic of dendritic normalisation. A neuron receives inputs across its dendritic tree (dark grey). In order to receive new inputs, the dendritic tree must expand (light grey), lowering the intrinsic excitability of the cell through increased membrane leak and spatial extent. **C**, Example 28 *×* 28 pixel greyscale images from the MNIST^35^ (left) and MNIST-Fashion^37^ (right) datasets. The MNIST images are handwritten digits from 0 to 9 and the MNIST-Fashion images have ten classes, respectively: T-shirt/top, trousers, pullover, dress, coat, sandal, shirt, sneaker, bag, and ankle boot. **D**, Learning improvement with dendritic normalisation (orange) compared to the unnormalised case (blue). Top row: Log-likelihood cost on training data. Bottom row: Classification accuracy on test data. From left to right: digits with *M* = 30 hidden neurons, fashion with *M* = 30, digits with *M* = 100, fashion with *M* = 100, digits with *M* = 300, fashion with *M* = 300. Solid lines show the mean over 10 trials and shaded areas the mean ± one standard deviation. SET hyperparameters are *ε* = 0.2 and *ζ* = 0.15.

**Figure 2:**
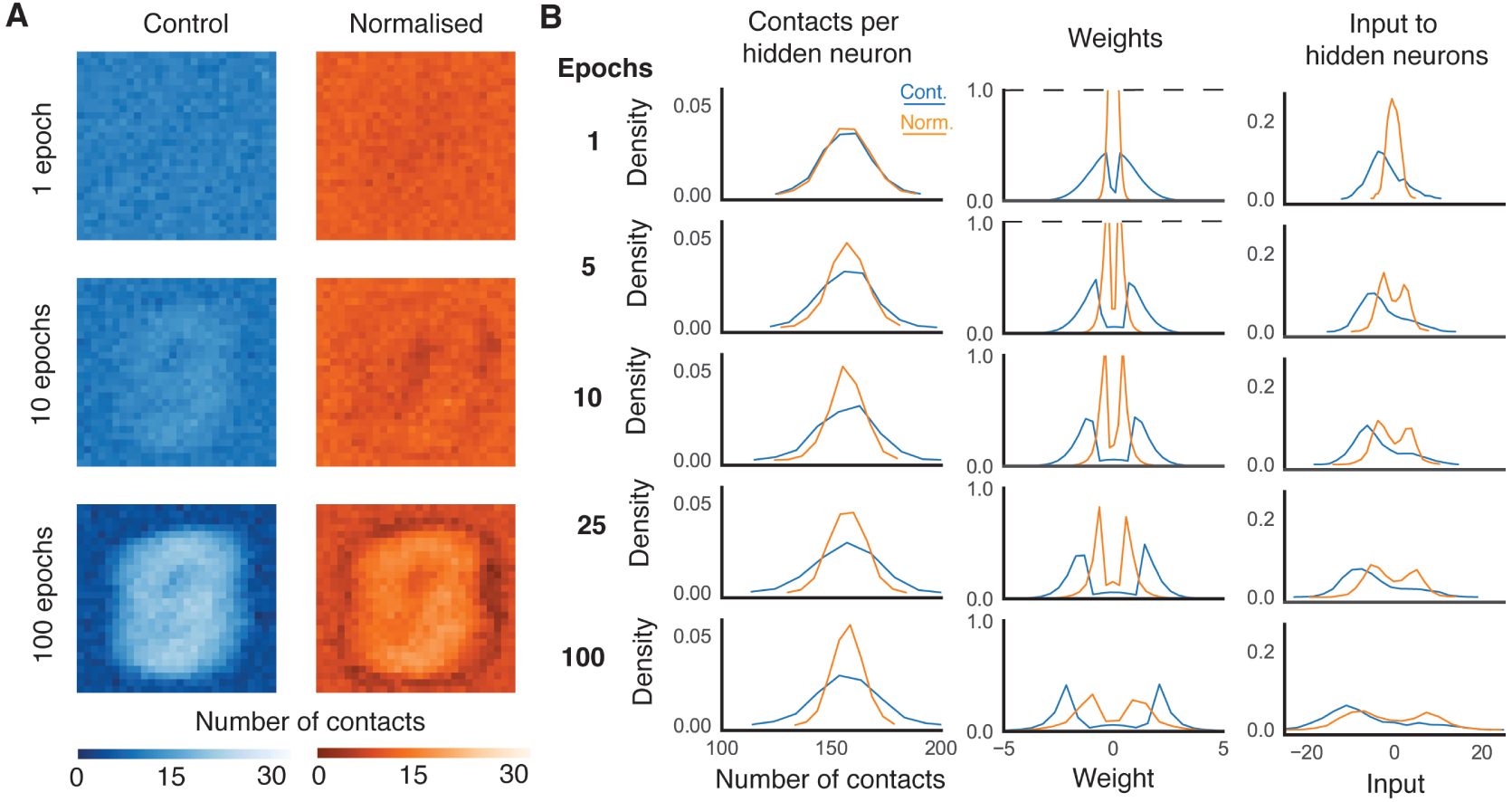
Evolution of synaptic weights. **A**, Number of efferent contacts from each input neuron (pixel) to neurons in the hidden layer as the weights evolve. The left panels (blue) show the unnormalised case and the right (orange) the normalised case. **B**, Afferent contacts for the unnormalised (blue) and normalised (orange) cases. From left to right: Distribution of the number of afferent contacts arriving at each hidden neuron, weights, and mean weighted input to each hiddden neuron over the test set. All panels show the average over 10 trials on the original MNIST dataset with hyperparameters *M* = 100, *ε* = 0.2, and *ζ* = 0.15. Dashed lines show where the vertical axis has been truncated to preserve the scale.

**Figure 3:**
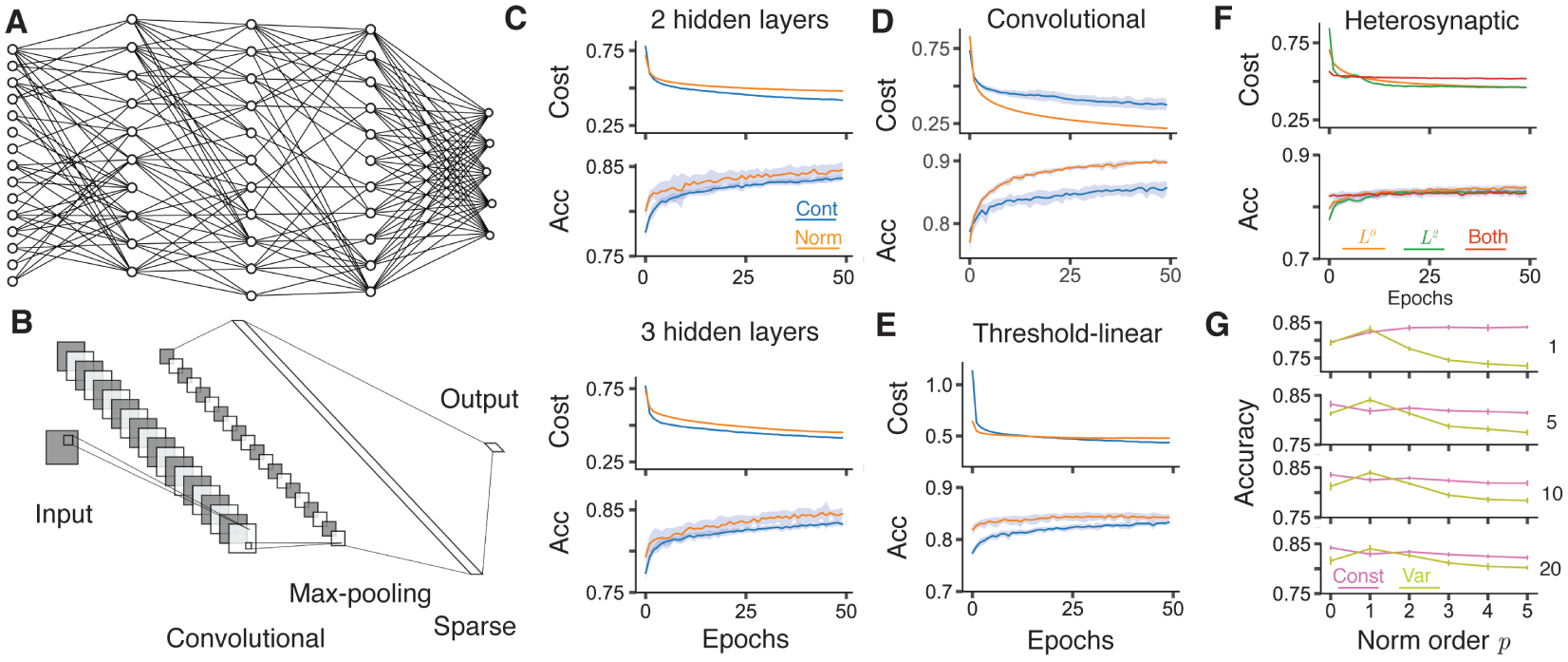
Improved training in deeper networks. **A** Schematic of a sparsely connected network with 3 hidden layers. The output layer is fully connected to the final hidden layer, but all other connections are sparse. **B** Schematic of a convolutional neural network^33^ with 20 5 *×* 5 features and 2 *×* 2 maxpooling, followed by a sparsely connected layer with *M* = 100 neurons. **C** Learning improvement with dendritic normalisation (orange) compared to the unnormalised control case (blue) for networks with 2 (top) and 3 (bottom, see panel **A**) sparsely-connected hidden layers, each with *M* = 100 neurons. Top of each: Log-likelihood cost on training data. Bottom of each: Classification accuracy on test data. **D** Improved learning in the convolutional network described in **B** for an unnormalised (blue) and normalised (orange) sparsely-connected layer. Top: Log-likelihood cost on training data. Bottom: Classification accuracy on test data. **E** Improved learning in a network with one hidden layer with *M* = 100 threshold-linear neurons for unnormalised (blue) and normalised (orange) sparsely-connected layers. Top: Log-likelihood cost on training data. Bottom: Classification accuracy on test data. **F** Comparison of dendritic (orange), heterosynaptic (green^32^), and joint (red, Eq 3) normalisations. Top: Log-likelihood cost on training data. Bottom: Classification accuracy on test data. **G** Comparison of test accuracy under different orders of norm *p* after (from top to bottom) 1, 5, 10, and 20 epochs. Pink shows constant (Eq 5) and olive variable (Eq 6) excitability. Solid lines show the mean over 20 trials and shaded areas and error bars the mean ± one standard deviation. All results are on the MNIST-Fashion dataset. Hyperparameters are *ε* = 0.2 and *ζ* = 0.15.

## 3 Results

### 3.1 Dendritic normalisation and stochastic gradient descent

Let **w**_*i*_ be the input weight vector to a given neuron *i*. Then the dendritic normalisation can be written as

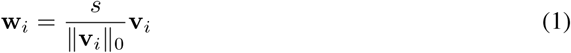

where **v**_*i*_ is a vector of the same size as **w**_**i**_, ‖**v‖** _**0**_ is the *L*^0^-norm of **v**_*i*_ (the number of non-zero elements), and *s* is a scalar that determines the magnitude of the vector **w**_*i*_. This normalisation arises from the fact that, for real neurons, increased synaptic connectivity is balanced by the additional leak conductance and spatial extent of the new dendritic cables required to collect the inputs (**Figure 1b**). The parametrisation here differs from that introduced by Salimans & Kingma (2016)^32^ for fully-connected networks with Euclidean normalisation in two fundamental ways. Firstly, the magnitude parameter *s* is the same across all neurons as it reflects a conserved relationship between connectivity and excitability. If a network were to include distinct classes of artificial neurons with distinct synaptic integration properties, different values of *s* may be appropriate for each class, but should not differ between neurons of the same class. Secondly, the *L*^0^-norm ‖v_i_‖_0_ is distinct from the Euclidean *L*^2^-norm in that it is almost surely constant with respect to **v**_*i*_: Connections are not created or destroyed by stochastic gradient descent.

The gradients of the cost function *C* with respect to **v**_*i*_ and *s* can be written as

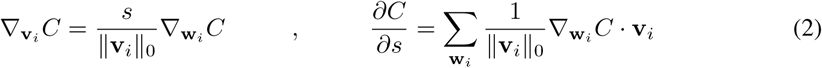

where 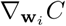 is the usual gradient of *C* with respect to the full weight vector **w**_*i*_ and the sum in the second equation is over all weight vectors in the network. An interesting consequence of these equations is that neurons with more afferent connections will have smaller weight updates, and so be more stable, than those with fewer afferent connections.

### 3.2 Dendritic normalisation improves learning in neural networks

We consider the performance of sparse neural networks with and without dendritic normalisation on the MNIST and MNIST-Fashion datasets (**Figure 1**). For sparse networks using sparse evolutionary training (SET) with one hidden layer consisting of 30, 100, and 300 neurons, with connection probability *ε* = 0.2 and SET excision rate *ζ* = 0.15, the normalised network consistently learns faster than the unnormalised control network (orange lines against blue lines). This result holds across both the cost on the training sets and the accuracy on the test sets for both datasets and across all network sizes, indicating a robust improvement in performance. In addition, the variability between different independent training regimes (shaded areas in **Figure 1d** show mean plus or minus one standard deviation over 10 independently initiated training regimes) is reduced, dendritic normalisation therefore also makes training more robust and reliable.

### 3.3 Evolution of connectivity

It is possible to visualise the connection structure that results from training the control and normalised networks on the MNIST data (**Figure 2**). **Figure 2a** shows how the number of efferent connections from each input neuron (organised as pixels) changes with the number of training epochs. Initially, the connections are randomly distributed and have no spatial structure, but the SET algorithm gradually imposes a heavier weighting on the central input neurons as training progresses. This feature was shown before by Mocanu et al (2018)^24^ as central pixels are likely to be more informative over the relevant datasets. Comparing the control (left, blue) and normalised (right, orange) networks, it is interesting to note that the bias towards central pixels is less strong in the normalised case: Input neurons over a relatively broad area are strongly connected to the hidden layer.

Postsynaptically, the number of contacts received by each hidden neuron is less variable in the normalised case (**Figure 2b**, left column), the weights of these contacts are typically smaller in absolute value and less dispersed (**Figure 2b**, central column); the resultant weighted inputs over the test data to hidden neurons are therefore more consistent (**Figure 2b**, right column). The normalisation appears to make better use of neural resources by distributing connections and weights more evenly across the available cells, whilst keeping expected inputs closer to 0, the steepest part of the activation function, where responses are most sensitive to inputs. In addition, the smaller connection weights suggest that normalised networks may be even more robust to overfitting than the equivalent unnormalised sparse network. This is supported by the increased improvement in the case of more complex networks, both in terms of more hidden units and more layers, as well as the greater improvement in evaluation accuracy compared to training cost (**Figures 1 and 3**).

### 3.4 Improved training in deeper networks

The improvement in learning performance generalises to deeper networks with multiple hidden layers (**Figure 3a**, as well as convolutional networks with a sparsely connected layer following the max pooling layer (**Figure 3b**). In all cases, learning is improved by dendritic normalisation (orange versus blue lines in **Figures 3c** and **d** for the MNIST-Fashion dataset) in a similar manner to that for the single-layered networks described above. Indeed, the improvement seems to increase with the complexity of the architecture. Dendritic normalisation is therefore applicable, and beneficial, as a universal technique for sparse layers in deep networks. The improvement in performance is not limited to artificial neurons with a sigmoid activation function. When neurons instead have non-saturating threshold linear activations the dendritic normalisation again improves learning (**Figure 3e**).

### 3.5 Comparison with other normalisations

Dendritic normalisation is just one of many mechanisms to promote and stabilise learning and activity in real neurons. Combining the *L*^0^-normalisation introduced here with the *L*^2^-normalisation of Salimans & Kingma (2016)^32^ provides a demonstration of the joint effects of passive dendritic constancy and compensatory heterosynaptic plasticity within an artificial neuron. Here, inputs to a cell are normalised by both number and magnitude

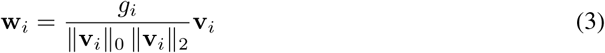

where 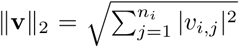 is the *L*^2^-norm of **v**_*i*_. The uniform excitability parameter *s* is now contained in the individual magnitudes *g*_*i*_ that differ for each cell. The gradients of the cost function *C* with respect to **v**_*i*_ and *g*_*i*_ can now be written as

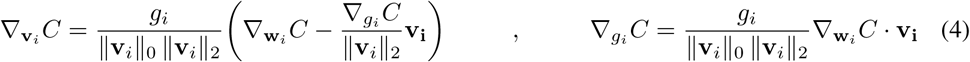

With both normalisations applied, well-connected neurons will again tend to change afferent weights more slowly, as will cells with lower excitability *σ*_*i*_. In consequence, cells receiving a few strong connections will be relatively fast-learning and unstable compared to those that receive many, individually less effective, connections.

The effects of dendritic (orange) and heterosynaptic (green) normalisations can be seen in **Figure 3f**. Here, a single layer of 100 hidden neurons is trained on the MNIST-Fashion data. Both normalisations individually have similar performance, with a slight advantage for the dendritic normalisation (orange): the mean test accuracy after 50 epochs is 83.75% compared to 82.66% for the heterosynaptic normalisation over 20 trials, and the accuracy is consistently higher. Interestingly, the joint normalisation given by Eq 3 (red lines in **Figure 3f**) reaches almost its highest level of accuracy after a single epoch and does not improve substantially thereafter. As neurons with many weak connections learn relatively slowly when both normalisations are in place it appears that good use is made of existing connectivity, but that less informative connections are not selectively weakened enough to be excised by the SET algorithm. The joint mechanism appears well suited to sparse, but static, connectivity whilst lacking the power of individual normalisations, either dendritic or heterosynaptic, to identify less necessary synapses. Whilst it is possible to exaggerate the biological relevance of learning through backpropagation, it is interesting to note that the dendritic *L*^0^-normalisation is intrinsic to real neurons, whilst heterosynaptic *L*^2^-normalisation can be regulated^21^.

Biologically plausible learning rules often include normalisation of afferent weights to ensure stability and improve convergence. Both Oja’s rule^6^ and the unsupervised learning procedure introduced by Krotov & Hopfield (2019)^14^, for example, ensure convergence to *L*^*p*^-normalised afferent weights for *p ≥* 2. Whilst a direct comparison with such distinct learning rules is beyond the scope of this study, many such normalisations rely on the *L*^*p*^ weight norm for some value of *p* with either constant (*s* in Eq 1) or variable (*g*_*i*_ in Eq 3) excitability ratios. For *p ≥* 1 the general gradients provided by such *L*^*p*^-normalisations are

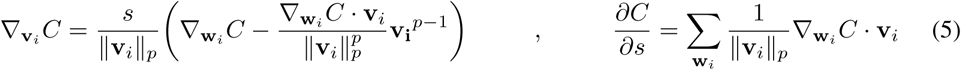

for constant excitability and

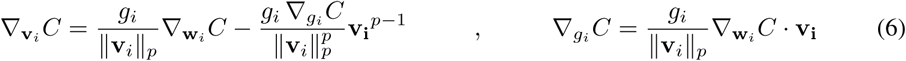

for variable excitability. In all cases, higher *L*^*p*^-norms imply slower learning; increasing the value of *p* means that the norms ‖**v**_*i*_ *‖*_*p*_ become increasingly insensitive to the smallest afferent weights.

**Figure 3g** shows the mean test accuracy for 100 hidden neurons on the MNIST-Fashion data after 1, 5, 10, and 20 training epochs as a function of the order *p* of normalisation for *p* = 0, 1, 2, 3, 4, and 5. The pink lines show the case of constant excitability (Eq 5) and the olive lines variable excitability (Eq 6). All normalisations show substantial improvement over the control case (**Figure 1d**) and performance is fairly similar across a broad range of parameters. The dendritic normalisation (*L*^0^-norm in pink) has the highest performance after 10 epochs, although the *L*^1^-norm with variable excitability is similar. Interestingly, for all orders except *p* = 1, the constant excitability case has better performance than the situation with variable excitability and this gap increases with *p*. In the constant excitability case, a single parameter *s* determines the postsynaptic response to afferent weights with a given norm, whereas in the variable case each neuron *i* has its own response *g*_*i*_ to normed inputs. It appears that maintaining a comparable excitability between neurons is superior in terms of learning to allowing this to vary between cells.

### 3.6 Performance on standard benchmarks

As a final test, we show that dendritic normalisation can enhance the accuracy of artificial neural networks on common benchmark datasets. We compare dendritically normalised sparse networks to the published results of comparable sparse networks in **Table 1**. It should be noted that the sparse results quoted in the literature are often in the context of improving the performance of fully-connected neural networks with the same architecture despite having many fewer parameters.In most cases, applying dendritic normalisation improves upon the published performance. The one exception is the COIL-100 data where the training/test split is random; using slightly different data to Pieterse & Mocanu (2019)^25^ could explain this difference.

**Table 1:** Table of performance for benchmark datasets compared to published results on sparse networks. We replicate the published architecture in each case: For the original MNIST dataset and CIFAR-10 datasets, Mocanu et al (2018)^24^ used three sparsely-connected layers of 1000 neurons each and 4% of possible connections existing. Pieterse & Mocanu (2019)^25^ used the same architecture for the COIL-100 dataset. For the Fashion-MNIST dataset, Pieterse & Mocanu (2019)^25^ used three sparsely-connected layers of 200 neurons each, with 20% of possible connections existing.

| Dataset | Size |  |  | Accuracy |  |
| --- | --- | --- | --- | --- | --- |
|  | Training | Test | Classes | Control (Source) | Normalisation |
| MNIST<sup>35</sup> | 60,000 | 10,000 | 10 | 98.74%<sup>24</sup> | **99.63%** |
| Fashion-MNIST<sup>37</sup> | 60,000 | 10,000 | 10 | 89.01%<sup>25</sup> | **92.23%** |
| CIFAR-10<sup>39</sup> | 50,000 | 10,000 | 10 | 74.84%<sup>24</sup> | **77.43%** |
| COIL-100<sup>40</sup> | 5,764 | 1,436 | 100 | **98.68%**<sup>25</sup> | 98.47% |

## 4 Discussion

We have shown that normalising afferent synaptic weights by their number (*L*^0^-normalisation), in a manner similar to real neurons^19^, improves the learning performance of sparse artificial neural networks with a variety of different structures. Such dendritic normalisation constrains the weights and expected inputs to be within relatively tight bands, potentially making better use of available neuronal resources. Neurons respond more to the proportion of their inputs that are active and highly-connected neurons are relatively less excitable. We believe that such a normalisation procedure is robust and should be applied to improve the performance of feedforward sparse networks.

Other results on normalisation^31;32^ have also demonstrated improvements in training performance in fully-connected feedforward networks. Such approaches work by keeping neurons relatively sensitive to changes in inputs and our results here can be seen as the sparse, and biologically justified, analogue, with similarly simple and broad applicability to the *L*^2^-normalisation introduced by Salimans & Kingma (2016)^32^. In situations of dynamic connectivity, the dendritic normalisation outperforms other techniques. The comparison between the heterosynaptic plasticity-like *L*^2^-normalisation and our dendritic *L*^0^-normalisation is particularly interesting. In real neurons the former relies on actively re-weighting afferent contacts^21^ whereas the latter arises purely from the passive properties of dendritic trees. Neurons often display complementary functionality between passive structure and active processes, for example in the equalisation of somatic responses to synaptic inputs at different locations both dendritic diameter taper^41^ and active signal enhancement^42^ play a role. Synaptic normalisation is in a similar vein. The effects are, however, distinct in some ways: while both normalisations keep neurons sensitive to inputs, the responses to learning differ. Heterosynaptic plasticity enhances changes in the relative weighting of contacts, whereas dendritic normalisation increases the stability of well-connected neurons while allowing faster learning in poorly connected cells (Eq 2). This makes dendritic normalisation particularly suited to situations with evolving connectivity.

In the context of biological realism, the normalisation here has room for development. We sought a straightforwardly demonstrable and quantifiable computational role for the dendritic normalisation described in Cuntz et al (2019)^19^ and so used the most well-developed theories within artificial neural networks^43^. Such networks have a number of features that are impossible to implement in the brain, so more investigation into the benefits of size-invariant excitability in living systems is necessary. Firstly, a single artificial neuron can form connections that both excite and inhibit efferent cells, a phenomenon which is not seen in the connectivity of real neurons^44^. It is possible to regard the mix of excitatory and inhibitory connections as a functional abstraction of real connections mediated by intermediate inhibitory interneurons^45^, but a more satisfying picture may emerge by considering distinct inhibitory populations of cells. Secondly, we train our networks using supervised backpropagation which employs global information unavailable to real synaptic connections. A large number of studies have developed more plausible local algorithms for training networks^6;7;11;13;14^. Such algorithms are a natural fit for our normalisation as it too is implemented biophysically through the morphology of the dendritic tree; an immediate goal is to incorporate our result into algorithms on unsupervised and reinforcement learning. Thirdly, our networks are strictly feedforward whereas typical neuronal networks possess recurrent connections. Many studies of such networks employ sparsity in recurrent excitatory connections^13;46^ and this is another natural arena for our normalisation; here the two streams of input would be in competition as more strongly recurrently connected cells would receive relatively weak feedforward inputs and vice versa. This is a particularly interesting avenue to explore. Fourthly, the neurons here are rate-based with either saturating or non-saturating outputs. Spiking networks can have different properties^10^ and spikes could be incorporated into any of the approaches described above.

Dendrites in general have much more to offer in terms of artificial neural computations. Synaptic connections are distributed spatially over branched dendritic trees, allowing for a number of putative computational operations to occur within a single cell^47^. Dendrites are able to passively amplify signals selectively based on their timing and direction^15^ and actively perform hierarchical computations over branches^48^. Cuntz et al (2019)^19^ noted that while mean neuronal firing rates are size-independent, the timing of individual spikes is not necessarily unaffected by morphology^18^. This means that signals encoded by rates are normalised whereas those encoded by spike timing may not be, implying that the two streams of information across the same circuit pathways are differentially affected by changing connectivity. Dendrites additionally hold continuous variables through their membrane potential and conductances that shape ongoing signal integration^49^. Such properties have potential computational roles that, while sometimes intuitive, have yet to be systematically studied at the level of neuronal circuits.

Overall, this study has two major consequences. The first is a practical normalisation procedure that drastically improves learning in sparse artificial neural networks trained using backpropagation. The procedure can be used effectively for any sparsely-connected layers in shallow or deep networks of any architecture. Given that sparse layers can display better scalability than fully-connected layers^24^, we believe that this procedure could become standard in deep learning. Furthermore, the biological plausibility of the procedure means that it is highly appropriate as a component of more physiologically realistic learning rules. Secondly, we have taken the insights from Cuntz et al (2019)^19^ and demonstrated a previously unappreciated way that the structure of dendrites, in particular their properties as leaky core conductors receiving distributed inputs, contributes to the computational function of neuronal circuits.

## Broader Impact

The results in this paper provide a theoretical advance that has two main components. The first is a practical normalisation technique that can be applied to sparse neural networks, increasing the efficiency and scalability of training such structures. The improved efficiency of training sparse networks is likely to enhance their utility and usage. In particular, the development of efficient sparse architectures could enable usage of deep learning techniques with less powerful processors and therefore broader adoption. The social impacts of this would not be qualitatively distinct from that of any general advance in machine learning.

The second component of our research is the demonstrated link between the passive spatial extent of biological neurons and learning. We feel that this is a particularly exciting development as it reinforces the link between the structure and computational function of neurons; we hope it will stimulate further research in this area.

## Supporting information

Expanded Methods.

## Acknowledgments and Disclosure of Funding

We would like to thank P Jedlicka, F Effenberger, and K Shapcott for helpful discussion. We acknowledge funding through BMBF grant 01GQ1406 (Bernstein Award 2013).

## Notes

### Competing Interest Statement

The authors have declared no competing interest.

