## Supplementary material for "Dendritic normalisation improves learning in sparsely connected artificial neural networks": Expanded Methods.

### 1 Expanded Methods

#### Network architectures

We first consider a simple artificial neural network (ANN) with one hidden layer to demonstrate the utility of our approach. The size of the input layer for both the MNIST and MNIST-Fashion datasets is 784, as each image is a $28 \times 28$ pixel greyscale picture. The hidden layer consists of $M$ neurons, each neuron $i$ receiving a number $n_i$ contacts from the previous layer. In **Figure 1**, $M = 30$, $100$, and $300$. In **Figures 2** and **3**, $M = 100$. Neuronal activation in the input and hidden layers as a function of input $z_i$ is controlled by a sigmoid function $\sigma(z_i)$

$$\sigma(z_i) = \frac{1}{1 + e^{-z_i}} \tag{1}$$

where $z_i$ is the weighted input to neuron $i$, given by

$$z_i = b_i + \sum_{n_i} w_{k,i} a_k \tag{2}$$

Here $b_i$ is the bias of each neuron $i$, $w_{k,i}$ is the synaptic weight from neuron $k$ in the previous layer to neuron $i$, and $a_k = \sigma(z_k)$ is the activation of presynaptic neuron $k$. The set of all $w_{k,i}$ for a given postsynaptic neuron $i$ form an afferent weight vector $\mathbf{w}_i$.

Both datasets have ten classes and the output of the ANN is a probability distribution assigning confidence to each possible classification. Neurons in the output layer are represented by softmax neurons where the activation function $\sigma_s(z_i)$ is given by

$$\sigma_s(z_i) = \frac{e^{z_i}}{\sum_{i=1}^{10} e^{z_i}} \tag{3}$$

The cost function $C$ is taken to be the log-likelihood

$$C = -\log(a_{\text{Correct}}) \tag{4}$$

where $a_{\text{Correct}}$ is the activation of the output neuron corresponding to the correct input.

For **Figure 3**, we generalise our results to deeper architectures and threshold-linear neuronal activations. In **Figures 3a** and **3c** we expand the above to include 2 and 3 sparse hidden hidden layers, each with $M = 100$ sigmoid neurons. In **Figures 3b** and **3c** we consider a simple convolutional neural network (LeCun et al, 2009) with 20 $5 \times 5$ features and $2 \times 2$ maxpooling. In **Figure 3d** we return to the original architecture with $M = 100$, but replace the sigmoid activation function $\sigma(z)$ for the hidden neurons with a non-saturating threshold-linear activation function $\tau(z)$ defined by

$$\tau(z_i) = \max(0, z) \tag{5}$$

Submitted to 34th Conference on Neural Information Processing Systems (NeurIPS 2020). Do not distribute.

In **Table 1**, we show the results of other architectures to match published performance benchmarks. We replicate the published architecture in each case: For the original MNIST dataset and CIFAR-10 datasets, Mocanu et al (2018) used three sparsely-connected layers of 1000 neurons each and $4\%$ of possible connections existing. Pieterse & Mocanu (2019) used the same architecture for the COIL-100 dataset. For the Fashion-MNIST dataset, Pieterse & Mocanu (2019) used three sparsely-connected layers of 200 neurons each, with $20\%$ of possible connections existing.

In all cases traditional stochastic gradient descent (Robbins & Monro, 1951; LeCun et al, 1998) is used with a minibatch size of 10 and a learning rate $\eta$ of $0.05$.

### Sparse evolutionary training (SET)

Connections between the input and hidden layers are established sparsely (**Figure1a**) using the sparse evolutionary training algorithm introduced by Mocanu et al (2018). Briefly, connections are initiated uniformly randomly with probability $\varepsilon$ to form an Erdős-Rényi random graph (Erdős & Rényi, 1959). After each training epoch, a fraction $\zeta$ of the weakest contacts are excised and an equal number of new random connections are formed. New connections are distributed normally with mean $0$ and standard deviation $1$.

### Datasets

The ANN is originally trained to classify $28 \times 28$ pixel greyscale images into one of ten classes. Two distinct datasets are initially used. The MNIST, introduced by LeCun et al (1998), consists of handwritten digits which must be sorted into the classes 0 to 9 (**Figure 1b**, left). The MNIST-Fashion dataset was introduced by Xiao et al (2017) as a direct alternative to the original MNIST and consists of images of clothing. The classes here are defined as T-shirt/top, trousers, pullover, dress, coat, sandal, shirt, sneaker, bag, and ankle boot (**Figure 1b**, right). Each dataset contains $60,000$ training images and $10,000$ test images. State-of-the-art classification accuracy for the original MNIST dataset is as high as $99.77\%$ (Cireşan et al, 2012), which likely exceeds human-level performance due to ambiguity in some of the images. For the newer MNIST-Fashion dataset state-of-the art networks can achieve classification accuracies of $96\%$. Such performance is achieved with deep network architectures, which we do not reproduce here, rather showing an improvement in training between comparable, and comparatively simple, artificial neural networks.

In Table 2, we also analyse other datasets. CIFAR-10 (Krizhevsky, 2012) contains $50,000$ training images and $10,000$ test images to be divided into the classes airplane, automobile, bird, cat, deer, dog, frog, horse, ship, and truck. Each image is $32 \times 32$ pixels in three colour channels. The COIL-100 dataset (**?**), which contains $7,200$ images in total, consists of images of 100 objects rotated in various ways. Each image is $128 \times 128$ pixels in three colour channels. There is no existing training/test split, so we follow Pieterse & Mocanu (2019) in randomly assigning $20\%$ of the available images to the test set.

### Code and data availability

All code is written in Python 3.6 and is freely available for download (see **Supplementary File 2**) alongside the MNIST and MNIST-Fashion datasets. These can also be downloaded from various places, including at the time of writing `yann.lecun.com/exdb/mnist/` and `github.com/zalandoresearch/fashion − mnist` respectively. The networks in **Figures 1** and **2** are coded using the standard Numpy package, and the networks in **Figure 3** make use of Keras with a TensorFlow backend (`keras.io`).

The application of dendritic normalisation in Keras with TensorFlow allows for immediate inclusion in Keras-based deep learning models. The normalisation requires a custom layer, constraint, and optimiser.

| Symbol | Interpretation |
|---|---|
| $a_i$ | Activation of neuron $i$ |
| $b_i$ | Bias of neuron $i$ |
| $C$ | Log-likelihood cost function) |
| $g_i$ | Excitability of neuron $i$ |
| $n_i$ | Number of afferent contacts to neuron $i$ (also written $\|\mathbf{v}_i\|_0$) |
| $s$ | (Uniform) Excitability of all neurons |
| $\mathbf{v}_i$ | Unnormalised input to neuron $i$ |
| $\mathbf{w}_i$ | Normalised input to neuron $i$ |
| $\varepsilon$ | SET connection probability |
| $\zeta$ | SET excision probability |
| $\eta$ | Learning rate for stochastic gradient descent |
| $\sigma$ | Sigmoid activation function |
| $\sigma_s$ | Softmax activation function |
| $\tau$ | Threshold-linear activation function |

**Table S1:** Table summarising symbols and interpretations.
