## Supplementary material for "Dendritic normalisation improves learning in sparsely connected artificial neural networks": Code and data.: Fig_1_data_gen.py

```
#!/usr/bin/env python3
# -*- coding: utf-8 -*-
"""
Created on Tue Nov 12 22:21:03 2019

@author: Alex
"""
###############################################################################
###############################################################################
###############################################################################
# Digits with 30 hidden neurons

n_epoch=100
nrep=10

pool = mp.Pool(mp.cpu_count()) # Multiprocessing         
multi_locs=[pool.apply(sp_func, args=([30],n_epoch,dig_training,dig_test)) for dummy_ind in range(nrep)] # Check all_funcs for defaults
pool.close()
sp_all_accs=np.zeros([nrep,n_epoch])
sp_all_costs=np.zeros([nrep,n_epoch])
for rep_ind in range(nrep):
    sp_all_accs[rep_ind]=multi_locs[rep_ind][0]
    sp_all_costs[rep_ind]=multi_locs[rep_ind][1]


pool = mp.Pool(mp.cpu_count()) # Multiprocessing         
multi_locs=[pool.apply(nm_func, args=([30],n_epoch,dig_training,dig_test)) for dummy_ind in range(nrep)] # Check all_funcs for defaults
pool.close()
nm_all_accs=np.zeros([nrep,n_epoch])
nm_all_costs=np.zeros([nrep,n_epoch])
for rep_ind in range(nrep):
    nm_all_accs[rep_ind]=multi_locs[rep_ind][0]
    nm_all_costs[rep_ind]=multi_locs[rep_ind][1]
    
    
np.save('sim_data/Fig_1_30_digits_sp_cst',sp_all_costs)
np.save('sim_data/Fig_1_30_digits_sp_acc',sp_all_accs)
np.save('sim_data/Fig_1_30_digits_nm_cst',nm_all_costs)
np.save('sim_data/Fig_1_30_digits_nm_acc',nm_all_accs)


#%%
###############################################################################
###############################################################################
###############################################################################
# Fashion with with 30 hidden neurons
pool = mp.Pool(mp.cpu_count()) # Multiprocessing         
multi_locs=[pool.apply(sp_func, args=([30],n_epoch,fash_training,fash_test)) for dummy_ind in range(nrep)] # Check all_funcs for defaults
pool.close()
sp_all_accs=np.zeros([nrep,n_epoch])
sp_all_costs=np.zeros([nrep,n_epoch])
for rep_ind in range(nrep):
    sp_all_accs[rep_ind]=multi_locs[rep_ind][0]
    sp_all_costs[rep_ind]=multi_locs[rep_ind][1]


pool = mp.Pool(mp.cpu_count()) # Multiprocessing         
multi_locs=[pool.apply(nm_func, args=([30],n_epoch,fash_training,fash_test)) for dummy_ind in range(nrep)] # Check all_funcs for defaults
pool.close()
nm_all_accs=np.zeros([nrep,n_epoch])
nm_all_costs=np.zeros([nrep,n_epoch])
for rep_ind in range(nrep):
    nm_all_accs[rep_ind]=multi_locs[rep_ind][0]
    nm_all_costs[rep_ind]=multi_locs[rep_ind][1]
    
    
np.save('sim_data/Fig_1_30_fashion_sp_cst',sp_all_costs)
np.save('sim_data/Fig_1_30_fashion_sp_acc',sp_all_accs)
np.save('sim_data/Fig_1_30_fashion_nm_cst',nm_all_costs)
np.save('sim_data/Fig_1_30_fashion_nm_acc',nm_all_accs)

#%%
###############################################################################
###############################################################################
###############################################################################
# Digits with 100 hidden neurons

pool = mp.Pool(mp.cpu_count()) # Multiprocessing         
multi_locs=[pool.apply(sp_func, args=([100],n_epoch,dig_training,dig_test)) for dummy_ind in range(nrep)] # Check all_funcs for defaults
pool.close()
sp_all_accs=np.zeros([nrep,n_epoch])
sp_all_costs=np.zeros([nrep,n_epoch])
for rep_ind in range(nrep):
    sp_all_accs[rep_ind]=multi_locs[rep_ind][0]
    sp_all_costs[rep_ind]=multi_locs[rep_ind][1]


pool = mp.Pool(mp.cpu_count()) # Multiprocessing         
multi_locs=[pool.apply(nm_func, args=([100],n_epoch,dig_training,dig_test)) for dummy_ind in range(nrep)] # Check all_funcs for defaults
pool.close()
nm_all_accs=np.zeros([nrep,n_epoch])
nm_all_costs=np.zeros([nrep,n_epoch])
for rep_ind in range(nrep):
    nm_all_accs[rep_ind]=multi_locs[rep_ind][0]
    nm_all_costs[rep_ind]=multi_locs[rep_ind][1]
    
    
np.save('sim_data/Fig_1_100_digits_sp_cst',sp_all_costs)
np.save('sim_data/Fig_1_100_digits_sp_acc',sp_all_accs)
np.save('sim_data/Fig_1_100_digits_nm_cst',nm_all_costs)
np.save('sim_data/Fig_1_100_digits_nm_acc',nm_all_accs)


#%%
###############################################################################
###############################################################################
###############################################################################
# Fashion with with 100 hidden neurons
pool = mp.Pool(mp.cpu_count()) # Multiprocessing         
multi_locs=[pool.apply(sp_func, args=([100],n_epoch,fash_training,fash_test)) for dummy_ind in range(nrep)] # Check all_funcs for defaults
pool.close()
sp_all_accs=np.zeros([nrep,n_epoch])
sp_all_costs=np.zeros([nrep,n_epoch])
for rep_ind in range(nrep):
    sp_all_accs[rep_ind]=multi_locs[rep_ind][0]
    sp_all_costs[rep_ind]=multi_locs[rep_ind][1]


pool = mp.Pool(mp.cpu_count()) # Multiprocessing         
multi_locs=[pool.apply(nm_func, args=([100],n_epoch,fash_training,fash_test)) for dummy_ind in range(nrep)] # Check all_funcs for defaults
pool.close()
nm_all_accs=np.zeros([nrep,n_epoch])
nm_all_costs=np.zeros([nrep,n_epoch])
for rep_ind in range(nrep):
    nm_all_accs[rep_ind]=multi_locs[rep_ind][0]
    nm_all_costs[rep_ind]=multi_locs[rep_ind][1]
    
    
np.save('sim_data/Fig_1_100_fashion_sp_cst',sp_all_costs)
np.save('sim_data/Fig_1_100_fashion_sp_acc',sp_all_accs)
np.save('sim_data/Fig_1_100_fashion_nm_cst',nm_all_costs)
np.save('sim_data/Fig_1_100_fashion_nm_acc',nm_all_accs)

#%%###############################################################################
###############################################################################
###############################################################################
# Digits with 300 hidden neurons


pool = mp.Pool(mp.cpu_count()) # Multiprocessing         
multi_locs=[pool.apply(sp_func, args=([300],n_epoch,dig_training,dig_test)) for dummy_ind in range(nrep)] # Check all_funcs for defaults
pool.close()
sp_all_accs=np.zeros([nrep,n_epoch])
sp_all_costs=np.zeros([nrep,n_epoch])
for rep_ind in range(nrep):
    sp_all_accs[rep_ind]=multi_locs[rep_ind][0]
    sp_all_costs[rep_ind]=multi_locs[rep_ind][1]


pool = mp.Pool(mp.cpu_count()) # Multiprocessing         
multi_locs=[pool.apply(nm_func, args=([300],n_epoch,dig_training,dig_test)) for dummy_ind in range(nrep)] # Check all_funcs for defaults
pool.close()
nm_all_accs=np.zeros([nrep,n_epoch])
nm_all_costs=np.zeros([nrep,n_epoch])
for rep_ind in range(nrep):
    nm_all_accs[rep_ind]=multi_locs[rep_ind][0]
    nm_all_costs[rep_ind]=multi_locs[rep_ind][1]
    
    
np.save('sim_data/Fig_1_300_digits_sp_cst',sp_all_costs)
np.save('sim_data/Fig_1_300_digits_sp_acc',sp_all_accs)
np.save('sim_data/Fig_1_300_digits_nm_cst',nm_all_costs)
np.save('sim_data/Fig_1_300_digits_nm_acc',nm_all_accs)


#%%
###############################################################################
###############################################################################
###############################################################################
# Fashion with with 300 hidden neurons
pool = mp.Pool(mp.cpu_count()) # Multiprocessing         
multi_locs=[pool.apply(sp_func, args=([300],n_epoch,fash_training,fash_test)) for dummy_ind in range(nrep)] # Check all_funcs for defaults
pool.close()
sp_all_accs=np.zeros([nrep,n_epoch])
sp_all_costs=np.zeros([nrep,n_epoch])
for rep_ind in range(nrep):
    sp_all_accs[rep_ind]=multi_locs[rep_ind][0]
    sp_all_costs[rep_ind]=multi_locs[rep_ind][1]


pool = mp.Pool(mp.cpu_count()) # Multiprocessing         
multi_locs=[pool.apply(nm_func, args=([300],n_epoch,fash_training,fash_test)) for dummy_ind in range(nrep)] # Check all_funcs for defaults
pool.close()
nm_all_accs=np.zeros([nrep,n_epoch])
nm_all_costs=np.zeros([nrep,n_epoch])
for rep_ind in range(nrep):
    nm_all_accs[rep_ind]=multi_locs[rep_ind][0]
    nm_all_costs[rep_ind]=multi_locs[rep_ind][1]
    
    
np.save('sim_data/Fig_1_300_fashion_sp_cst',sp_all_costs)
np.save('sim_data/Fig_1_300_fashion_sp_acc',sp_all_accs)
np.save('sim_data/Fig_1_300_fashion_nm_cst',nm_all_costs)
np.save('sim_data/Fig_1_300_fashion_nm_acc',nm_all_accs)
```
