## Supplementary material for "Dendritic normalisation improves learning in sparsely connected artificial neural networks": Code and data.: Fig_1_plot.py

```
# -*- coding: utf-8 -*-
"""
Created on Tue Nov 19 14:45:18 2019

@author: Alex Bird
"""
#%%
# Panel D_1 30 neurons, digits
sp_cst=np.load('sim_data/Fig_1_30_digits_sp_cst.npy')
sp_acc=np.load('sim_data/Fig_1_30_digits_sp_acc.npy')/10000
nm_cst=np.load('sim_data/Fig_1_30_digits_nm_cst.npy')
nm_acc=np.load('sim_data/Fig_1_30_digits_nm_acc.npy')/10000

n_rep=sp_cst.shape[0]
n_epoch=sp_cst.shape[1]
sp_cst_mn=np.zeros(n_epoch)
sp_cst_sd=np.zeros(n_epoch)
sp_acc_mn=np.zeros(n_epoch)
sp_acc_sd=np.zeros(n_epoch)
nm_cst_mn=np.zeros(n_epoch)
nm_cst_sd=np.zeros(n_epoch)
nm_acc_mn=np.zeros(n_epoch)
nm_acc_sd=np.zeros(n_epoch)

for ep_ind in range(n_epoch):
    vec_sp_cst=sp_cst[:,ep_ind]
    sp_cst_mn[ep_ind]=np.mean(vec_sp_cst)
    sp_cst_sd[ep_ind]=(n_rep/(n_rep-1)*np.var(vec_sp_cst))**0.5
    vec_sp_acc=sp_acc[:,ep_ind]
    sp_acc_mn[ep_ind]=np.mean(vec_sp_acc)
    sp_acc_sd[ep_ind]=(n_rep/(n_rep-1)*np.var(vec_sp_acc))**0.5
    vec_nm_cst=nm_cst[:,ep_ind]
    nm_cst_mn[ep_ind]=np.mean(vec_nm_cst)
    nm_cst_sd[ep_ind]=(n_rep/(n_rep-1)*np.var(vec_nm_cst))**0.5
    vec_nm_acc=nm_acc[:,ep_ind]
    nm_acc_mn[ep_ind]=np.mean(vec_nm_acc)
    nm_acc_sd[ep_ind]=(n_rep/(n_rep-1)*np.var(vec_nm_acc))**0.5


sns.set(style="white", palette="muted", color_codes=True)
f, axes = plt.subplots(2, 1, figsize=(1.1811,2.36), sharex=True , sharey=False)
sns.despine      
axes[0].plot(range(n_epoch),sp_cst_mn,'tab:blue',range(n_epoch),nm_cst_mn,'tab:orange',linewidth=0.5)
axes[0].fill_between(range(n_epoch),sp_cst_mn-sp_cst_sd,sp_cst_mn+sp_cst_sd,facecolor='tab:blue',color=None,alpha=0.2)
axes[0].fill_between(range(n_epoch),nm_cst_mn-nm_cst_sd,nm_cst_mn+nm_cst_sd,facecolor='tab:orange',color=None,alpha=0.2)      
axes[0].spines['top'].set_visible(False)
axes[0].spines['right'].set_visible(False)
axes[0].spines['bottom'].set_edgecolor('black')
axes[0].spines['bottom'].set_facecolor('black')
axes[0].spines['bottom'].set_linewidth(0.5)
axes[0].spines['left'].set_edgecolor('black')
axes[0].spines['left'].set_facecolor('black')
axes[0].spines['left'].set_linewidth(0.5)
axes[0].set_xticks([0,50,100])
axes[0].set_yticks([0.2,0.5,0.8],[0.2,0.5,0.8])
axes[0].tick_params(direction='out', length=2, width=1 , labelsize=6 , pad=1)
 

axes[1].plot(range(n_epoch),sp_acc_mn,'tab:blue',range(n_epoch),nm_acc_mn,'tab:orange',linewidth=0.5)
axes[1].fill_between(range(n_epoch),sp_acc_mn-sp_acc_sd,sp_acc_mn+sp_acc_sd,facecolor='tab:blue',color=None,alpha=0.2)
axes[1].fill_between(range(n_epoch),nm_acc_mn-nm_acc_sd,nm_acc_mn+nm_acc_sd,facecolor='tab:orange',color=None,alpha=0.2)      
axes[1].spines['top'].set_visible(False)
axes[1].spines['right'].set_visible(False)
axes[1].spines['bottom'].set_edgecolor('black')
axes[1].spines['bottom'].set_facecolor('black')
axes[1].spines['bottom'].set_linewidth(0.5)
axes[1].spines['left'].set_edgecolor('black')
axes[1].spines['left'].set_facecolor('black')
axes[1].spines['left'].set_linewidth(0.5)
axes[1].set_yticks=[7000,8000,9000]
axes[1].tick_params(direction='out', length=2, width=1, labelsize=6 , pad=1)  

plt.savefig('fig/Fig_1_d_1.pdf', format='pdf',bbox_inches='tight',pad_inches=0.25)

#%%
# Panel D_2 30 neurons, fashion
sp_cst=np.load('sim_data/Fig_1_30_fashion_sp_cst.npy')
sp_acc=np.load('sim_data/Fig_1_30_fashion_sp_acc.npy')/10000
nm_cst=np.load('sim_data/Fig_1_30_fashion_nm_cst.npy')
nm_acc=np.load('sim_data/Fig_1_30_fashion_nm_acc.npy')/10000

n_rep=sp_cst.shape[0]
n_epoch=sp_cst.shape[1]
sp_cst_mn=np.zeros(n_epoch)
sp_cst_sd=np.zeros(n_epoch)
sp_acc_mn=np.zeros(n_epoch)
sp_acc_sd=np.zeros(n_epoch)
nm_cst_mn=np.zeros(n_epoch)
nm_cst_sd=np.zeros(n_epoch)
nm_acc_mn=np.zeros(n_epoch)
nm_acc_sd=np.zeros(n_epoch)

for ep_ind in range(n_epoch):
    vec_sp_cst=sp_cst[:,ep_ind]
    sp_cst_mn[ep_ind]=np.mean(vec_sp_cst)
    sp_cst_sd[ep_ind]=(n_rep/(n_rep-1)*np.var(vec_sp_cst))**0.5
    vec_sp_acc=sp_acc[:,ep_ind]
    sp_acc_mn[ep_ind]=np.mean(vec_sp_acc)
    sp_acc_sd[ep_ind]=(n_rep/(n_rep-1)*np.var(vec_sp_acc))**0.5
    vec_nm_cst=nm_cst[:,ep_ind]
    nm_cst_mn[ep_ind]=np.mean(vec_nm_cst)
    nm_cst_sd[ep_ind]=(n_rep/(n_rep-1)*np.var(vec_nm_cst))**0.5
    vec_nm_acc=nm_acc[:,ep_ind]
    nm_acc_mn[ep_ind]=np.mean(vec_nm_acc)
    nm_acc_sd[ep_ind]=(n_rep/(n_rep-1)*np.var(vec_nm_acc))**0.5


sns.set(style="white", palette="muted", color_codes=True)
f, axes = plt.subplots(2, 1, figsize=(1.1811,2.36), sharex=True , sharey=False)
sns.despine      
axes[0].plot(range(n_epoch),sp_cst_mn,'tab:blue',range(n_epoch),nm_cst_mn,'tab:orange',linewidth=0.5)
axes[0].fill_between(range(n_epoch),sp_cst_mn-sp_cst_sd,sp_cst_mn+sp_cst_sd,facecolor='tab:blue',color=None,alpha=0.2)
axes[0].fill_between(range(n_epoch),nm_cst_mn-nm_cst_sd,nm_cst_mn+nm_cst_sd,facecolor='tab:orange',color=None,alpha=0.2)      
axes[0].spines['top'].set_visible(False)
axes[0].spines['right'].set_visible(False)
axes[0].spines['bottom'].set_edgecolor('black')
axes[0].spines['bottom'].set_facecolor('black')
axes[0].spines['bottom'].set_linewidth(0.5)
axes[0].spines['left'].set_edgecolor('black')
axes[0].spines['left'].set_facecolor('black')
axes[0].spines['left'].set_linewidth(0.5)
axes[0].set_xticks([0,50,100])
axes[0].set_yticks([0.2,0.5,0.8],[0.2,0.5,0.8])
axes[0].tick_params(direction='out', length=2, width=1 , labelsize=6 , pad=1)
 

axes[1].plot(range(n_epoch),sp_acc_mn,'tab:blue',range(n_epoch),nm_acc_mn,'tab:orange',linewidth=0.5)
axes[1].fill_between(range(n_epoch),sp_acc_mn-sp_acc_sd,sp_acc_mn+sp_acc_sd,facecolor='tab:blue',color=None,alpha=0.2)
axes[1].fill_between(range(n_epoch),nm_acc_mn-nm_acc_sd,nm_acc_mn+nm_acc_sd,facecolor='tab:orange',color=None,alpha=0.2)      
axes[1].spines['top'].set_visible(False)
axes[1].spines['right'].set_visible(False)
axes[1].spines['bottom'].set_edgecolor('black')
axes[1].spines['bottom'].set_facecolor('black')
axes[1].spines['bottom'].set_linewidth(0.5)
axes[1].spines['left'].set_edgecolor('black')
axes[1].spines['left'].set_facecolor('black')
axes[1].spines['left'].set_linewidth(0.5)
axes[1].set_yticks=[7000,8000,9000]
axes[1].tick_params(direction='out', length=2, width=1, labelsize=6 , pad=1)  

plt.savefig('fig/Fig_1_d_2.pdf', format='pdf',bbox_inches='tight',pad_inches=0.25)


#%%
# Panel D_3 100 neurons, digits
sp_cst=np.load('sim_data/Fig_1_100_digits_sp_cst.npy')
sp_acc=np.load('sim_data/Fig_1_100_digits_sp_acc.npy')/10000
nm_cst=np.load('sim_data/Fig_1_100_digits_nm_cst.npy')
nm_acc=np.load('sim_data/Fig_1_100_digits_nm_acc.npy')/10000

n_rep=sp_cst.shape[0]
n_epoch=sp_cst.shape[1]
sp_cst_mn=np.zeros(n_epoch)
sp_cst_sd=np.zeros(n_epoch)
sp_acc_mn=np.zeros(n_epoch)
sp_acc_sd=np.zeros(n_epoch)
nm_cst_mn=np.zeros(n_epoch)
nm_cst_sd=np.zeros(n_epoch)
nm_acc_mn=np.zeros(n_epoch)
nm_acc_sd=np.zeros(n_epoch)

for ep_ind in range(n_epoch):
    vec_sp_cst=sp_cst[:,ep_ind]
    sp_cst_mn[ep_ind]=np.mean(vec_sp_cst)
    sp_cst_sd[ep_ind]=(n_rep/(n_rep-1)*np.var(vec_sp_cst))**0.5
    vec_sp_acc=sp_acc[:,ep_ind]
    sp_acc_mn[ep_ind]=np.mean(vec_sp_acc)
    sp_acc_sd[ep_ind]=(n_rep/(n_rep-1)*np.var(vec_sp_acc))**0.5
    vec_nm_cst=nm_cst[:,ep_ind]
    nm_cst_mn[ep_ind]=np.mean(vec_nm_cst)
    nm_cst_sd[ep_ind]=(n_rep/(n_rep-1)*np.var(vec_nm_cst))**0.5
    vec_nm_acc=nm_acc[:,ep_ind]
    nm_acc_mn[ep_ind]=np.mean(vec_nm_acc)
    nm_acc_sd[ep_ind]=(n_rep/(n_rep-1)*np.var(vec_nm_acc))**0.5


sns.set(style="white", palette="muted", color_codes=True)
f, axes = plt.subplots(2, 1, figsize=(1.1811,2.36), sharex=True , sharey=False)
sns.despine      
axes[0].plot(range(n_epoch),sp_cst_mn,'tab:blue',range(n_epoch),nm_cst_mn,'tab:orange',linewidth=0.5)
axes[0].fill_between(range(n_epoch),sp_cst_mn-sp_cst_sd,sp_cst_mn+sp_cst_sd,facecolor='tab:blue',color=None,alpha=0.2)
axes[0].fill_between(range(n_epoch),nm_cst_mn-nm_cst_sd,nm_cst_mn+nm_cst_sd,facecolor='tab:orange',color=None,alpha=0.2)      
axes[0].spines['top'].set_visible(False)
axes[0].spines['right'].set_visible(False)
axes[0].spines['bottom'].set_edgecolor('black')
axes[0].spines['bottom'].set_facecolor('black')
axes[0].spines['bottom'].set_linewidth(0.5)
axes[0].spines['left'].set_edgecolor('black')
axes[0].spines['left'].set_facecolor('black')
axes[0].spines['left'].set_linewidth(0.5)
axes[0].set_xticks([0,50,100])
axes[0].set_yticks([0.2,0.5,0.8],[0.2,0.5,0.8])
axes[0].tick_params(direction='out', length=2, width=1 , labelsize=6 , pad=1)
 

axes[1].plot(range(n_epoch),sp_acc_mn,'tab:blue',range(n_epoch),nm_acc_mn,'tab:orange',linewidth=0.5)
axes[1].fill_between(range(n_epoch),sp_acc_mn-sp_acc_sd,sp_acc_mn+sp_acc_sd,facecolor='tab:blue',color=None,alpha=0.2)
axes[1].fill_between(range(n_epoch),nm_acc_mn-nm_acc_sd,nm_acc_mn+nm_acc_sd,facecolor='tab:orange',color=None,alpha=0.2)      
axes[1].spines['top'].set_visible(False)
axes[1].spines['right'].set_visible(False)
axes[1].spines['bottom'].set_edgecolor('black')
axes[1].spines['bottom'].set_facecolor('black')
axes[1].spines['bottom'].set_linewidth(0.5)
axes[1].spines['left'].set_edgecolor('black')
axes[1].spines['left'].set_facecolor('black')
axes[1].spines['left'].set_linewidth(0.5)
axes[1].set_yticks=[7000,8000,9000]
axes[1].tick_params(direction='out', length=2, width=1, labelsize=6 , pad=1)  

plt.savefig('fig/Fig_1_d_3.pdf', format='pdf',bbox_inches='tight',pad_inches=0.25)

#%%
# Panel D_4 100 neurons, fashion
sp_cst=np.load('sim_data/Fig_1_100_fashion_sp_cst.npy')
sp_acc=np.load('sim_data/Fig_1_100_fashion_sp_acc.npy')/10000
nm_cst=np.load('sim_data/Fig_1_100_fashion_nm_cst.npy')
nm_acc=np.load('sim_data/Fig_1_100_fashion_nm_acc.npy')/10000

n_rep=sp_cst.shape[0]
n_epoch=sp_cst.shape[1]
sp_cst_mn=np.zeros(n_epoch)
sp_cst_sd=np.zeros(n_epoch)
sp_acc_mn=np.zeros(n_epoch)
sp_acc_sd=np.zeros(n_epoch)
nm_cst_mn=np.zeros(n_epoch)
nm_cst_sd=np.zeros(n_epoch)
nm_acc_mn=np.zeros(n_epoch)
nm_acc_sd=np.zeros(n_epoch)

for ep_ind in range(n_epoch):
    vec_sp_cst=sp_cst[:,ep_ind]
    sp_cst_mn[ep_ind]=np.mean(vec_sp_cst)
    sp_cst_sd[ep_ind]=(n_rep/(n_rep-1)*np.var(vec_sp_cst))**0.5
    vec_sp_acc=sp_acc[:,ep_ind]
    sp_acc_mn[ep_ind]=np.mean(vec_sp_acc)
    sp_acc_sd[ep_ind]=(n_rep/(n_rep-1)*np.var(vec_sp_acc))**0.5
    vec_nm_cst=nm_cst[:,ep_ind]
    nm_cst_mn[ep_ind]=np.mean(vec_nm_cst)
    nm_cst_sd[ep_ind]=(n_rep/(n_rep-1)*np.var(vec_nm_cst))**0.5
    vec_nm_acc=nm_acc[:,ep_ind]
    nm_acc_mn[ep_ind]=np.mean(vec_nm_acc)
    nm_acc_sd[ep_ind]=(n_rep/(n_rep-1)*np.var(vec_nm_acc))**0.5


sns.set(style="white", palette="muted", color_codes=True)
f, axes = plt.subplots(2, 1, figsize=(1.1811,2.36), sharex=True , sharey=False)
sns.despine      
axes[0].plot(range(n_epoch),sp_cst_mn,'tab:blue',range(n_epoch),nm_cst_mn,'tab:orange',linewidth=0.5)
axes[0].fill_between(range(n_epoch),sp_cst_mn-sp_cst_sd,sp_cst_mn+sp_cst_sd,facecolor='tab:blue',color=None,alpha=0.2)
axes[0].fill_between(range(n_epoch),nm_cst_mn-nm_cst_sd,nm_cst_mn+nm_cst_sd,facecolor='tab:orange',color=None,alpha=0.2)      
axes[0].spines['top'].set_visible(False)
axes[0].spines['right'].set_visible(False)
axes[0].spines['bottom'].set_edgecolor('black')
axes[0].spines['bottom'].set_facecolor('black')
axes[0].spines['bottom'].set_linewidth(0.5)
axes[0].spines['left'].set_edgecolor('black')
axes[0].spines['left'].set_facecolor('black')
axes[0].spines['left'].set_linewidth(0.5)
axes[0].set_xticks([0,50,100])
axes[0].set_yticks([0.2,0.5,0.8],[0.2,0.5,0.8])
axes[0].tick_params(direction='out', length=2, width=1 , labelsize=6 , pad=1)
 

axes[1].plot(range(n_epoch),sp_acc_mn,'tab:blue',range(n_epoch),nm_acc_mn,'tab:orange',linewidth=0.5)
axes[1].fill_between(range(n_epoch),sp_acc_mn-sp_acc_sd,sp_acc_mn+sp_acc_sd,facecolor='tab:blue',color=None,alpha=0.2)
axes[1].fill_between(range(n_epoch),nm_acc_mn-nm_acc_sd,nm_acc_mn+nm_acc_sd,facecolor='tab:orange',color=None,alpha=0.2)      
axes[1].spines['top'].set_visible(False)
axes[1].spines['right'].set_visible(False)
axes[1].spines['bottom'].set_edgecolor('black')
axes[1].spines['bottom'].set_facecolor('black')
axes[1].spines['bottom'].set_linewidth(0.5)
axes[1].spines['left'].set_edgecolor('black')
axes[1].spines['left'].set_facecolor('black')
axes[1].spines['left'].set_linewidth(0.5)
axes[1].set_yticks=[7000,8000,9000]
axes[1].tick_params(direction='out', length=2, width=1, labelsize=6 , pad=1)  

plt.savefig('fig/Fig_1_d_4.pdf', format='pdf',bbox_inches='tight',pad_inches=0.25)


#%%
# Panel D_5 300 neurons, digits
sp_cst=np.load('sim_data/Fig_1_300_digits_sp_cst.npy')
sp_acc=np.load('sim_data/Fig_1_300_digits_sp_acc.npy')/10000
nm_cst=np.load('sim_data/Fig_1_300_digits_nm_cst.npy')
nm_acc=np.load('sim_data/Fig_1_300_digits_nm_acc.npy')/10000

n_rep=sp_cst.shape[0]
n_epoch=sp_cst.shape[1]
sp_cst_mn=np.zeros(n_epoch)
sp_cst_sd=np.zeros(n_epoch)
sp_acc_mn=np.zeros(n_epoch)
sp_acc_sd=np.zeros(n_epoch)
nm_cst_mn=np.zeros(n_epoch)
nm_cst_sd=np.zeros(n_epoch)
nm_acc_mn=np.zeros(n_epoch)
nm_acc_sd=np.zeros(n_epoch)

for ep_ind in range(n_epoch):
    vec_sp_cst=sp_cst[:,ep_ind]
    sp_cst_mn[ep_ind]=np.mean(vec_sp_cst)
    sp_cst_sd[ep_ind]=(n_rep/(n_rep-1)*np.var(vec_sp_cst))**0.5
    vec_sp_acc=sp_acc[:,ep_ind]
    sp_acc_mn[ep_ind]=np.mean(vec_sp_acc)
    sp_acc_sd[ep_ind]=(n_rep/(n_rep-1)*np.var(vec_sp_acc))**0.5
    vec_nm_cst=nm_cst[:,ep_ind]
    nm_cst_mn[ep_ind]=np.mean(vec_nm_cst)
    nm_cst_sd[ep_ind]=(n_rep/(n_rep-1)*np.var(vec_nm_cst))**0.5
    vec_nm_acc=nm_acc[:,ep_ind]
    nm_acc_mn[ep_ind]=np.mean(vec_nm_acc)
    nm_acc_sd[ep_ind]=(n_rep/(n_rep-1)*np.var(vec_nm_acc))**0.5


sns.set(style="white", palette="muted", color_codes=True)
f, axes = plt.subplots(2, 1, figsize=(1.1811,2.36), sharex=True , sharey=False)
sns.despine      
axes[0].plot(range(n_epoch),sp_cst_mn,'tab:blue',range(n_epoch),nm_cst_mn,'tab:orange',linewidth=0.5)
axes[0].fill_between(range(n_epoch),sp_cst_mn-sp_cst_sd,sp_cst_mn+sp_cst_sd,facecolor='tab:blue',color=None,alpha=0.2)
axes[0].fill_between(range(n_epoch),nm_cst_mn-nm_cst_sd,nm_cst_mn+nm_cst_sd,facecolor='tab:orange',color=None,alpha=0.2)      
axes[0].spines['top'].set_visible(False)
axes[0].spines['right'].set_visible(False)
axes[0].spines['bottom'].set_edgecolor('black')
axes[0].spines['bottom'].set_facecolor('black')
axes[0].spines['bottom'].set_linewidth(0.5)
axes[0].spines['left'].set_edgecolor('black')
axes[0].spines['left'].set_facecolor('black')
axes[0].spines['left'].set_linewidth(0.5)
axes[0].set_xticks([0,50,100])
axes[0].set_yticks([0.2,0.5,0.8],[0.2,0.5,0.8])
axes[0].tick_params(direction='out', length=2, width=1 , labelsize=6 , pad=1)
 

axes[1].plot(range(n_epoch),sp_acc_mn,'tab:blue',range(n_epoch),nm_acc_mn,'tab:orange',linewidth=0.5)
axes[1].fill_between(range(n_epoch),sp_acc_mn-sp_acc_sd,sp_acc_mn+sp_acc_sd,facecolor='tab:blue',color=None,alpha=0.2)
axes[1].fill_between(range(n_epoch),nm_acc_mn-nm_acc_sd,nm_acc_mn+nm_acc_sd,facecolor='tab:orange',color=None,alpha=0.2)      
axes[1].spines['top'].set_visible(False)
axes[1].spines['right'].set_visible(False)
axes[1].spines['bottom'].set_edgecolor('black')
axes[1].spines['bottom'].set_facecolor('black')
axes[1].spines['bottom'].set_linewidth(0.5)
axes[1].spines['left'].set_edgecolor('black')
axes[1].spines['left'].set_facecolor('black')
axes[1].spines['left'].set_linewidth(0.5)
axes[1].set_yticks=[7000,8000,9000]
axes[1].tick_params(direction='out', length=2, width=1, labelsize=6 , pad=1)  

plt.savefig('fig/Fig_1_d_5.pdf', format='pdf',bbox_inches='tight',pad_inches=0.25)

#%%
# Panel D_6 300 neurons, fashion
sp_cst=np.load('sim_data/Fig_1_300_fashion_sp_cst.npy')
sp_acc=np.load('sim_data/Fig_1_300_fashion_sp_acc.npy')/10000
nm_cst=np.load('sim_data/Fig_1_300_fashion_nm_cst.npy')
nm_acc=np.load('sim_data/Fig_1_300_fashion_nm_acc.npy')/10000

n_rep=sp_cst.shape[0]
n_epoch=sp_cst.shape[1]
sp_cst_mn=np.zeros(n_epoch)
sp_cst_sd=np.zeros(n_epoch)
sp_acc_mn=np.zeros(n_epoch)
sp_acc_sd=np.zeros(n_epoch)
nm_cst_mn=np.zeros(n_epoch)
nm_cst_sd=np.zeros(n_epoch)
nm_acc_mn=np.zeros(n_epoch)
nm_acc_sd=np.zeros(n_epoch)

for ep_ind in range(n_epoch):
    vec_sp_cst=sp_cst[:,ep_ind]
    sp_cst_mn[ep_ind]=np.mean(vec_sp_cst)
    sp_cst_sd[ep_ind]=(n_rep/(n_rep-1)*np.var(vec_sp_cst))**0.5
    vec_sp_acc=sp_acc[:,ep_ind]
    sp_acc_mn[ep_ind]=np.mean(vec_sp_acc)
    sp_acc_sd[ep_ind]=(n_rep/(n_rep-1)*np.var(vec_sp_acc))**0.5
    vec_nm_cst=nm_cst[:,ep_ind]
    nm_cst_mn[ep_ind]=np.mean(vec_nm_cst)
    nm_cst_sd[ep_ind]=(n_rep/(n_rep-1)*np.var(vec_nm_cst))**0.5
    vec_nm_acc=nm_acc[:,ep_ind]
    nm_acc_mn[ep_ind]=np.mean(vec_nm_acc)
    nm_acc_sd[ep_ind]=(n_rep/(n_rep-1)*np.var(vec_nm_acc))**0.5


sns.set(style="white", palette="muted", color_codes=True)
f, axes = plt.subplots(2, 1, figsize=(1.1811,2.36), sharex=True , sharey=False)
sns.despine      
axes[0].plot(range(n_epoch),sp_cst_mn,'tab:blue',range(n_epoch),nm_cst_mn,'tab:orange',linewidth=0.5)
axes[0].fill_between(range(n_epoch),sp_cst_mn-sp_cst_sd,sp_cst_mn+sp_cst_sd,facecolor='tab:blue',color=None,alpha=0.2)
axes[0].fill_between(range(n_epoch),nm_cst_mn-nm_cst_sd,nm_cst_mn+nm_cst_sd,facecolor='tab:orange',color=None,alpha=0.2)      
axes[0].spines['top'].set_visible(False)
axes[0].spines['right'].set_visible(False)
axes[0].spines['bottom'].set_edgecolor('black')
axes[0].spines['bottom'].set_facecolor('black')
axes[0].spines['bottom'].set_linewidth(0.5)
axes[0].spines['left'].set_edgecolor('black')
axes[0].spines['left'].set_facecolor('black')
axes[0].spines['left'].set_linewidth(0.5)
axes[0].set_xticks([0,50,100])
axes[0].set_yticks([0.2,0.5,0.8],[0.2,0.5,0.8])
axes[0].tick_params(direction='out', length=2, width=1 , labelsize=6 , pad=1)
 

axes[1].plot(range(n_epoch),sp_acc_mn,'tab:blue',range(n_epoch),nm_acc_mn,'tab:orange',linewidth=0.5)
axes[1].fill_between(range(n_epoch),sp_acc_mn-sp_acc_sd,sp_acc_mn+sp_acc_sd,facecolor='tab:blue',color=None,alpha=0.2)
axes[1].fill_between(range(n_epoch),nm_acc_mn-nm_acc_sd,nm_acc_mn+nm_acc_sd,facecolor='tab:orange',color=None,alpha=0.2)      
axes[1].spines['top'].set_visible(False)
axes[1].spines['right'].set_visible(False)
axes[1].spines['bottom'].set_edgecolor('black')
axes[1].spines['bottom'].set_facecolor('black')
axes[1].spines['bottom'].set_linewidth(0.5)
axes[1].spines['left'].set_edgecolor('black')
axes[1].spines['left'].set_facecolor('black')
axes[1].spines['left'].set_linewidth(0.5)
axes[1].set_yticks=[7000,8000,9000]
axes[1].tick_params(direction='out', length=2, width=1, labelsize=6 , pad=1)  

plt.savefig('fig/Fig_1_d_6.pdf', format='pdf',bbox_inches='tight',pad_inches=0.25)
```
