## Supplementary material for "Dendritic normalisation improves learning in sparsely connected artificial neural networks": Code and data.: Fig_2_data_gen.py

```
# -*- coding: utf-8 -*-
"""
Created on Mon Nov 18 18:06:23 2019

@author: Alex Bird
"""

# Digits with 100 hidden neurons


n_epoch=100
n_rep=25

sp_neff_1=[] # Number of contacts leaving each pixel
nm_neff_1=[]
sp_neff_10=[] 
nm_neff_10=[]
sp_neff_100=[] 
nm_neff_100=[]

sp_naff_1=[] # Number of contacts arriving at each hidden neuron 
nm_naff_1=[]
sp_naff_5=[] 
nm_naff_5=[]
sp_naff_10=[] 
nm_naff_10=[]
sp_naff_25=[] 
nm_naff_25=[]
sp_naff_100=[] 
nm_naff_100=[]

sp_wght_1=[] # Weights 
nm_wght_1=[]
sp_wght_5=[] 
nm_wght_5=[]
sp_wght_10=[] 
nm_wght_10=[]
sp_wght_25=[] 
nm_wght_25=[]
sp_wght_100=[] 
nm_wght_100=[]

sp_mnact_1=[] # Mean activation over test set
nm_mnact_1=[]
sp_mnact_5=[] 
nm_mnact_5=[]
sp_mnact_10=[] 
nm_mnact_10=[]
sp_mnact_25=[] 
nm_mnact_25=[]
sp_mnact_100=[] 
nm_mnact_100=[]


for ind in range(n_rep):
    net_sp=network_sp.Network([784,100,10],0.2,0.15)
    net_nm=network_nm.Network([784,100,10],0.2,0.15)
    
    ##########################################################################
    ##########################################################################
    ##########################################################################
    # 1 epoch
    isp_acc,isp_cst=net_sp.SGD(dig_training,1,10,0.5,evaluation_data=dig_test)
    inm_acc,inm_cst=net_nm.SGD(dig_training,1,10,0.5,evaluation_data=dig_test)
    
    i_sp_wght=net_sp.weights[0]
    i_nm_wght=net_nm.weights[0]
    
    # Number of contacts leaving each pixel    
    nnz_sp=np.count_nonzero(i_sp_wght,0)
    nnz_sp.reshape([28,28])
    sp_neff_1.append(nnz_sp)
    
    nnz_nm=np.count_nonzero(i_nm_wght,0)
    nnz_nm.reshape([28,28])
    nm_neff_1.append(nnz_nm)
      
    # Number of contacts arriving at each hidden neuron 
    nnz_sp=np.count_nonzero(i_sp_wght,1)
    sp_naff_1.append(nnz_sp)
    
    nnz_nm=np.count_nonzero(i_nm_wght,1)
    nm_naff_1.append(nnz_nm)
    
    # Weights 
    wght_sp=i_sp_wght[np.nonzero(i_sp_wght)]
    sp_wght_1.append(wght_sp)
    
    wght_nm=i_nm_wght[np.nonzero(i_nm_wght)]
    nm_wght_1.append(wght_nm)
    
    # Mean activations over test set
    a_sp=input_to_hidden_layer(net_sp, dig_test)
    sp_mnact_1.append(a_sp)
    
    a_nm=input_to_hidden_layer(net_nm, dig_test)
    nm_mnact_1.append(a_nm)
    
    ##########################################################################
    ##########################################################################
    ##########################################################################
    # 5 epochs
    isp_acc,isp_cst=net_sp.SGD(dig_training,4,10,0.5,evaluation_data=dig_test)
    inm_acc,inm_cst=net_nm.SGD(dig_training,4,10,0.5,evaluation_data=dig_test)
    
    i_sp_wght=net_sp.weights[0]
    i_nm_wght=net_nm.weights[0]    
      
    # Number of contacts arriving at each hidden neuron 
    nnz_sp=np.count_nonzero(i_sp_wght,1)
    sp_naff_5.append(nnz_sp)
    
    nnz_nm=np.count_nonzero(i_nm_wght,1)
    nm_naff_5.append(nnz_nm)
    
    # Weights 
    wght_sp=i_sp_wght[np.nonzero(i_sp_wght)]
    sp_wght_5.append(wght_sp)
    
    wght_nm=i_nm_wght[np.nonzero(i_nm_wght)]
    nm_wght_5.append(wght_nm)
    
    # Mean activations over test set
    a_sp=input_to_hidden_layer(net_sp, dig_test)
    sp_mnact_5.append(a_sp)
    
    a_nm=input_to_hidden_layer(net_nm, dig_test)
    nm_mnact_5.append(a_nm)
    
    ##########################################################################
    ##########################################################################
    ##########################################################################
    # 10 epochs
    
    isp_acc,isp_cst=net_sp.SGD(dig_training,5,10,0.5,evaluation_data=dig_test)
    inm_acc,inm_cst=net_nm.SGD(dig_training,5,10,0.5,evaluation_data=dig_test)
    
    i_sp_wght=net_sp.weights[0]
    i_nm_wght=net_nm.weights[0]
    
    # Number of contacts leaving each pixel    
    nnz_sp=np.count_nonzero(i_sp_wght,0)
    nnz_sp.reshape([28,28])
    sp_neff_10.append(nnz_sp)
    
    nnz_nm=np.count_nonzero(i_nm_wght,0)
    nnz_nm.reshape([28,28])
    nm_neff_10.append(nnz_nm)
      
    # Number of contacts arriving at each hidden neuron 
    nnz_sp=np.count_nonzero(i_sp_wght,1)
    sp_naff_10.append(nnz_sp)
    
    nnz_nm=np.count_nonzero(i_nm_wght,1)
    nm_naff_10.append(nnz_nm)
    
    # Weights 
    wght_sp=i_sp_wght[np.nonzero(i_sp_wght)]
    sp_wght_10.append(wght_sp)
    
    wght_nm=i_nm_wght[np.nonzero(i_nm_wght)]
    nm_wght_10.append(wght_nm)
    
    # Mean activations over test set
    a_sp=input_to_hidden_layer(net_sp, dig_test)
    sp_mnact_10.append(a_sp)
    
    a_nm=input_to_hidden_layer(net_nm, dig_test)
    nm_mnact_10.append(a_nm)
    
    ##########################################################################
    ##########################################################################
    ##########################################################################
    # 25 epochs
    isp_acc,isp_cst=net_sp.SGD(dig_training,15,10,0.5,evaluation_data=dig_test)
    inm_acc,inm_cst=net_nm.SGD(dig_training,15,10,0.5,evaluation_data=dig_test)
    
    i_sp_wght=net_sp.weights[0]
    i_nm_wght=net_nm.weights[0]    
      
    # Number of contacts arriving at each hidden neuron 
    nnz_sp=np.count_nonzero(i_sp_wght,1)
    sp_naff_25.append(nnz_sp)
    
    nnz_nm=np.count_nonzero(i_nm_wght,1)
    nm_naff_25.append(nnz_nm)
    
    # Weights 
    wght_sp=i_sp_wght[np.nonzero(i_sp_wght)]
    sp_wght_25.append(wght_sp)
    
    wght_nm=i_nm_wght[np.nonzero(i_nm_wght)]
    nm_wght_25.append(wght_nm)
    
    # Mean activations over test set
    a_sp=input_to_hidden_layer(net_sp, dig_test)
    sp_mnact_25.append(a_sp)
    
    a_nm=input_to_hidden_layer(net_nm, dig_test)
    nm_mnact_25.append(a_nm)
  
    ##########################################################################
    ##########################################################################
    ##########################################################################
    # 100 epochs
    
    isp_acc,isp_cst=net_sp.SGD(dig_training,75,10,0.5,evaluation_data=dig_test)
    inm_acc,inm_cst=net_nm.SGD(dig_training,75,10,0.5,evaluation_data=dig_test)
    
    i_sp_wght=net_sp.weights[0]
    i_nm_wght=net_nm.weights[0]
    
    # Number of contacts leaving each pixel    
    nnz_sp=np.count_nonzero(i_sp_wght,0)
    nnz_sp.reshape([28,28])
    sp_neff_100.append(nnz_sp)
    
    nnz_nm=np.count_nonzero(i_nm_wght,0)
    nnz_nm.reshape([28,28])
    nm_neff_100.append(nnz_nm)
      
    # Number of contacts arriving at each hidden neuron 
    nnz_sp=np.count_nonzero(i_sp_wght,1)
    sp_naff_100.append(nnz_sp)
    
    nnz_nm=np.count_nonzero(i_nm_wght,1)
    nm_naff_100.append(nnz_nm)
    
    # Weights 
    wght_sp=i_sp_wght[np.nonzero(i_sp_wght)]
    sp_wght_100.append(wght_sp)
    
    wght_nm=i_nm_wght[np.nonzero(i_nm_wght)]
    nm_wght_100.append(wght_nm)
    
    # Mean activations over test set
    a_sp=input_to_hidden_layer(net_sp, dig_test)
    sp_mnact_100.append(a_sp)
    
    a_nm=input_to_hidden_layer(net_nm, dig_test)
    nm_mnact_100.append(a_nm)
    
#%%    
coll_sp_neff_1=np.zeros([28,28]) # Number of contacts leaving each pixel
coll_nm_neff_1=np.zeros([28,28])
coll_sp_neff_10=np.zeros([28,28])
coll_nm_neff_10=np.zeros([28,28])
coll_sp_neff_100=np.zeros([28,28]) 
coll_nm_neff_100=np.zeros([28,28])

for ind in range(n_rep):
    coll_sp_neff_1=coll_sp_neff_1+np.reshape(sp_neff_1[ind],[28,28])/n_rep
    coll_nm_neff_1=coll_nm_neff_1+np.reshape(nm_neff_1[ind],[28,28])/n_rep
    coll_sp_neff_10=coll_sp_neff_10+np.reshape(sp_neff_10[ind],[28,28])/n_rep
    coll_nm_neff_10=coll_nm_neff_10+np.reshape(nm_neff_10[ind],[28,28])/n_rep
    coll_sp_neff_100=coll_sp_neff_100+np.reshape(sp_neff_100[ind],[28,28])/n_rep
    coll_nm_neff_100=coll_nm_neff_100+np.reshape(nm_neff_100[ind],[28,28])/n_rep
    
np.save('sim_data/Fig_2_neff_sp_1',coll_sp_neff_1)
np.save('sim_data/Fig_2_neff_nm_1',coll_nm_neff_1)
np.save('sim_data/Fig_2_neff_sp_10',coll_sp_neff_10)
np.save('sim_data/Fig_2_neff_nm_10',coll_nm_neff_10)
np.save('sim_data/Fig_2_neff_sp_100',coll_sp_neff_100)
np.save('sim_data/Fig_2_neff_nm_100',coll_nm_neff_100)


coll_sp_naff_1=[item for sublist in sp_naff_1 for item in sublist] # Number of contacts arriving at each hidden neuron 
coll_nm_naff_1=[item for sublist in nm_naff_1 for item in sublist] 
coll_sp_naff_5=[item for sublist in sp_naff_5 for item in sublist]  
coll_nm_naff_5=[item for sublist in nm_naff_5 for item in sublist] 
coll_sp_naff_10=[item for sublist in sp_naff_10 for item in sublist]
coll_nm_naff_10=[item for sublist in nm_naff_10 for item in sublist] 
coll_sp_naff_25=[item for sublist in sp_naff_25 for item in sublist] 
coll_nm_naff_25=[item for sublist in nm_naff_25 for item in sublist]
coll_sp_naff_100=[item for sublist in sp_naff_100 for item in sublist]
coll_nm_naff_100=[item for sublist in nm_naff_100 for item in sublist]
np.save('sim_data/Fig_2_naff_sp_1',coll_sp_naff_1)
np.save('sim_data/Fig_2_naff_nm_1',coll_nm_naff_1)
np.save('sim_data/Fig_2_naff_sp_5',coll_sp_naff_5)
np.save('sim_data/Fig_2_naff_nm_5',coll_nm_naff_5)
np.save('sim_data/Fig_2_naff_sp_10',coll_sp_naff_10)
np.save('sim_data/Fig_2_naff_nm_10',coll_nm_naff_10)
np.save('sim_data/Fig_2_naff_sp_25',coll_sp_naff_25)
np.save('sim_data/Fig_2_naff_nm_25',coll_nm_naff_25)
np.save('sim_data/Fig_2_naff_sp_100',coll_sp_naff_100)
np.save('sim_data/Fig_2_naff_nm_100',coll_nm_naff_100)


coll_sp_wght_1=[item for sublist in sp_wght_1 for item in sublist] # Weights 
coll_nm_wght_1=[item for sublist in nm_wght_1 for item in sublist] 
coll_sp_wght_5=[item for sublist in sp_wght_5 for item in sublist]  
coll_nm_wght_5=[item for sublist in nm_wght_5 for item in sublist] 
coll_sp_wght_10=[item for sublist in sp_wght_10 for item in sublist]
coll_nm_wght_10=[item for sublist in nm_wght_10 for item in sublist] 
coll_sp_wght_25=[item for sublist in sp_wght_25 for item in sublist] 
coll_nm_wght_25=[item for sublist in nm_wght_25 for item in sublist]
coll_sp_wght_100=[item for sublist in sp_wght_100 for item in sublist]
coll_nm_wght_100=[item for sublist in nm_wght_100 for item in sublist]
np.save('sim_data/Fig_2_wght_sp_1',coll_sp_wght_1)
np.save('sim_data/Fig_2_wght_nm_1',coll_nm_wght_1)
np.save('sim_data/Fig_2_wght_sp_5',coll_sp_wght_5)
np.save('sim_data/Fig_2_wght_nm_5',coll_nm_wght_5)
np.save('sim_data/Fig_2_wght_sp_10',coll_sp_wght_10)
np.save('sim_data/Fig_2_wght_nm_10',coll_nm_wght_10)
np.save('sim_data/Fig_2_wght_sp_25',coll_sp_wght_25)
np.save('sim_data/Fig_2_wght_nm_25',coll_nm_wght_25)
np.save('sim_data/Fig_2_wght_sp_100',coll_sp_wght_100)
np.save('sim_data/Fig_2_wght_nm_100',coll_nm_wght_100)


coll_sp_mnact_1=[item for sublist in sp_mnact_1 for item in sublist] # Mean activation over training set
coll_nm_mnact_1=[item for sublist in nm_mnact_1 for item in sublist] 
coll_sp_mnact_5=[item for sublist in sp_mnact_5 for item in sublist]  
coll_nm_mnact_5=[item for sublist in nm_mnact_5 for item in sublist] 
coll_sp_mnact_10=[item for sublist in sp_mnact_10 for item in sublist]
coll_nm_mnact_10=[item for sublist in nm_mnact_10 for item in sublist] 
coll_sp_mnact_25=[item for sublist in sp_mnact_25 for item in sublist] 
coll_nm_mnact_25=[item for sublist in nm_mnact_25 for item in sublist]
coll_sp_mnact_100=[item for sublist in sp_mnact_100 for item in sublist]
coll_nm_mnact_100=[item for sublist in nm_mnact_100 for item in sublist]
np.save('sim_data/Fig_2_mnact_sp_1',coll_sp_mnact_1)
np.save('sim_data/Fig_2_mnact_nm_1',coll_nm_mnact_1)
np.save('sim_data/Fig_2_mnact_sp_5',coll_sp_mnact_5)
np.save('sim_data/Fig_2_mnact_nm_5',coll_nm_mnact_5)
np.save('sim_data/Fig_2_mnact_sp_10',coll_sp_mnact_10)
np.save('sim_data/Fig_2_mnact_nm_10',coll_nm_mnact_10)
np.save('sim_data/Fig_2_mnact_sp_25',coll_sp_mnact_25)
np.save('sim_data/Fig_2_mnact_nm_25',coll_nm_mnact_25)
np.save('sim_data/Fig_2_mnact_sp_100',coll_sp_mnact_100)
np.save('sim_data/Fig_2_mnact_nm_100',coll_nm_mnact_100)
```
