## Supplementary material for "Dendritic normalisation improves learning in sparsely connected artificial neural networks": Code and data.: Fig_2_plot.py

```
# -*- coding: utf-8 -*-
"""
Created on Tue Nov 19 14:45:18 2019

@author: Alex Bird
"""
# Panel A, Efferent contact numbers
sp_neff_1=np.load('sim_data/Fig_2_neff_sp_1.npy')
nm_neff_1=np.load('sim_data/Fig_2_neff_nm_1.npy')
sp_neff_10=np.load('sim_data/Fig_2_neff_sp_10.npy')
nm_neff_10=np.load('sim_data/Fig_2_neff_nm_10.npy')
sp_neff_100=np.load('sim_data/Fig_2_neff_sp_100.npy')
nm_neff_100=np.load('sim_data/Fig_2_neff_nm_100.npy')

max_cons=max(np.max(sp_neff_1),np.max(sp_neff_10),np.max(sp_neff_100),np.max(nm_neff_1),np.max(nm_neff_10),np.max(nm_neff_100))

sns.set(style="white", palette="muted", color_codes=True)
f, axes = plt.subplots(3, 2, figsize=(2.3622,3.54331,), sharex=True)
sns.despine
sns.heatmap(sp_neff_1, vmin=0,vmax=max_cons,cbar=False,xticklabels=False,yticklabels=False,cmap='Blues',ax=axes[0,0])
sns.heatmap(sp_neff_10, vmin=0,vmax=max_cons,cbar=False,xticklabels=False,yticklabels=False,cmap='Blues',ax=axes[1,0])
sns.heatmap(sp_neff_100, vmin=0,vmax=max_cons,cbar=False,xticklabels=False,yticklabels=False,cmap='Blues',ax=axes[2,0])
sns.heatmap(nm_neff_1, vmin=0,vmax=max_cons,cbar=False,xticklabels=False,yticklabels=False,cmap='Oranges',ax=axes[0,1])
sns.heatmap(nm_neff_10, vmin=0,vmax=max_cons,cbar=False,xticklabels=False,yticklabels=False,cmap='Oranges',ax=axes[1,1])
sns.heatmap(nm_neff_100, vmin=0,vmax=max_cons,cbar=False,xticklabels=False,yticklabels=False,cmap='Oranges',ax=axes[2,1])
plt.savefig('fig/Fig_2_a.eps', format='eps')

sns.set(style="white", palette="muted", color_codes=True)
f, axes = plt.subplots(1, 2, figsize=(2.3622,1.1811), sharex=True)
sns.despine
sns.heatmap(sp_neff_1, vmin=0,vmax=max_cons,cbar=True,xticklabels=False,yticklabels=False,cmap='Blues',ax=axes[0])
sns.heatmap(sp_neff_10, vmin=0,vmax=max_cons,cbar=True,xticklabels=False,yticklabels=False,cmap='Oranges',ax=axes[1])
plt.savefig('fig/Fig_2_a_Colourbar.eps', format='eps')
#%%
# Panel B, Afferent contact numbers
res=10
#n_rep=np.load('sim_data/n_rep.npy')
n_rep=25
sp_naff_1=np.load('sim_data/Fig_2_naff_sp_1.npy')
nm_naff_1=np.load('sim_data/Fig_2_naff_nm_1.npy')
sp_naff_5=np.load('sim_data/Fig_2_naff_sp_5.npy')
nm_naff_5=np.load('sim_data/Fig_2_naff_nm_5.npy')
sp_naff_10=np.load('sim_data/Fig_2_naff_sp_10.npy')
nm_naff_10=np.load('sim_data/Fig_2_naff_nm_10.npy')
sp_naff_25=np.load('sim_data/Fig_2_naff_sp_25.npy')
nm_naff_25=np.load('sim_data/Fig_2_naff_nm_25.npy')
sp_naff_100=np.load('sim_data/Fig_2_naff_sp_100.npy')
nm_naff_100=np.load('sim_data/Fig_2_naff_nm_100.npy')


n_norm=n_rep*100
sns.set(style="white", palette="muted", color_codes=True)
f, axes = plt.subplots(5, 1, figsize=(1.1811,3.54331,), sharex=True , sharey=True)
sns.despine      
    
[sp_dist,sp_ends]=np.histogram(sp_naff_1,res)
[nm_dist,nm_ends]=np.histogram(nm_naff_1,res) 
sp_mids=(sp_ends[1:(res+1)]+sp_ends[0:res])/2
nm_mids=(nm_ends[1:(res+1)]+nm_ends[0:res])/2
axes[0].plot(sp_mids,sp_dist/np.trapz(sp_dist,sp_mids),'tab:blue',nm_mids,nm_dist/np.trapz(nm_dist,nm_mids),'tab:orange',linewidth=0.5)
#axes[0].fill_between(sp_mids,sp_dist/n_norm,0,facecolor='tab:blue',color='tab:blue',alpha=0.1)  
#axes[0].fill_between(nm_mids,nm_dist/n_norm,0,facecolor='tab:orange',color='tab:orange',alpha=0.1)
axes[0].spines['top'].set_visible(False)
axes[0].spines['right'].set_visible(False)
axes[0].spines['bottom'].set_edgecolor(None)
axes[0].spines['bottom'].set_facecolor('black')
axes[0].spines['bottom'].set_linewidth(1)
axes[0].spines['left'].set_edgecolor(None)
axes[0].spines['left'].set_facecolor('black')
axes[0].spines['left'].set_linewidth(1)
axes[0].set_xlim((100,200))
axes[0].set_xticks=[100,150,200]
axes[0].tick_params(direction='out', length=2, width=0.5 , labelsize=6 , pad=1)
 

[sp_dist,sp_ends]=np.histogram(sp_naff_5,res)
[nm_dist,nm_ends]=np.histogram(nm_naff_5,res) 
sp_mids=(sp_ends[1:(res+1)]+sp_ends[0:res])/2
nm_mids=(nm_ends[1:(res+1)]+nm_ends[0:res])/2
axes[1].plot(sp_mids,sp_dist/np.trapz(sp_dist,sp_mids),'tab:blue',nm_mids,nm_dist/np.trapz(nm_dist,nm_mids),'tab:orange',linewidth=0.5)
#axes[1].fill_between(sp_mids,sp_dist/n_norm,0,facecolor='tab:blue',color='tab:blue',alpha=0.1)  
#axes[1].fill_between(nm_mids,nm_dist/n_norm,0,facecolor='tab:orange',color='tab:orange',alpha=0.1)
axes[1].spines['top'].set_visible(False)
axes[1].spines['right'].set_visible(False)
axes[1].spines['bottom'].set_edgecolor(None)
axes[1].spines['bottom'].set_facecolor('black')
axes[1].spines['bottom'].set_linewidth(1)
axes[1].spines['left'].set_edgecolor(None)
axes[1].spines['left'].set_facecolor('black')
axes[1].spines['left'].set_linewidth(1)
axes[1].tick_params(direction='out', length=2, width=0.5, labelsize=6 , pad=1)  

[sp_dist,sp_ends]=np.histogram(sp_naff_10,res)
[nm_dist,nm_ends]=np.histogram(nm_naff_10,res) 
sp_mids=(sp_ends[1:(res+1)]+sp_ends[0:res])/2
nm_mids=(nm_ends[1:(res+1)]+nm_ends[0:res])/2
axes[2].plot(sp_mids,sp_dist/np.trapz(sp_dist,sp_mids),'tab:blue',nm_mids,nm_dist/np.trapz(nm_dist,nm_mids),'tab:orange',linewidth=0.5)
#axes[2].fill_between(sp_mids,sp_dist/n_norm,0,facecolor='tab:blue',color='tab:blue',alpha=0.1)  
#axes[2].fill_between(nm_mids,nm_dist/n_norm,0,facecolor='tab:orange',color='tab:orange',alpha=0.1)
axes[2].spines['top'].set_visible(False)
axes[2].spines['right'].set_visible(False)
axes[2].spines['bottom'].set_edgecolor(None)
axes[2].spines['bottom'].set_facecolor('black')
axes[2].spines['bottom'].set_linewidth(1)
axes[2].spines['left'].set_edgecolor(None)
axes[2].spines['left'].set_facecolor('black')
axes[2].spines['left'].set_linewidth(1)
axes[2].tick_params(direction='out', length=2, width=0.5 , labelsize=6, pad=1)

 
[sp_dist,sp_ends]=np.histogram(sp_naff_25,res)
[nm_dist,nm_ends]=np.histogram(nm_naff_25,res) 
sp_mids=(sp_ends[1:(res+1)]+sp_ends[0:res])/2
nm_mids=(nm_ends[1:(res+1)]+nm_ends[0:res])/2
axes[3].plot(sp_mids,sp_dist/np.trapz(sp_dist,sp_mids),'tab:blue',nm_mids,nm_dist/np.trapz(nm_dist,nm_mids),'tab:orange',linewidth=0.5)
#axes[3].fill_between(sp_mids,sp_dist/n_norm,0,facecolor='tab:blue',color='tab:blue',alpha=0.1)  
#axes[3].fill_between(nm_mids,nm_dist/n_norm,0,facecolor='tab:orange',color='tab:orange',alpha=0.1)  
axes[3].spines['top'].set_visible(False)
axes[3].spines['right'].set_visible(False)
axes[3].spines['bottom'].set_edgecolor(None)
axes[3].spines['bottom'].set_facecolor('black')
axes[3].spines['bottom'].set_linewidth(1)
axes[3].spines['left'].set_edgecolor(None)
axes[3].spines['left'].set_facecolor('black')
axes[3].spines['left'].set_linewidth(1)
axes[3].tick_params(direction='out', length=2, width=0.5 , labelsize=6, pad=1)

[sp_dist,sp_ends]=np.histogram(sp_naff_100,res)
[nm_dist,nm_ends]=np.histogram(nm_naff_100,res) 
sp_mids=(sp_ends[1:(res+1)]+sp_ends[0:res])/2
nm_mids=(nm_ends[1:(res+1)]+nm_ends[0:res])/2
axes[4].plot(sp_mids,sp_dist/np.trapz(sp_dist,sp_mids),'tab:blue',nm_mids,nm_dist/np.trapz(nm_dist,nm_mids),'tab:orange',linewidth=0.5)
#axes[4].fill_between(sp_mids,sp_dist/n_norm,0,facecolor='tab:blue',color='tab:blue',alpha=0.1)  
#axes[4].fill_between(nm_mids,nm_dist/n_norm,0,facecolor='tab:orange',color='tab:orange',alpha=0.1)
axes[4].spines['top'].set_visible(False)
axes[4].spines['right'].set_visible(False)
axes[0].spines['bottom'].set_edgecolor(None)
axes[4].spines['bottom'].set_facecolor('black')
axes[4].spines['bottom'].set_linewidth(1)
axes[4].spines['left'].set_edgecolor(None)
axes[4].spines['left'].set_facecolor('black')
axes[4].spines['left'].set_linewidth(1)
axes[4].tick_params(direction='out', length=2, width=0.5 , labelsize=6, pad=1) 
 
plt.savefig('fig/Fig_2_b.eps', format='eps',bbox_inches='tight',pad_inches=0.25)
#%%
# Panel C, Afferent weights
res=50
#n_rep=np.load('sim_data/n_rep.npy')
sp_wght_1=np.load('sim_data/Fig_2_wght_sp_1.npy')
nm_wght_1=np.load('sim_data/Fig_2_wght_nm_1.npy')
sp_wght_5=np.load('sim_data/Fig_2_wght_sp_5.npy')
nm_wght_5=np.load('sim_data/Fig_2_wght_nm_5.npy')
sp_wght_10=np.load('sim_data/Fig_2_wght_sp_10.npy')
nm_wght_10=np.load('sim_data/Fig_2_wght_nm_10.npy')
sp_wght_25=np.load('sim_data/Fig_2_wght_sp_25.npy')
nm_wght_25=np.load('sim_data/Fig_2_wght_nm_25.npy')
sp_wght_100=np.load('sim_data/Fig_2_wght_sp_100.npy')
nm_wght_100=np.load('sim_data/Fig_2_wght_nm_100.npy')


n_norm=len(sp_wght_1)
sns.set(style="white", palette="muted", color_codes=True)
f, axes = plt.subplots(5, 1, figsize=(1.1811,3.54331,), sharex=True, sharey=True)
sns.despine      
    
[sp_dist,sp_ends]=np.histogram(sp_wght_1,res)
[nm_dist,nm_ends]=np.histogram(nm_wght_1,res) 
sp_mids=(sp_ends[1:(res+1)]+sp_ends[0:res])/2
nm_mids=(nm_ends[1:(res+1)]+nm_ends[0:res])/2
axes[0].plot(sp_mids,sp_dist/np.trapz(sp_dist,sp_mids),'tab:blue',nm_mids,nm_dist/np.trapz(nm_dist,nm_mids),'tab:orange',linewidth=0.5)
#axes[0].fill_between(sp_mids,sp_dist/n_norm,0,facecolor='tab:blue',color='tab:blue',alpha=0.1)  
#axes[0].fill_between(nm_mids,nm_dist/n_norm,0,facecolor='tab:orange',color='tab:orange',alpha=0.1)
axes[0].spines['top'].set_visible(False)
axes[0].spines['right'].set_visible(False)
axes[0].spines['bottom'].set_facecolor('black')
axes[0].spines['bottom'].set_linewidth(1)
axes[0].spines['left'].set_edgecolor(None)
axes[0].spines['left'].set_facecolor('black')
axes[0].spines['left'].set_linewidth(1)
axes[0].set_xlim((-5,5))
axes[0].set_xticks=[-5,0,5]
axes[0].set_ylim((0,1))
axes[0].set_yticks=[0,1]
axes[0].tick_params(direction='out', length=2, width=0.5 , labelsize=6 , pad=1)


[sp_dist,sp_ends]=np.histogram(sp_wght_5,res)
[nm_dist,nm_ends]=np.histogram(nm_wght_5,res) 
sp_mids=(sp_ends[1:(res+1)]+sp_ends[0:res])/2
nm_mids=(nm_ends[1:(res+1)]+nm_ends[0:res])/2
axes[1].plot(sp_mids,sp_dist/np.trapz(sp_dist,sp_mids),'tab:blue',nm_mids,nm_dist/np.trapz(nm_dist,nm_mids),'tab:orange',linewidth=0.5)
#axes[1].fill_between(sp_mids,sp_dist/n_norm,0,facecolor='tab:blue',color='tab:blue',alpha=0.1)  
#axes[1].fill_between(nm_mids,nm_dist/n_norm,0,facecolor='tab:orange',color='tab:orange',alpha=0.1)
axes[1].spines['top'].set_visible(False)
axes[1].spines['right'].set_visible(False)
axes[1].spines['bottom'].set_edgecolor(None)
axes[1].spines['bottom'].set_facecolor('black')
axes[1].spines['bottom'].set_linewidth(1)
axes[1].spines['left'].set_edgecolor(None)
axes[1].spines['left'].set_facecolor('black')
axes[1].spines['left'].set_linewidth(1)
axes[1].tick_params(direction='out', length=2, width=0.5, labelsize=6 , pad=1) 

[sp_dist,sp_ends]=np.histogram(sp_wght_10,res)
[nm_dist,nm_ends]=np.histogram(nm_wght_10,res) 
sp_mids=(sp_ends[1:(res+1)]+sp_ends[0:res])/2
nm_mids=(nm_ends[1:(res+1)]+nm_ends[0:res])/2
axes[2].plot(sp_mids,sp_dist/np.trapz(sp_dist,sp_mids),'tab:blue',nm_mids,nm_dist/np.trapz(nm_dist,nm_mids),'tab:orange',linewidth=0.5)
#axes[2].fill_between(sp_mids,sp_dist/n_norm,0,facecolor='tab:blue',color='tab:blue',alpha=0.1)  
#axes[2].fill_between(nm_mids,nm_dist/n_norm,0,facecolor='tab:orange',color='tab:orange',alpha=0.1)
axes[2].spines['top'].set_visible(False)
axes[2].spines['right'].set_visible(False)
axes[1].spines['bottom'].set_edgecolor(None)
axes[2].spines['bottom'].set_facecolor('black')
axes[2].spines['bottom'].set_linewidth(1)
axes[2].spines['left'].set_edgecolor(None)
axes[2].spines['left'].set_facecolor('black')
axes[2].spines['left'].set_linewidth(1)
axes[2].tick_params(direction='out', length=2, width=0.5, labelsize=6 , pad=1) 

 
[sp_dist,sp_ends]=np.histogram(sp_wght_25,res)
[nm_dist,nm_ends]=np.histogram(nm_wght_25,res) 
sp_mids=(sp_ends[1:(res+1)]+sp_ends[0:res])/2
nm_mids=(nm_ends[1:(res+1)]+nm_ends[0:res])/2
axes[3].plot(sp_mids,sp_dist/np.trapz(sp_dist,sp_mids),'tab:blue',nm_mids,nm_dist/np.trapz(nm_dist,nm_mids),'tab:orange',linewidth=0.5)
#axes[3].fill_between(sp_mids,sp_dist/n_norm,0,facecolor='tab:blue',color='tab:blue',alpha=0.1)  
#axes[3].fill_between(nm_mids,nm_dist/n_norm,0,facecolor='tab:orange',color='tab:orange',alpha=0.1)  
axes[3].spines['top'].set_visible(False)
axes[3].spines['right'].set_visible(False)
axes[3].spines['bottom'].set_edgecolor(None)
axes[3].spines['bottom'].set_facecolor('black')
axes[3].spines['bottom'].set_linewidth(1)
axes[3].spines['left'].set_edgecolor(None)
axes[3].spines['left'].set_facecolor('black')
axes[3].spines['left'].set_linewidth(1)
axes[3].tick_params(direction='out', length=2, width=0.5, labelsize=6 , pad=1) 

[sp_dist,sp_ends]=np.histogram(sp_wght_100,res)
[nm_dist,nm_ends]=np.histogram(nm_wght_100,res) 
sp_mids=(sp_ends[1:(res+1)]+sp_ends[0:res])/2
nm_mids=(nm_ends[1:(res+1)]+nm_ends[0:res])/2
axes[4].plot(sp_mids,sp_dist/np.trapz(sp_dist,sp_mids),'tab:blue',nm_mids,nm_dist/np.trapz(nm_dist,nm_mids),'tab:orange',linewidth=0.5)
#axes[4].fill_between(sp_mids,sp_dist/n_norm,0,facecolor='tab:blue',color='tab:blue',alpha=0.1)  
#axes[4].fill_between(nm_mids,nm_dist/n_norm,0,facecolor='tab:orange',color='tab:orange',alpha=0.1)
axes[4].spines['top'].set_visible(False)
axes[4].spines['right'].set_visible(False)
axes[4].spines['bottom'].set_edgecolor(None)
axes[4].spines['bottom'].set_facecolor('black')
axes[4].spines['bottom'].set_linewidth(1)
axes[4].spines['left'].set_edgecolor(None)
axes[4].spines['left'].set_facecolor('black')
axes[4].spines['left'].set_linewidth(1)
axes[4].tick_params(direction='out', length=2, width=0.5, labelsize=6 , pad=1) 
 
plt.savefig('fig/Fig_2_c.eps', format='eps',bbox_inches='tight',pad_inches=0.25)


#%%
# Panel D, Mean activations
res=20
#n_rep=np.load('sim_data/n_rep.npy')
sp_mnact_1=np.load('sim_data/Fig_2_mnact_sp_1.npy')
nm_mnact_1=np.load('sim_data/Fig_2_mnact_nm_1.npy')
sp_mnact_5=np.load('sim_data/Fig_2_mnact_sp_5.npy')
nm_mnact_5=np.load('sim_data/Fig_2_mnact_nm_5.npy')
sp_mnact_10=np.load('sim_data/Fig_2_mnact_sp_10.npy')
nm_mnact_10=np.load('sim_data/Fig_2_mnact_nm_10.npy')
sp_mnact_25=np.load('sim_data/Fig_2_mnact_sp_25.npy')
nm_mnact_25=np.load('sim_data/Fig_2_mnact_nm_25.npy')
sp_mnact_100=np.load('sim_data/Fig_2_mnact_sp_100.npy')
nm_mnact_100=np.load('sim_data/Fig_2_mnact_nm_100.npy')


n_norm=n_rep*100
sns.set(style="white", palette="muted", color_codes=True)
f, axes = plt.subplots(5, 1, figsize=(1.1811,3.54331,), sharex=True , sharey=True)
sns.despine      
    
[sp_dist,sp_ends]=np.histogram(sp_mnact_1,res)
[nm_dist,nm_ends]=np.histogram(nm_mnact_1,res) 
sp_mids=(sp_ends[1:(res+1)]+sp_ends[0:res])/2
nm_mids=(nm_ends[1:(res+1)]+nm_ends[0:res])/2
axes[0].plot(sp_mids,sp_dist/np.trapz(sp_dist,sp_mids),'tab:blue',nm_mids,nm_dist/np.trapz(nm_dist,nm_mids),'tab:orange',linewidth=0.5)
#axes[0].fill_between(sp_mids,sp_dist/n_norm,0,facecolor='tab:blue',color='tab:blue',alpha=0.1)  
#axes[0].fill_between(nm_mids,nm_dist/n_norm,0,facecolor='tab:orange',color='tab:orange',alpha=0.1)
axes[0].spines['top'].set_visible(False)
axes[0].spines['right'].set_visible(False)
axes[0].spines['bottom'].set_edgecolor(None)
axes[0].spines['bottom'].set_facecolor('black')
axes[0].spines['bottom'].set_linewidth(1)
axes[0].spines['left'].set_edgecolor(None)
axes[0].spines['left'].set_facecolor('black')
axes[0].spines['left'].set_linewidth(1)
axes[0].set_xlim((-25,25))
axes[0].set_xticks=[-25,0,25]
axes[0].tick_params(direction='out', length=2, width=0.5 , labelsize=6 , pad=1)

 

[sp_dist,sp_ends]=np.histogram(sp_mnact_5,res)
[nm_dist,nm_ends]=np.histogram(nm_mnact_5,res) 
sp_mids=(sp_ends[1:(res+1)]+sp_ends[0:res])/2
nm_mids=(nm_ends[1:(res+1)]+nm_ends[0:res])/2
axes[1].plot(sp_mids,sp_dist/np.trapz(sp_dist,sp_mids),'tab:blue',nm_mids,nm_dist/np.trapz(nm_dist,nm_mids),'tab:orange',linewidth=0.5)
#axes[1].fill_between(sp_mids,sp_dist/n_norm,0,facecolor='tab:blue',color='tab:blue',alpha=0.1)  
#axes[1].fill_between(nm_mids,nm_dist/n_norm,0,facecolor='tab:orange',color='tab:orange',alpha=0.1)
axes[1].spines['top'].set_visible(False)
axes[1].spines['right'].set_visible(False)
axes[1].spines['bottom'].set_facecolor('black')
axes[1].spines['bottom'].set_linewidth(1)
axes[1].spines['left'].set_edgecolor(None)
axes[1].spines['left'].set_facecolor('black')
axes[1].spines['left'].set_linewidth(1)
axes[1].tick_params(direction='out', length=2, width=0.5 , labelsize=6 , pad=1)

[sp_dist,sp_ends]=np.histogram(sp_mnact_10,res)
[nm_dist,nm_ends]=np.histogram(nm_mnact_10,res) 
sp_mids=(sp_ends[1:(res+1)]+sp_ends[0:res])/2
nm_mids=(nm_ends[1:(res+1)]+nm_ends[0:res])/2
axes[2].plot(sp_mids,sp_dist/np.trapz(sp_dist,sp_mids),'tab:blue',nm_mids,nm_dist/np.trapz(nm_dist,nm_mids),'tab:orange',linewidth=0.5)
#axes[2].fill_between(sp_mids,sp_dist/n_norm,0,facecolor='tab:blue',color='tab:blue',alpha=0.1)  
#axes[2].fill_between(nm_mids,nm_dist/n_norm,0,facecolor='tab:orange',color='tab:orange',alpha=0.1)
axes[2].spines['top'].set_visible(False)
axes[2].spines['right'].set_visible(False)
axes[2].spines['bottom'].set_facecolor('black')
axes[2].spines['bottom'].set_linewidth(1)
axes[2].spines['left'].set_edgecolor(None)
axes[2].spines['left'].set_facecolor('black')
axes[2].spines['left'].set_linewidth(1)
axes[2].tick_params(direction='out', length=2, width=0.5 , labelsize=6 , pad=1)

 
[sp_dist,sp_ends]=np.histogram(sp_mnact_25,res)
[nm_dist,nm_ends]=np.histogram(nm_mnact_25,res) 
sp_mids=(sp_ends[1:(res+1)]+sp_ends[0:res])/2
nm_mids=(nm_ends[1:(res+1)]+nm_ends[0:res])/2
axes[3].plot(sp_mids,sp_dist/np.trapz(sp_dist,sp_mids),'tab:blue',nm_mids,nm_dist/np.trapz(nm_dist,nm_mids),'tab:orange',linewidth=0.5)
#axes[3].fill_between(sp_mids,sp_dist/n_norm,0,facecolor='tab:blue',color='tab:blue',alpha=0.1)  
#axes[3].fill_between(nm_mids,nm_dist/n_norm,0,facecolor='tab:orange',color='tab:orange',alpha=0.1)  
axes[3].spines['top'].set_visible(False)
axes[3].spines['right'].set_visible(False)
axes[3].spines['bottom'].set_facecolor('black')
axes[3].spines['bottom'].set_linewidth(1)
axes[3].spines['left'].set_edgecolor(None)
axes[3].spines['left'].set_facecolor('black')
axes[3].spines['left'].set_linewidth(1)
axes[3].tick_params(direction='out', length=2, width=0.5 , labelsize=6 , pad=1)

[sp_dist,sp_ends]=np.histogram(sp_mnact_100,res)
[nm_dist,nm_ends]=np.histogram(nm_mnact_100,res) 
sp_mids=(sp_ends[1:(res+1)]+sp_ends[0:res])/2
nm_mids=(nm_ends[1:(res+1)]+nm_ends[0:res])/2
axes[4].plot(sp_mids,sp_dist/np.trapz(sp_dist,sp_mids),'tab:blue',nm_mids,nm_dist/np.trapz(nm_dist,nm_mids),'tab:orange',linewidth=0.5)
#axes[4].fill_between(sp_mids,sp_dist/n_norm,0,facecolor='tab:blue',color='tab:blue',alpha=0.1)  
#axes[4].fill_between(nm_mids,nm_dist/n_norm,0,facecolor='tab:orange',color='tab:orange',alpha=0.1)
axes[4].spines['top'].set_visible(False)
axes[4].spines['right'].set_visible(False)
axes[4].spines['bottom'].set_facecolor('black')
axes[4].spines['bottom'].set_linewidth(1)
axes[4].spines['left'].set_edgecolor(None)
axes[4].spines['left'].set_facecolor('black')
axes[4].spines['left'].set_linewidth(1)
axes[4].tick_params(direction='out', length=2, width=0.5 , labelsize=6 , pad=1)
 
plt.savefig('fig/Fig_2_d.eps', format='eps',bbox_inches='tight',pad_inches=0.25)
```
