## Supplementary material for "Dendritic normalisation improves learning in sparsely connected artificial neural networks": Code and data.: Fig_3_data_gen.py

```
#!/usr/bin/env python3
# -*- coding: utf-8 -*-
"""
Created on Wed Dec 11 20:19:33 2019

@author: Alex
"""

###############################################################################
###############################################################################
###############################################################################
# Fashion with 2 hidden layers

n_epoch=50
n_rep=20

sp_all_accs=np.zeros([n_rep,n_epoch])
sp_all_costs=np.zeros([n_rep,n_epoch])
nm_all_accs=np.zeros([n_rep,n_epoch])
nm_all_costs=np.zeros([n_rep,n_epoch])

for rep_ind in range(n_rep):
    runfile('/Users/Alex/Dendritic_normalisation/network_sp_2lyr.py', wdir='/Users/Alex/Dendritic_normalisation')
    sp_all_accs[rep_ind]=accuracies
    sp_all_costs[rep_ind]=costs
    print('Two layers control '+str(rep_ind))
    runfile('/Users/Alex/Dendritic_normalisation/network_nm_2lyr.py', wdir='/Users/Alex/Dendritic_normalisation')
    nm_all_accs[rep_ind]=accuracies
    nm_all_costs[rep_ind]=costs 
    print('Two layers norm '+str(rep_ind))
    
np.save('sim_data/Fig_3_2lyr_sp_cst',sp_all_costs)
np.save('sim_data/Fig_3_2lyr_sp_acc',sp_all_accs)
np.save('sim_data/Fig_3_2lyr_nm_cst',nm_all_costs)
np.save('sim_data/Fig_3_2lyr_nm_acc',nm_all_accs)

#%%
###############################################################################
###############################################################################
###############################################################################

# Fashion with three layers of 100 hidden neurons

n_epoch=50
n_rep=20

sp_all_accs=np.zeros([n_rep,n_epoch])
sp_all_costs=np.zeros([n_rep,n_epoch])
nm_all_accs=np.zeros([n_rep,n_epoch])
nm_all_costs=np.zeros([n_rep,n_epoch])

for rep_ind in range(n_rep):
    runfile('/Users/Alex/Dendritic_normalisation/network_sp_3lyr.py', wdir='/Users/Alex/Dendritic_normalisation')
    sp_all_accs[rep_ind]=accuracies
    sp_all_costs[rep_ind]=costs
    print('Three layers control '+str(rep_ind))
    runfile('/Users/Alex/Dendritic_normalisation/network_nm_3lyr.py', wdir='/Users/Alex/Dendritic_normalisation')
    nm_all_accs[rep_ind]=accuracies
    nm_all_costs[rep_ind]=costs
    print('Three layers norm '+str(rep_ind))
    
np.save('sim_data/Fig_3_3lyr_sp_cst',sp_all_costs)
np.save('sim_data/Fig_3_3lyr_sp_acc',sp_all_accs)
np.save('sim_data/Fig_3_3lyr_nm_cst',nm_all_costs)
np.save('sim_data/Fig_3_3lyr_nm_acc',nm_all_accs)

#%%
###############################################################################
###############################################################################
###############################################################################
#Fashion with convolutional nets
#
#n_epoch=50
#n_rep=10
#
#sp_all_accs=np.zeros([n_rep,n_epoch])
#sp_all_costs=np.zeros([n_rep,n_epoch])
#nm_all_accs=np.zeros([n_rep,n_epoch])
#nm_all_costs=np.zeros([n_rep,n_epoch])
#
#for rep_ind in range(n_rep):
#    runfile('/Users/Alex/Dendritic_normalisation/network_sp_conv.py', wdir='/Users/Alex/Dendritic_normalisation')
#    sp_all_accs[rep_ind]=accuracies
#    sp_all_costs[rep_ind]=costs
#    runfile('/Users/Alex/Dendritic_normalisation/network_nm_conv.py', wdir='/Users/Alex/Dendritic_normalisation')
#    nm_all_accs[rep_ind]=accuracies
#    nm_all_costs[rep_ind]=costs
#    
#np.save('sim_data/Fig_3_conv_sp_cst',sp_all_costs)
#np.save('sim_data/Fig_3_conv_sp_acc',sp_all_accs)
#np.save('sim_data/Fig_3_conv_nm_cst',nm_all_costs)
#np.save('sim_data/Fig_3_conv_nm_acc',nm_all_accs)


#%%
###############################################################################
###############################################################################
###############################################################################
# Fashion with threshold linear cells

n_epoch=50
n_rep=20

sp_all_accs=np.zeros([n_rep,n_epoch])
sp_all_costs=np.zeros([n_rep,n_epoch])
nm_all_accs=np.zeros([n_rep,n_epoch])
nm_all_costs=np.zeros([n_rep,n_epoch])

for rep_ind in range(n_rep):
    runfile('/Users/Alex/Dendritic_normalisation/network_sp_thlin.py', wdir='/Users/Alex/Dendritic_normalisation')
    sp_all_accs[rep_ind]=accuracies
    sp_all_costs[rep_ind]=costs
    print('Thlin control '+str(rep_ind))
    runfile('/Users/Alex/Dendritic_normalisation/network_nm_thlin.py', wdir='/Users/Alex/Dendritic_normalisation')
    nm_all_accs[rep_ind]=accuracies
    nm_all_costs[rep_ind]=costs
    print('Thlin norm '+str(rep_ind))
    
    
np.save('sim_data/Fig_3_thlin_sp_cst',sp_all_costs)
np.save('sim_data/Fig_3_thlin_sp_acc',sp_all_accs)
np.save('sim_data/Fig_3_thlin_nm_cst',nm_all_costs)
np.save('sim_data/Fig_3_thlin_nm_acc',nm_all_accs)


#%%
###############################################################################
###############################################################################
###############################################################################
# Fashion with different norms

n_epoch=50
n_rep=20

L2_nm_all_accs=np.zeros([n_rep,n_epoch])
L2_nm_all_costs=np.zeros([n_rep,n_epoch])
L0_nm_all_accs=np.zeros([n_rep,n_epoch])
L0_nm_all_costs=np.zeros([n_rep,n_epoch])
L2L0_nm_all_accs=np.zeros([n_rep,n_epoch])
L2L0_nm_all_costs=np.zeros([n_rep,n_epoch])

for rep_ind in range(n_rep):
    runfile('/Users/Alex/Dendritic_normalisation/network_nm_L2.py', wdir='/Users/Alex/Dendritic_normalisation')
    L2_nm_all_accs[rep_ind]=accuracies
    L2_nm_all_costs[rep_ind]=costs
    print('L2 '+str(rep_ind))
    runfile('/Users/Alex/Dendritic_normalisation/network_nm_L0.py', wdir='/Users/Alex/Dendritic_normalisation')
    L0_nm_all_accs[rep_ind]=accuracies
    L0_nm_all_costs[rep_ind]=costs
    print('L0'+str(rep_ind))
    runfile('/Users/Alex/Dendritic_normalisation/network_nm_L2_and_L0.py', wdir='/Users/Alex/Dendritic_normalisation')
    L2L0_nm_all_accs[rep_ind]=accuracies
    L2L0_nm_all_costs[rep_ind]=costs
    print('Both '+str(rep_ind))
    
np.save('sim_data/Fig_3_L2_nm_cst',L2_nm_all_costs)
np.save('sim_data/Fig_3_L2_nm_acc',L2_nm_all_accs)
np.save('sim_data/Fig_3_L0_nm_cst',L0_nm_all_costs)
np.save('sim_data/Fig_3_L0_nm_acc',L0_nm_all_accs)
np.save('sim_data/Fig_3_L2L0_nm_cst',L2L0_nm_all_costs)
np.save('sim_data/Fig_3_L2L0_nm_acc',L2L0_nm_all_accs)


#%%
###############################################################################
###############################################################################
###############################################################################
# Fashion with different p-norms

epochs=20
n_rep=20
val_vec=[1,5,10,20]

const_accs=np.zeros([6,np.size(val_vec),n_rep])
var_accs=np.zeros([6,np.size(val_vec),n_rep])

for rep_ind in range(n_rep):
    # Constant excitability
    print('L0-norm, rep '+str(rep_ind)+': Constant excitability')
    runfile('/Users/Alex/Dendritic_normalisation/network_nm_L0.py', wdir='/Users/Alex/Dendritic_normalisation')
    i_accs=accuracies
    for ind in range(np.size(val_vec)):
        val=val_vec[ind]-1
        const_accs[0,ind,rep_ind]=i_accs[val]
    for p in range(1,6):
        print('L'+str(p)+'-norm, rep '+str(rep_ind)+': Constant excitability')
        runfile('/Users/Alex/Dendritic_normalisation/network_nm_Lp_const.py', wdir='/Users/Alex/Dendritic_normalisation')
        i_accs=accuracies
        for ind in range(np.size(val_vec)):
            val=val_vec[ind]-1
            const_accs[p,ind,rep_ind]=i_accs[val]
    # Variable excitability
    print('L0-norm, rep '+str(rep_ind)+': Variable excitability')
    runfile('/Users/Alex/Dendritic_normalisation/network_nm_L0_var.py', wdir='/Users/Alex/Dendritic_normalisation')
    i_accs=accuracies
    for ind in range(np.size(val_vec)):
        val=val_vec[ind]-1
        var_accs[0,ind,rep_ind]=i_accs[val]
    for p in range(1,6):
        print('L'+str(p)+'-norm, rep '+str(rep_ind)+': Variable excitability')
        runfile('/Users/Alex/Dendritic_normalisation/network_nm_Lp_var.py', wdir='/Users/Alex/Dendritic_normalisation')
        i_accs=accuracies
        for ind in range(np.size(val_vec)):
            val=val_vec[ind]-1
            var_accs[p,ind,rep_ind]=i_accs[val]
    
np.save('sim_data/Fig_3_Lp_const',const_accs)
np.save('sim_data/Fig_3_Lp_var',var_accs)
```
