## Supplementary material for "Dendritic normalisation improves learning in sparsely connected artificial neural networks": Code and data.: Fig_3_plot.py

```
#!/usr/bin/env python3
# -*- coding: utf-8 -*-
"""
Created on Fri Dec 13 22:53:39 2019

@author: Alex
"""
# Multiple layer networks


sp_2lyr_cst=np.load('sim_data/Fig_3_2lyr_sp_cst.npy')
nm_2lyr_cst=np.load('sim_data/Fig_3_2lyr_nm_cst.npy')
sp_2lyr_acc=np.load('sim_data/Fig_3_2lyr_sp_acc.npy')
nm_2lyr_acc=np.load('sim_data/Fig_3_2lyr_nm_acc.npy')

n_rep=sp_2lyr_cst.shape[0]
n_epoch=sp_2lyr_cst.shape[1]
sp_2lyr_cst_mn=np.zeros(n_epoch)
sp_2lyr_cst_sd=np.zeros(n_epoch)
sp_2lyr_acc_mn=np.zeros(n_epoch)
sp_2lyr_acc_sd=np.zeros(n_epoch)
nm_2lyr_cst_mn=np.zeros(n_epoch)
nm_2lyr_cst_sd=np.zeros(n_epoch)
nm_2lyr_acc_mn=np.zeros(n_epoch)
nm_2lyr_acc_sd=np.zeros(n_epoch)

for ep_ind in range(n_epoch):
    vec_sp_cst=sp_2lyr_cst[:,ep_ind]
    sp_2lyr_cst_mn[ep_ind]=np.mean(vec_sp_cst)
    sp_2lyr_cst_sd[ep_ind]=(n_rep/(n_rep-1)*np.var(vec_sp_cst))**0.5
    vec_sp_acc=sp_2lyr_acc[:,ep_ind]
    sp_2lyr_acc_mn[ep_ind]=np.mean(vec_sp_acc)
    sp_2lyr_acc_sd[ep_ind]=(n_rep/(n_rep-1)*np.var(vec_sp_acc))**0.5
    vec_nm_cst=nm_2lyr_cst[:,ep_ind]
    nm_2lyr_cst_mn[ep_ind]=np.mean(vec_nm_cst)
    nm_2lyr_cst_sd[ep_ind]=(n_rep/(n_rep-1)*np.var(vec_nm_cst))**0.5
    vec_nm_acc=nm_2lyr_acc[:,ep_ind]
    nm_2lyr_acc_mn[ep_ind]=np.mean(vec_nm_acc)
    nm_2lyr_acc_sd[ep_ind]=(n_rep/(n_rep-1)*np.var(vec_nm_acc))**0.5
    
sp_3lyr_cst=np.load('sim_data/Fig_3_3lyr_sp_cst.npy')
nm_3lyr_cst=np.load('sim_data/Fig_3_3lyr_nm_cst.npy')
sp_3lyr_acc=np.load('sim_data/Fig_3_3lyr_sp_acc.npy')
nm_3lyr_acc=np.load('sim_data/Fig_3_3lyr_nm_acc.npy')

n_rep=sp_3lyr_cst.shape[0]
n_epoch=sp_3lyr_cst.shape[1]
sp_3lyr_cst_mn=np.zeros(n_epoch)
sp_3lyr_cst_sd=np.zeros(n_epoch)
sp_3lyr_acc_mn=np.zeros(n_epoch)
sp_3lyr_acc_sd=np.zeros(n_epoch)
nm_3lyr_cst_mn=np.zeros(n_epoch)
nm_3lyr_cst_sd=np.zeros(n_epoch)
nm_3lyr_acc_mn=np.zeros(n_epoch)
nm_3lyr_acc_sd=np.zeros(n_epoch)
for ep_ind in range(n_epoch):
    vec_sp_cst=sp_3lyr_cst[:,ep_ind]
    sp_3lyr_cst_mn[ep_ind]=np.mean(vec_sp_cst)
    sp_3lyr_cst_sd[ep_ind]=(n_rep/(n_rep-1)*np.var(vec_sp_cst))**0.5
    vec_sp_acc=sp_3lyr_acc[:,ep_ind]
    sp_3lyr_acc_mn[ep_ind]=np.mean(vec_sp_acc)
    sp_3lyr_acc_sd[ep_ind]=(n_rep/(n_rep-1)*np.var(vec_sp_acc))**0.5
    vec_nm_cst=nm_3lyr_cst[:,ep_ind]
    nm_3lyr_cst_mn[ep_ind]=np.mean(vec_nm_cst)
    nm_3lyr_cst_sd[ep_ind]=(n_rep/(n_rep-1)*np.var(vec_nm_cst))**0.5
    vec_nm_acc=nm_3lyr_acc[:,ep_ind]
    nm_3lyr_acc_mn[ep_ind]=np.mean(vec_nm_acc)
    nm_3lyr_acc_sd[ep_ind]=(n_rep/(n_rep-1)*np.var(vec_nm_acc))**0.5

sns.set(style="white", palette="muted", color_codes=True)
f, axes = plt.subplots(4, 1, figsize=(1.1811,2.3622), sharex=True)
sns.despine      
    
axes[0].plot(range(0,50),sp_2lyr_cst_mn,'tab:blue',range(0,50),nm_2lyr_cst_mn,'tab:orange',linewidth=0.5)
axes[0].fill_between(range(0,50),sp_2lyr_cst_mn-sp_2lyr_cst_sd,sp_2lyr_cst_mn+sp_2lyr_cst_sd,facecolor='tab:blue',color=None,alpha=0.2)
axes[0].fill_between(range(0,50),nm_2lyr_cst_mn-nm_2lyr_cst_sd,nm_2lyr_cst_mn+nm_2lyr_cst_sd,facecolor='tab:orange',color=None,alpha=0.2)      
axes[0].spines['top'].set_visible(False)
axes[0].spines['right'].set_visible(False)
axes[0].spines['bottom'].set_edgecolor('black')
axes[0].spines['bottom'].set_facecolor('black')
axes[0].spines['bottom'].set_linewidth(0.5)
axes[0].spines['left'].set_edgecolor('black')
axes[0].spines['left'].set_facecolor('black')
axes[0].spines['left'].set_linewidth(0.5)
axes[0].set_xticks([0,25,50])
axes[0].set_yticks([0.2,0.5,0.8],[0.2,0.5,0.8])
axes[0].tick_params(direction='out', length=2, width=1 , labelsize=6 , pad=0.5)
 

axes[1].plot(range(0,50),sp_2lyr_acc_mn,'tab:blue',range(0,50),nm_2lyr_acc_mn,'tab:orange',linewidth=0.5)
axes[1].fill_between(range(0,50),sp_2lyr_acc_mn-sp_2lyr_acc_sd,sp_2lyr_acc_mn+sp_2lyr_acc_sd,facecolor='tab:blue',color=None,alpha=0.2)
axes[1].fill_between(range(0,50),nm_2lyr_acc_mn-nm_2lyr_acc_sd,nm_2lyr_acc_mn+nm_2lyr_acc_sd,facecolor='tab:orange',color=None,alpha=0.2)      
axes[1].spines['top'].set_visible(False)
axes[1].spines['right'].set_visible(False)
axes[1].spines['bottom'].set_edgecolor('black')
axes[1].spines['bottom'].set_facecolor('black')
axes[1].spines['bottom'].set_linewidth(0.5)
axes[1].spines['left'].set_edgecolor('black')
axes[1].spines['left'].set_facecolor('black')
axes[1].spines['left'].set_linewidth(0.5)
axes[1].set_xticks([0,25,50])
axes[1].set_yticks([0.75,0.85],[0.75,0.85])
axes[1].tick_params(direction='out', length=2, width=1 , labelsize=6 , pad=0.5)
 
axes[2].plot(range(0,50),sp_3lyr_cst_mn,'tab:blue',range(0,50),nm_3lyr_cst_mn,'tab:orange',linewidth=0.5)
axes[2].fill_between(range(0,50),sp_3lyr_cst_mn-sp_3lyr_cst_sd,sp_3lyr_cst_mn+sp_3lyr_cst_sd,facecolor='tab:blue',color=None,alpha=0.2)
axes[2].fill_between(range(0,50),nm_3lyr_cst_mn-nm_3lyr_cst_sd,nm_3lyr_cst_mn+nm_3lyr_cst_sd,facecolor='tab:orange',color=None,alpha=0.2)      
axes[2].spines['top'].set_visible(False)
axes[2].spines['right'].set_visible(False)
axes[2].spines['bottom'].set_edgecolor('black')
axes[2].spines['bottom'].set_facecolor('black')
axes[2].spines['bottom'].set_linewidth(0.5)
axes[2].spines['left'].set_edgecolor('black')
axes[2].spines['left'].set_facecolor('black')
axes[2].spines['left'].set_linewidth(0.5)
axes[2].set_xticks([0,25,50])
axes[2].set_yticks([0.2,0.5,0.8],[0.2,0.5,0.8])
axes[2].tick_params(direction='out', length=2, width=1 , labelsize=6 , pad=0.5)
 

axes[3].plot(range(0,50),sp_3lyr_acc_mn,'tab:blue',range(0,50),nm_3lyr_acc_mn,'tab:orange',linewidth=0.5)
axes[3].fill_between(range(0,50),sp_3lyr_acc_mn-sp_3lyr_acc_sd,sp_3lyr_acc_mn+sp_3lyr_acc_sd,facecolor='tab:blue',color=None,alpha=0.2)
axes[3].fill_between(range(0,50),nm_3lyr_acc_mn-nm_3lyr_acc_sd,nm_3lyr_acc_mn+nm_3lyr_acc_sd,facecolor='tab:orange',color=None,alpha=0.2)      
axes[3].spines['top'].set_visible(False)
axes[3].spines['right'].set_visible(False)
axes[3].spines['bottom'].set_edgecolor('black')
axes[3].spines['bottom'].set_facecolor('black')
axes[3].spines['bottom'].set_linewidth(0.5)
axes[3].spines['left'].set_edgecolor('black')
axes[3].spines['left'].set_facecolor('black')
axes[3].spines['left'].set_linewidth(0.5)
axes[3].set_xticks([0,25,50])
axes[3].set_yticks([0.75,0.85],[0.75,0.85])
axes[3].tick_params(direction='out', length=2, width=1 , labelsize=6 , pad=0.5)

plt.savefig('fig/Fig_3_c.pdf', format='pdf',bbox_inches='tight',pad_inches=0.25)

#%%
sp_conv_cst=np.load('sim_data/Fig_3_conv_sp_cst.npy')
nm_conv_cst=np.load('sim_data/Fig_3_conv_nm_cst.npy')
sp_conv_acc=np.load('sim_data/Fig_3_conv_sp_acc.npy')
nm_conv_acc=np.load('sim_data/Fig_3_conv_nm_acc.npy')

n_rep=sp_conv_cst.shape[0]
n_epoch=sp_conv_cst.shape[1]
sp_conv_cst_mn=np.zeros(n_epoch)
sp_conv_cst_sd=np.zeros(n_epoch)
sp_conv_acc_mn=np.zeros(n_epoch)
sp_conv_acc_sd=np.zeros(n_epoch)
nm_conv_cst_mn=np.zeros(n_epoch)
nm_conv_cst_sd=np.zeros(n_epoch)
nm_conv_acc_mn=np.zeros(n_epoch)
nm_conv_acc_sd=np.zeros(n_epoch)
for ep_ind in range(n_epoch):
    vec_sp_cst=sp_conv_cst[:,ep_ind]
    sp_conv_cst_mn[ep_ind]=np.mean(vec_sp_cst)
    sp_conv_cst_sd[ep_ind]=(n_rep/(n_rep-1)*np.var(vec_sp_cst))**0.5
    vec_sp_acc=sp_conv_acc[:,ep_ind]
    sp_conv_acc_mn[ep_ind]=np.mean(vec_sp_acc)
    sp_conv_acc_sd[ep_ind]=(n_rep/(n_rep-1)*np.var(vec_sp_acc))**0.5
    vec_nm_cst=nm_conv_cst[:,ep_ind]
    nm_conv_cst_mn[ep_ind]=np.mean(vec_nm_cst)
    nm_conv_cst_sd[ep_ind]=(n_rep/(n_rep-1)*np.var(vec_nm_cst))**0.5
    vec_nm_acc=nm_conv_acc[:,ep_ind]
    nm_conv_acc_mn[ep_ind]=np.mean(vec_nm_acc)
    nm_conv_acc_sd[ep_ind]=(n_rep/(n_rep-1)*np.var(vec_nm_acc))**0.5
    
sns.set(style="white", palette="muted", color_codes=True)
f, axes = plt.subplots(2, 1, figsize=(1.1811,1.2811), sharex=True)
sns.despine      
    
axes[0].plot(range(n_epoch),sp_conv_cst_mn,'tab:blue',range(n_epoch),nm_conv_cst_mn,'tab:orange',linewidth=0.5)
axes[0].fill_between(range(n_epoch),sp_conv_cst_mn-sp_conv_cst_sd,sp_conv_cst_mn+sp_conv_cst_sd,facecolor='tab:blue',color=None,alpha=0.2)
axes[0].fill_between(range(n_epoch),nm_conv_cst_mn-nm_conv_cst_sd,nm_conv_cst_mn+nm_conv_cst_sd,facecolor='tab:orange',color=None,alpha=0.2)      
axes[0].spines['top'].set_visible(False)
axes[0].spines['right'].set_visible(False)
axes[0].spines['bottom'].set_edgecolor('black')
axes[0].spines['bottom'].set_facecolor('black')
axes[0].spines['bottom'].set_linewidth(0.5)
axes[0].spines['left'].set_edgecolor('black')
axes[0].spines['left'].set_facecolor('black')
axes[0].spines['left'].set_linewidth(0.5)
axes[0].set_xticks([0,25,50])
axes[0].set_yticks([0.2,0.5,0.8],[0.2,0.5,0.8])
axes[0].tick_params(direction='out', length=2, width=1 , labelsize=6 , pad=1)
 

axes[1].plot(range(n_epoch),sp_conv_acc_mn,'tab:blue',range(n_epoch),nm_conv_acc_mn,'tab:orange',linewidth=0.5)
axes[1].fill_between(range(n_epoch),sp_conv_acc_mn-sp_conv_acc_sd,sp_conv_acc_mn+sp_conv_acc_sd,facecolor='tab:blue',color=None,alpha=0.2)
axes[1].fill_between(range(n_epoch),nm_conv_acc_mn-nm_conv_acc_sd,nm_conv_acc_mn+nm_conv_acc_sd,facecolor='tab:orange',color=None,alpha=0.2)      
axes[1].spines['top'].set_visible(False)
axes[1].spines['right'].set_visible(False)
axes[1].spines['bottom'].set_edgecolor('black')
axes[1].spines['bottom'].set_facecolor('black')
axes[1].spines['bottom'].set_linewidth(0.5)
axes[1].spines['left'].set_edgecolor('black')
axes[1].spines['left'].set_facecolor('black')
axes[1].spines['left'].set_linewidth(0.5)
axes[1].set_xticks([0,25,50])
#axes[1].set_yticks([0.2,0.5,0.8],[0.2,0.5,0.8])
axes[1].tick_params(direction='out', length=2, width=1 , labelsize=6 , pad=1)

plt.savefig('fig/Fig_3_d.pdf', format='pdf',bbox_inches='tight',pad_inches=0.25)

#%%
sp_th_cst=np.load('sim_data/Fig_3_thlin_sp_cst.npy')
nm_th_cst=np.load('sim_data/Fig_3_thlin_nm_cst.npy')
sp_th_acc=np.load('sim_data/Fig_3_thlin_sp_acc.npy')
nm_th_acc=np.load('sim_data/Fig_3_thlin_nm_acc.npy')

n_rep=sp_th_cst.shape[0]
n_epoch=sp_th_cst.shape[1]

sp_th_cst_mn=np.zeros(n_epoch)
sp_th_cst_sd=np.zeros(n_epoch)
sp_th_acc_mn=np.zeros(n_epoch)
sp_th_acc_sd=np.zeros(n_epoch)
nm_th_cst_mn=np.zeros(n_epoch)
nm_th_cst_sd=np.zeros(n_epoch)
nm_th_acc_mn=np.zeros(n_epoch)
nm_th_acc_sd=np.zeros(n_epoch)
for ep_ind in range(n_epoch):
    vec_sp_cst=sp_th_cst[:,ep_ind]
    sp_th_cst_mn[ep_ind]=np.mean(vec_sp_cst)
    sp_th_cst_sd[ep_ind]=(n_rep/(n_rep-1)*np.var(vec_sp_cst))**0.5
    vec_sp_acc=sp_th_acc[:,ep_ind]
    sp_th_acc_mn[ep_ind]=np.mean(vec_sp_acc)
    sp_th_acc_sd[ep_ind]=(n_rep/(n_rep-1)*np.var(vec_sp_acc))**0.5
    vec_nm_cst=nm_th_cst[:,ep_ind]
    nm_th_cst_mn[ep_ind]=np.mean(vec_nm_cst)
    nm_th_cst_sd[ep_ind]=(n_rep/(n_rep-1)*np.var(vec_nm_cst))**0.5
    vec_nm_acc=nm_th_acc[:,ep_ind]
    nm_th_acc_mn[ep_ind]=np.mean(vec_nm_acc)
    nm_th_acc_sd[ep_ind]=(n_rep/(n_rep-1)*np.var(vec_nm_acc))**0.5     
        
sns.set(style="white", palette="muted", color_codes=True)
f, axes = plt.subplots(2, 1, figsize=(1.1811,1.2811), sharex=True)
sns.despine      
    
axes[0].plot(range(n_epoch),sp_th_cst_mn,'tab:blue',range(n_epoch),nm_th_cst_mn,'tab:orange',linewidth=0.5)
axes[0].fill_between(range(n_epoch),sp_th_cst_mn-sp_th_cst_sd,sp_th_cst_mn+sp_th_cst_sd,facecolor='tab:blue',color=None,alpha=0.2)
axes[0].fill_between(range(n_epoch),nm_th_cst_mn-nm_th_cst_sd,nm_th_cst_mn+nm_th_cst_sd,facecolor='tab:orange',color=None,alpha=0.2)      
axes[0].spines['top'].set_visible(False)
axes[0].spines['right'].set_visible(False)
axes[0].spines['bottom'].set_edgecolor('black')
axes[0].spines['bottom'].set_facecolor('black')
axes[0].spines['bottom'].set_linewidth(0.5)
axes[0].spines['left'].set_edgecolor('black')
axes[0].spines['left'].set_facecolor('black')
axes[0].spines['left'].set_linewidth(0.5)
axes[0].set_xticks([0,25,50])
axes[0].set_yticks([0.2,0.5,0.8],[0.2,0.5,0.8])
axes[0].tick_params(direction='out', length=2, width=1 , labelsize=6 , pad=1)
 

axes[1].plot(range(n_epoch),sp_th_acc_mn,'tab:blue',range(n_epoch),nm_th_acc_mn,'tab:orange',linewidth=0.5)
axes[1].fill_between(range(n_epoch),sp_th_acc_mn-sp_th_acc_sd,sp_th_acc_mn+sp_th_acc_sd,facecolor='tab:blue',color=None,alpha=0.2)
axes[1].fill_between(range(n_epoch),nm_th_acc_mn-nm_th_acc_sd,nm_th_acc_mn+nm_th_acc_sd,facecolor='tab:orange',color=None,alpha=0.2)      
axes[1].spines['top'].set_visible(False)
axes[1].spines['right'].set_visible(False)
axes[1].spines['bottom'].set_edgecolor('black')
axes[1].spines['bottom'].set_facecolor('black')
axes[1].spines['bottom'].set_linewidth(0.5)
axes[1].spines['left'].set_edgecolor('black')
axes[1].spines['left'].set_facecolor('black')
axes[1].spines['left'].set_linewidth(0.5)
axes[1].set_xticks([0,25,50])
axes[1].set_ylim(0.7,0.9)
axes[1].set_yticks([0.7,0.8,0.9],[0.7,0.8,0.9])
axes[1].tick_params(direction='out', length=2, width=1 , labelsize=6 , pad=1)

plt.savefig('fig/Fig_3_e.pdf', format='pdf',bbox_inches='tight',pad_inches=0.25)


#%%

L2_cst=np.load('sim_data/Fig_3_L2_nm_cst.npy')
L2_acc=np.load('sim_data/Fig_3_L2_nm_acc.npy')
L0_cst=np.load('sim_data/Fig_3_L0_nm_cst.npy')
L0_acc=np.load('sim_data/Fig_3_L0_nm_acc.npy')
L2L0_cst=np.load('sim_data/Fig_3_L2L0_nm_cst.npy')
L2L0_acc=np.load('sim_data/Fig_3_L2L0_nm_acc.npy')


n_rep=L2_cst.shape[0]
n_epoch=L2_cst.shape[1]

L2_cst_mn=np.zeros(n_epoch)
L2_cst_sd=np.zeros(n_epoch)
L2_acc_mn=np.zeros(n_epoch)
L2_acc_sd=np.zeros(n_epoch)

L0_cst_mn=np.zeros(n_epoch)
L0_cst_sd=np.zeros(n_epoch)
L0_acc_mn=np.zeros(n_epoch)
L0_acc_sd=np.zeros(n_epoch)

L2L0_cst_mn=np.zeros(n_epoch)
L2L0_cst_sd=np.zeros(n_epoch)
L2L0_acc_mn=np.zeros(n_epoch)
L2L0_acc_sd=np.zeros(n_epoch)

for ep_ind in range(n_epoch):
    vec_L2_cst=L2_cst[:,ep_ind]
    L2_cst_mn[ep_ind]=np.mean(vec_L2_cst)
    L2_cst_sd[ep_ind]=(n_rep/(n_rep-1)*np.var(vec_L2_cst))**0.5
    
    vec_L2_acc=L2_acc[:,ep_ind]
    L2_acc_mn[ep_ind]=np.mean(vec_L2_acc)
    L2_acc_sd[ep_ind]=(n_rep/(n_rep-1)*np.var(vec_L2_acc))**0.5
    
    vec_L0_cst=L0_cst[:,ep_ind]
    L0_cst_mn[ep_ind]=np.mean(vec_L0_cst)
    L0_cst_sd[ep_ind]=(n_rep/(n_rep-1)*np.var(vec_L0_cst))**0.5
    
    vec_L0_acc=L0_acc[:,ep_ind]
    L0_acc_mn[ep_ind]=np.mean(vec_L0_acc)
    L0_acc_sd[ep_ind]=(n_rep/(n_rep-1)*np.var(vec_L0_acc))**0.5
    
    vec_L2L0_cst=L2L0_cst[:,ep_ind]
    L2L0_cst_mn[ep_ind]=np.mean(vec_L2L0_cst)
    L2L0_cst_sd[ep_ind]=(n_rep/(n_rep-1)*np.var(vec_L2L0_cst))**0.5
      
    vec_L2L0_acc=L2L0_acc[:,ep_ind]
    L2L0_acc_mn[ep_ind]=np.mean(vec_L2L0_acc)
    L2L0_acc_sd[ep_ind]=(n_rep/(n_rep-1)*np.var(vec_L2L0_acc))**0.5
         
sns.set(style="white", palette="muted", color_codes=True)
f, axes = plt.subplots(2, 1, figsize=(1.1811,1.2811), sharex=True)
sns.despine      
    
axes[0].plot(range(n_epoch),L0_cst_mn,'tab:orange',range(n_epoch),L2_cst_mn,'tab:green',L2L0_cst_mn,'tab:red',linewidth=0.5)
axes[0].fill_between(range(n_epoch),L0_cst_mn-L0_cst_sd,L0_cst_mn+L0_cst_sd,facecolor='tab:orange',color=None,alpha=0.2)
axes[0].fill_between(range(n_epoch),L2_cst_mn-L2_cst_sd,L2_cst_mn+L2_cst_sd,facecolor='tab:green',color=None,alpha=0.2) 
axes[0].fill_between(range(n_epoch),L2L0_cst_mn-L2L0_cst_sd,L2L0_cst_mn+L2L0_cst_sd,facecolor='tab:green',color=None,alpha=0.2)       
axes[0].spines['top'].set_visible(False)
axes[0].spines['right'].set_visible(False)
axes[0].spines['bottom'].set_edgecolor('black')
axes[0].spines['bottom'].set_facecolor('black')
axes[0].spines['bottom'].set_linewidth(0.5)
axes[0].spines['left'].set_edgecolor('black')
axes[0].spines['left'].set_facecolor('black')
axes[0].spines['left'].set_linewidth(0.5)
axes[0].set_xticks([0,25,50])
axes[0].set_yticks([0.2,0.5,0.8],[0.2,0.5,0.8])
axes[0].tick_params(direction='out', length=2, width=1 , labelsize=6 , pad=1)
 

axes[1].plot(range(n_epoch),L0_acc_mn,'tab:orange',range(n_epoch),L2_acc_mn,'tab:green',L2L0_acc_mn,'tab:red',linewidth=0.5)
axes[1].fill_between(range(n_epoch),L0_acc_mn-L0_acc_sd,L0_acc_mn+L0_acc_sd,facecolor='tab:orange',color=None,alpha=0.2)
axes[1].fill_between(range(n_epoch),L2_acc_mn-L2_acc_sd,L2_acc_mn+L2_acc_sd,facecolor='tab:green',color=None,alpha=0.2) 
axes[1].fill_between(range(n_epoch),L2L0_acc_mn-L2L0_acc_sd,L2L0_acc_mn+L2L0_acc_sd,facecolor='tab:green',color=None,alpha=0.2)       
axes[1].spines['top'].set_visible(False)
axes[1].spines['right'].set_visible(False)
axes[1].spines['bottom'].set_edgecolor('black')
axes[1].spines['bottom'].set_facecolor('black')
axes[1].spines['bottom'].set_linewidth(0.5)
axes[1].spines['left'].set_edgecolor('black')
axes[1].spines['left'].set_facecolor('black')
axes[1].spines['left'].set_linewidth(0.5)
axes[1].set_xticks([0,25,50])
axes[1].set_ylim(0.7,0.9)
axes[1].set_yticks([0.7,0.8,0.9],[0.7,0.8,0.9])
axes[1].tick_params(direction='out', length=2, width=1 , labelsize=6 , pad=1)

plt.savefig('fig/Fig_3_f.pdf', format='pdf',bbox_inches='tight',pad_inches=0.25)

#%%

const_accs=np.load('sim_data/Fig_3_Lp_const.npy')
var_accs=np.load('sim_data/Fig_3_Lp_var.npy')

n_rep=const_accs.shape[2]

const_acc_mn=np.zeros([6,4])
const_acc_sd=np.zeros([6,4])
var_acc_mn=np.zeros([6,4])
var_acc_sd=np.zeros([6,4])


for p_ind in range(6):
    for e_ind in range(4):
        vec_const_acc=const_accs[p_ind,e_ind]
        const_acc_mn[p_ind,e_ind]=np.mean(vec_const_acc)
        const_acc_sd[p_ind,e_ind]=(n_rep/(n_rep-1)*np.var(vec_const_acc))**0.5
     
        vec_var_acc=var_accs[p_ind,e_ind]
        var_acc_mn[p_ind,e_ind]=np.mean(vec_var_acc)
        var_acc_sd[p_ind,e_ind]=(n_rep/(n_rep-1)*np.var(vec_var_acc))**0.5

for ind in range(1,4):
    const_acc_mn[0,ind] , const_acc_mn[1,ind] = const_acc_mn[1,ind] , const_acc_mn[0,ind]
    const_acc_sd[0,ind] , const_acc_sd[1,ind] = const_acc_sd[1,ind] , const_acc_sd[0,ind]
      
sns.set(style="white", palette="muted", color_codes=True)
f, axes = plt.subplots(4, 1, figsize=(1.1811,1.2811), sharex=True)
sns.despine      

for ind in range(4):
    axes[ind].errorbar(range(6),const_acc_mn[:,ind],yerr=const_acc_sd[:,ind],color='tab:pink',linewidth=0.5)
    axes[ind].errorbar(range(6),var_acc_mn[:,ind],yerr=var_acc_sd[:,ind],color='tab:olive',linewidth=0.5)  
    axes[ind].spines['top'].set_visible(False)
    axes[ind].spines['right'].set_visible(False)
    axes[ind].spines['bottom'].set_edgecolor('black')
    axes[ind].spines['bottom'].set_facecolor('black')
    axes[ind].spines['bottom'].set_linewidth(0.5)
    axes[ind].spines['left'].set_edgecolor('black')
    axes[ind].spines['left'].set_facecolor('black')
    axes[ind].spines['left'].set_linewidth(0.5)
    axes[ind].set_xticks(range(6))
    axes[ind].set_yticks([0.75,0.8,0.85],[0.75,0.8,0.85])
    axes[ind].tick_params(direction='out', length=2, width=1 , labelsize=6 , pad=1)
 

plt.savefig('fig/Fig_3_g.pdf', format='pdf',bbox_inches='tight',pad_inches=0.25)
```
