## Supplementary material for "Dendritic normalisation improves learning in sparsely connected artificial neural networks": Code and data.: hidden_acts.py

```
# -*- coding: utf-8 -*-
"""
Created on Mon Nov 18 18:41:10 2019

@author: Alex Bird
"""

def input_to_hidden_layer(network, dataset):
    """Return the inputs to the hidden layer"""
    
    ndata=len(dataset)
    w=network.weights[0]
    b=network.biases[0]
    
    a=np.zeros([network.sizes[1],1])
    for ind in range(ndata):      
        ia = np.dot(w, dataset[ind][0])+b    
        a=a+ia
    a=a/ndata
        
    return a
```
