## Supplementary material for "Dendritic normalisation improves learning in sparsely connected artificial neural networks": Code and data.: import_all_data.py

```
#!/usr/bin/env python3
# -*- coding: utf-8 -*-
"""
Created on Mon Nov  4 18:31:17 2019

@author: Alex
"""
import mnist_loader

# MNIST - Digits
dig_training=[]
tr_images , tr_labels=load_mnist('data/','dig_train')
for ind in range(60000):
    dig_training.append([np.reshape(tr_images[ind],[784,1])/256 , vectorised_result(tr_labels[ind])])

dig_test=[]
tst_images , tst_labels=load_mnist('data/','dig_t10k')
for ind in range(10000):
    dig_test.append([np.reshape(tst_images[ind],[784,1])/256 , tst_labels[ind]])

# MNIST - Fashion
fash_training=[]
tr_images , tr_labels=load_mnist('data/','fash_train')
for ind in range(60000):
    fash_training.append([np.reshape(tr_images[ind],[784,1])/256 , vectorised_result(tr_labels[ind])])

fash_test=[]
tst_images , tst_labels=load_mnist('data/','fash_t10k')
for ind in range(10000):
    fash_test.append([np.reshape(tst_images[ind],[784,1] )/256, tst_labels[ind]])
```
