## Supplementary material for "Dendritic normalisation improves learning in sparsely connected artificial neural networks": Code and data.: mnist_loader.py

```
"""
mnist_loader
~~~~~~~~~~~~
"""

#### Libraries

def load_mnist(path, kind):
    import os
    import gzip
    import numpy as np

    """Load MNIST data from `path`"""
    labels_path = os.path.join(path,
                               '%s-labels-idx1-ubyte.gz'
                               % kind)
    images_path = os.path.join(path,
                               '%s-images-idx3-ubyte.gz'
                               % kind)

    with gzip.open(labels_path, 'rb') as lbpath:
        labels = np.frombuffer(lbpath.read(), dtype=np.uint8,
                               offset=8)

    with gzip.open(images_path, 'rb') as imgpath:
        images = np.frombuffer(imgpath.read(), dtype=np.uint8,
                               offset=16).reshape(len(labels), 784)

    return images, labels

def vectorised_result(j):
    """Return a 10-dimensional unit vector with a 1.0 in the j'th position
    and zeroes elsewhere.
    """
    e = np.zeros((10, 1))
    e[j] = 1.0
    return e
```
