## Supplementary material for "Dendritic normalisation improves learning in sparsely connected artificial neural networks": Code and data.: net_funcs.py

```
#!/usr/bin/env python3
# -*- coding: utf-8 -*-
"""
Created on Wed Dec 11 17:33:02 2019

@author: Alex
"""

def nm_func(ly_sz,n_epoch,training,test):
    sizes=[784]
    for l_ind in range(np.size(ly_sz)):
        sizes.append(ly_sz[l_ind])
    sizes.append(10)
    net_nm=network_nm.Network(sizes,0.2,0.15)
    [accuracy , cost] = net_nm.SGD(training,n_epoch,10,0.05,evaluation_data=test,show_progress=False)
    return accuracy , cost

def sp_func(ly_sz,n_epoch,training,test):
    sizes=[784]
    for l_ind in range(np.size(ly_sz)):
        sizes.append(ly_sz[l_ind])
    sizes.append(10)
    net_sp=network_sp.Network(sizes,0.2,0.15)
    [accuracy , cost] = net_sp.SGD(training,n_epoch,10,0.05,evaluation_data=test,show_progress=False)
    return accuracy , cost

def nm_thlin_func(ly_sz,n_epoch,training,test):
    sizes=[784]
    for l_ind in range(np.size(ly_sz)):
        sizes.append(ly_sz[l_ind])
    sizes.append(10)
    net_nm=network_nm_thlin.Network(sizes,0.2,0.15)
    [accuracy , cost] = net_nm.SGD(training,n_epoch,10,0.05,evaluation_data=test,show_progress=False)
    return accuracy , cost

def sp_thlin_func(ly_sz,n_epoch,training,test):
    sizes=[784]
    for l_ind in range(np.size(ly_sz)):
        sizes.append(ly_sz[l_ind])
    sizes.append(10)
    net_sp=network_sp_thlin.Network(sizes,0.2,0.15)
    [accuracy , cost] = net_sp.SGD(training,n_epoch,10,0.05,evaluation_data=test,show_progress=False)
    return accuracy , cost
```
