## Supplementary material for "Dendritic normalisation improves learning in sparsely connected artificial neural networks": Code and data.: network_nm.py

```
"""network_nm.py
~~~~~~~~~~~~~~
Neural network with dendritic normalisation.

"""

#### Libraries
import numpy as np


#### Define the cost function

class LogLikelihoodCost(object):

    @staticmethod
    def fn(a, y):
        return -np.log(a[np.nonzero(y)])

    @staticmethod
    def delta(z, a, y):
        return (a-y)


#### Main Network class
class Network(object):

    def __init__(self, sizes, con_prob, con_delt, s_init=25, cost=LogLikelihoodCost):
        """The list ``sizes`` contains the number of neurons in the respective
        layers of the network.  For example, if the list was [784, 30, 10]
        then it would be a three-layer network, with the first layer
        containing 784 neurons, the second layer 30 neurons, and the
        third layer 10 neurons. 'con_prob' gives the connection probability and 'con_delt' the proportion of weakest contacts which are excised at each epoch under the SET algorithm. 

        """
        self.num_layers = len(sizes)
        self.sizes = sizes
        self.s=s_init
        self.weight_initializer(con_prob)
        self.cost=cost
        self.con_delt=con_delt

    def weight_initializer(self,con_prob):
        s=self.s
        self.biases = [np.random.randn(y, 1) for y in self.sizes[1:]]     
        self.veights=[]
        self.weights=[]
        for l_ind in range(len(self.sizes)-2):
            tot_sz=self.sizes[l_ind]*self.sizes[l_ind+1]
            ivght=np.zeros([tot_sz,1])
            n_allowed=int(np.round(con_prob*tot_sz))
            ind_allowed=np.random.choice(tot_sz,n_allowed,replace=False)
            vals_allowed=np.random.randn(n_allowed,1)
            ivght[ind_allowed]=vals_allowed
            ivght=np.reshape(ivght,[self.sizes[l_ind+1],self.sizes[l_ind]])                        
            self.veights.append(ivght)
            
            n_aff=np.count_nonzero(ivght,1)
            mrph_nm=np.transpose(np.tile(n_aff,[self.sizes[l_ind],1]))
            iwght=s*ivght/mrph_nm
            iwght[np.isnan(iwght)]=0
            self.weights.append(iwght)
                
        ivght=np.random.randn(self.sizes[-1],self.sizes[-2])
        self.veights.append(ivght)
        self.weights.append(ivght)

    def feedforward(self, a):
        """Return the output of the network"""
        
        for b, w in zip(self.biases[:-1], self.weights[:-1]):
            a = sigmoid(np.dot(w, a)+b)
        
        lS=self.num_layers-2
        w=self.weights[lS]
        b=self.biases[lS]
        a=softmax(np.dot(w, a)+b)
        return a

    def SGD(self, training_data, epochs, mini_batch_size, eta,
            evaluation_data=None,
            show_progress=True):
        """Train the neural network using stochastic gradient
        descent with dendritic normalisation (Eq 7 in the paper) and the SET to regulate sparsity.

        """
        if evaluation_data: n_data = len(evaluation_data)
        n = len(training_data)
        evaluation_cost, evaluation_accuracy = [], []
        training_cost, training_accuracy = [], []
        
        con_delt=self.con_delt
        for j in range(epochs):
            np.random.shuffle(training_data)
            mini_batches = [training_data[k:k+mini_batch_size] for k in range(0, n, mini_batch_size)]
            for mini_batch in mini_batches:
                self.update_mini_batch(mini_batch, eta, len(training_data))
                     
            # SET step
            for l_ind in range(self.num_layers-2):
                vght=self.veights[l_ind]
                wght=self.weights[l_ind]
                totsz=np.size(wght)
                n_con=np.count_nonzero(wght)
                n_change=round(n_con*con_delt)
                w_vals=wght[np.nonzero(wght)]                
                w_abs=np.sort(np.abs(w_vals))
                w_thresh=w_abs[n_change]
                
                ex_vals=np.abs(wght)<w_thresh
                wght[ex_vals]=0 
                vght[ex_vals]=0
                            
                if j<epochs: # Add random new weights if not last run
                    n_con_2=np.count_nonzero(vght)
                    n_change=n_con-n_con_2
                    
                    pos_locs=np.nonzero(vght==0)
                    pos_rws=pos_locs[0]
                    pos_cls=pos_locs[1]
                
                    new_vs=np.random.randn(n_change)
                    new_inds=np.random.choice(np.size(pos_rws),n_change,replace=False)
                
                    vght[pos_rws[new_inds],pos_cls[new_inds]]=new_vs
                
                self.veights[l_ind]=vght
                
                n_aff=np.count_nonzero(vght,1)
                mrph_nm=np.transpose(np.tile(n_aff,[self.sizes[l_ind],1]))
                iwght=self.s*vght/mrph_nm
                iwght[np.isnan(iwght)]=0   
                self.weights[l_ind]=iwght
            
            cost = self.total_cost(training_data)
            training_cost.append(cost)
            accuracy = self.accuracy(evaluation_data)
            evaluation_accuracy.append(accuracy)
            
            if show_progress:
                print("Epoch %s training complete" % j)
                print("Cost on training data: {}".format(cost))
                print("Accuracy on evaluation data: {} / {}".format(self.accuracy(evaluation_data), n_data))                               
                
        return evaluation_accuracy, training_cost

    def update_mini_batch(self, mini_batch, eta, n):
        nabla_b = [np.zeros(b.shape) for b in self.biases]
        nabla_v = [np.zeros(w.shape) for w in self.weights]
        delta_s = 0
        for x, y in mini_batch:
            delta_nabla_b, delta_nabla_v, delta_delta_s = self.backprop(x, y)
            
            nabla_b = [nb+dnb for nb, dnb in zip(nabla_b, delta_nabla_b)]
            nabla_v = [nv+dnv for nv, dnv in zip(nabla_v, delta_nabla_v)]  
            delta_s = delta_s+delta_delta_s   
        
        self.biases = [b-(eta/len(mini_batch))*nb
                       for b, nb in zip(self.biases, nabla_b)]
        self.veights = [1-(eta/len(mini_batch))*nv
                        for v, nv in zip(self.veights, nabla_v)]
        
        for l_ind in range(self.num_layers-2):
            L0_nm=np.count_nonzero(self.veights[l_ind],1)
            mrph_nm=np.transpose(np.tile(L0_nm,[self.sizes[l_ind],1]))        
            self.weights[l_ind]=self.s*self.veights[l_ind]/mrph_nm
                     
        self.s=self.s-eta/len(mini_batch)*delta_s
        
       

    def backprop(self, x, y):
        nabla_b = [np.zeros(b.shape) for b in self.biases]
        nabla_w = [np.zeros(w.shape) for w in self.weights]
        nabla_v = [np.zeros(w.shape) for w in self.weights]
        
        s=self.s
        # feedforward
        activation = x
        activations = [x]
        zs = []
        
        
        for b, w in zip(self.biases[:-1], self.weights[:-1]):
             z = np.dot(w, activation)+b
             zs.append(z)
             activation = sigmoid(z)
             activations.append(activation)            
        
        lS=len(self.sizes)-2
        w=self.weights[lS]
        b=self.biases[lS]
        z = np.dot(w, activation)+b                
        zs.append(z)
        activation = softmax(z)
        activations.append(activation)
     
        # backward pass
        delta = (self.cost).delta(zs[-1], activations[-1], y)
        nabla_b[-1] = delta
        nabla_w[-1] = np.dot(delta, activations[-2].transpose())
        nabla_v[-1] = np.dot(delta, activations[-2].transpose())
        delt_s=0
        for l in range(2, self.num_layers):
            z = zs[-l]
            sp = sigmoid_prime(z)
            delta = np.dot(self.weights[-l+1].transpose(), delta) * sp
            nabla_b[-l] = delta
            msk=np.zeros(np.shape(self.weights[-l]))
            msk[np.nonzero(self.weights[-l])]=1            
            nabla_w[-l] = np.dot(delta, activations[-l-1].transpose())*msk
            iwght=nabla_w[-l]
            
            L0_nm=np.count_nonzero(iwght,1)
            mrph_nm=np.transpose(np.tile(L0_nm,[self.sizes[-l-1],1])) 
            ivght=s*iwght/mrph_nm
            ivght[np.isnan(ivght)]=0
            nabla_v[-l]=ivght
            
            s_cont=self.weights[-l]*nabla_w[-l]
            delt_s=delt_s+s_cont.sum()/s
            
            
        return (nabla_b, nabla_v , delt_s)

    def accuracy(self, data):
        results = [(np.argmax(self.feedforward(x)), y) for (x, y) in data]
        return sum(int(x == y) for (x, y) in results)

    def total_cost(self, data):
        cost = 0.0
        for x, y in data:
            a = self.feedforward(x)
            cost += self.cost.fn(a, y)/len(data)
        return cost

    
#### Miscellaneous functions

def sigmoid(z):
    return 1.0/(1.0+np.exp(-z))

def softmax(z):
    rawexp=np.exp(z)
    S=rawexp.sum()
    return rawexp/S

def sigmoid_prime(z):
    return sigmoid(z)*(1-sigmoid(z))
```
