## Supplementary material for "Dendritic normalisation improves learning in sparsely connected artificial neural networks": Code and data.: network_nm_2lyr.py

```
#!/usr/bin/env python3
# -*- coding: utf-8 -*-
"""
Created on Wed Nov 27 00:37:07 2019

@author: Alex
"""


from __future__ import absolute_import
from __future__ import division
from __future__ import print_function

import numpy as np

import tensorflow as tf
import copy
import types as python_types
import warnings

from keras import backend as K
from keras import activations
from keras import initializers
from keras import regularizers
from keras import constraints
from keras.engine.base_layer import InputSpec
from keras.engine.base_layer import Layer
from keras.utils.generic_utils import func_dump
from keras.utils.generic_utils import func_load
from keras.utils.generic_utils import deserialize_keras_object
from keras.utils.generic_utils import has_arg
from keras.legacy import interfaces
from keras.models import Sequential
from keras.layers import Dense, Activation, Flatten, Conv2D, MaxPooling2D
from keras import optimizers


class Morph_layer(Layer):
    @interfaces.legacy_dense_support
    def __init__(self, units,
                 activation=None,
                 use_bias=True,
                 kernel_initializer='glorot_uniform',
                 bias_initializer='zeros',
                 kernel_regularizer=None,
                 bias_regularizer=None,
                 activity_regularizer=None,
                 kernel_constraint=None,
                 bias_constraint=None,
                 **kwargs):
        if 'input_shape' not in kwargs and 'input_dim' in kwargs:
            kwargs['input_shape'] = (kwargs.pop('input_dim'),)
        super(Morph_layer, self).__init__(**kwargs)
        self.units = units
        self.activation = activations.get(activation)
        self.use_bias = use_bias
        self.kernel_initializer = initializers.get(kernel_initializer)
        self.bias_initializer = initializers.get(bias_initializer)
        self.kernel_regularizer = regularizers.get(kernel_regularizer)
        self.bias_regularizer = regularizers.get(bias_regularizer)
        self.activity_regularizer = regularizers.get(activity_regularizer)
        self.kernel_constraint = constraints.get(kernel_constraint)
        self.bias_constraint = constraints.get(bias_constraint)
        self.input_spec = InputSpec(min_ndim=2)
        self.supports_masking = True

    def build(self, input_shape):
        assert len(input_shape) >= 2
        input_dim = input_shape[-1]

        self.kernel = self.add_weight(shape=(input_dim, self.units),
                                      initializer=self.kernel_initializer,
                                      name='kernel',
                                      regularizer=self.kernel_regularizer,
                                      constraint=self.kernel_constraint)
        if self.use_bias:
            self.bias = self.add_weight(shape=(self.units,),
                                        initializer=self.bias_initializer,
                                        name='bias',
                                        regularizer=self.bias_regularizer,
                                        constraint=self.bias_constraint)
        else:
            self.bias = None
        self.s=self.add_weight(shape=(1,),
                                        initializer=self.bias_initializer,
                                        name='s')  
        
        self.input_spec = InputSpec(min_ndim=2, axes={-1: input_dim})
        self.built = True

    def call(self, inputs):
        output = K.dot(inputs, self.kernel)
        if self.use_bias:
            output = K.bias_add(output, self.bias, data_format='channels_last')
        if self.activation is not None:
            output = self.activation(output)
        return output

    def compute_output_shape(self, input_shape):
        assert input_shape and len(input_shape) >= 2
        assert input_shape[-1]
        output_shape = list(input_shape)
        output_shape[-1] = self.units
        return tuple(output_shape)

    def get_config(self):
        config = {
            'units': self.units,
            'activation': activations.serialize(self.activation),
            'use_bias': self.use_bias,
            'kernel_initializer': initializers.serialize(self.kernel_initializer),
            'bias_initializer': initializers.serialize(self.bias_initializer),
            'kernel_regularizer': regularizers.serialize(self.kernel_regularizer),
            'bias_regularizer': regularizers.serialize(self.bias_regularizer),
            'activity_regularizer':
                regularizers.serialize(self.activity_regularizer),
            'kernel_constraint': constraints.serialize(self.kernel_constraint),
            'bias_constraint': constraints.serialize(self.bias_constraint)
        }
        base_config = super(Morph_layer, self).get_config()
        return dict(list(base_config.items()) + list(config.items()))
#%%

from keras.optimizers import Optimizer
from keras.legacy import interfaces

class Morpho_SGD(Optimizer):     
    def __init__(self, learning_rate=0.05, **kwargs):
        learning_rate = kwargs.pop('lr', learning_rate)
        super(Morpho_SGD, self).__init__(**kwargs)
        with K.name_scope(self.__class__.__name__):
            self.iterations = K.variable(0, dtype='int64', name='iterations')
            self.learning_rate = K.variable(learning_rate, name='learning_rate')
        
    @interfaces.legacy_get_updates_support
    @K.symbolic
    def get_updates(self, loss, params):
        grads=[]
        grads.append(self.get_gradients(loss, params[0]))
        grads.append(self.get_gradients(loss, params[1]))
        grads.append(0)
        grads.append(self.get_gradients(loss, params[3]))
        grads.append(self.get_gradients(loss, params[4]))
        grads.append(0)
        grads.append(self.get_gradients(loss, params[6]))
        grads.append(self.get_gradients(loss, params[7]))
        self.updates = [K.update_add(self.iterations, 1)]

        weights=params
        grmn_1=grads[0]
        wght_1=weights[0]
        s_1=weights[2]
       
        grmn_2=grads[3]
        wght_2=weights[3]
        s_2=weights[5]
        
        n_all= K.not_equal(wght_1, 0.0)
        n_aff=K.sum(K.cast(n_all,'float32'),axis=0)
        n_aff=K.expand_dims(n_aff,axis=1)
        mrph_nm=K.transpose(K.tile(n_aff,784))  
        n_grad_1=tf.math.divide_no_nan(s_1*grmn_1,mrph_nm)       
        
        n_all= K.not_equal(wght_2, 0.0)
        n_aff=K.sum(K.cast(n_all,'float32'),axis=0)
        n_aff=K.expand_dims(n_aff,axis=1)
        mrph_nm=K.transpose(K.tile(n_aff,100))
        n_grad_2=tf.math.divide_no_nan(s_2*grmn_2,mrph_nm)
        
        s_delt=(K.sum(grmn_1*wght_1)+K.sum(grmn_2*wght_2))/s_1
        
        grads[0]=n_grad_1
        grads[2]=K.expand_dims(s_delt,axis=0)
        grads[3]=n_grad_2
        grads[5]=K.expand_dims(s_delt,axis=0)


        lr = self.learning_rate       
        for p, g in zip(params, grads):
            v = - lr * g           
            new_p = p + v
            new_p=K.squeeze(new_p,axis=0)
            if getattr(p, 'constraint', None) is not None:
                new_p = p.constraint(new_p)
            self.updates.append(K.update(p, new_p))
        return self.updates

    def get_config(self):
        config = {'learning_rate': float(K.get_value(self.learning_rate))}
        base_config = super(Morpho_SGD, self).get_config()
        return dict(list(base_config.items()) + list(config.items()))

    
#%%

class Constraint(object):

    def __call__(self, w):
        return w

    def get_config(self):
        return {}

class MaskWeights(Constraint):

    def __init__(self, mask):
        self.mask = mask
        self.mask = K.cast(self.mask, K.floatx())

    def __call__(self, w):
        w *= self.mask
        return w

    def get_config(self):
        return {'mask': self.mask}


con_prob=0.2
con_delt=0.15
s=[25]

units=100
input_dim=784

tot_sz=units*input_dim
iwght=np.zeros([tot_sz,1])
iker=np.zeros([tot_sz,1])
n_allowed=int(np.round(con_prob*tot_sz))
ind_allowed=np.random.choice(tot_sz,n_allowed,replace=False)
vals_allowed=np.random.randn(n_allowed,1)
ker_allowed=np.ones((n_allowed,1))
iwght[ind_allowed]=vals_allowed
iker[ind_allowed]=ker_allowed

veight=np.reshape(iwght,[input_dim,units])
non_sparse_1=np.reshape(iker,[input_dim,units])

n_aff=np.count_nonzero(veight,0)
mrph_nm=np.tile(n_aff,[784,1])
weight_1=s*veight/mrph_nm
weight_1[np.isnan(weight)]=0

############

tot_sz=units*units
iwght=np.zeros([tot_sz,1])
iker=np.zeros([tot_sz,1])
n_allowed=int(np.round(con_prob*tot_sz))
ind_allowed=np.random.choice(tot_sz,n_allowed,replace=False)
vals_allowed=np.random.randn(n_allowed,1)
ker_allowed=np.ones((n_allowed,1))
iwght[ind_allowed]=vals_allowed
iker[ind_allowed]=ker_allowed

veight=np.reshape(iwght,[units,units])
non_sparse_2=np.reshape(iker,[units,units])

n_aff=np.count_nonzero(veight,0)
mrph_nm=np.tile(n_aff,[100,1])
weight_2=s*veight/mrph_nm
weight_2[np.isnan(weight_2)]=0

fin_wght_vals=np.random.randn(10*units,1)
fin_wghts=np.reshape(fin_wght_vals,[units,10])

model_nm = Sequential()
model_nm.add(Flatten(input_shape=(28,28,1)))
model_nm.add(Morph_layer(units,kernel_constraint=MaskWeights(non_sparse_1),activation='sigmoid'))
model_nm.add(Morph_layer(units,kernel_constraint=MaskWeights(non_sparse_2),activation='sigmoid'))
model_nm.add(Dense(10, activation='softmax'))

wghtv=model_nm.get_weights()
wghtv[0]=weight_1
wghtv[2]=s
wghtv[3]=weight_2
wghtv[5]=s
wghtv[6]=fin_wghts
model_nm.set_weights(wghtv)

(x_train, y_train), (x_test, y_test) = tf.keras.datasets.fashion_mnist.load_data()
x_train = x_train.astype('float32') / 255
x_test = x_test.astype('float32') / 255
x_train = x_train.reshape(-1,28, 28, 1)
x_test = x_test.reshape(-1,28, 28, 1)


#%%
epochs=50
eta=0.05
accuracies=np.zeros(epochs)
costs=np.zeros(epochs)
for epoch in range(0,epochs):
    model_nm.compile(loss='sparse_categorical_crossentropy', optimizer=Morpho_SGD(0.05), metrics=['accuracy'])
    hist=model_nm.fit(x=x_train,y=y_train,validation_data=(x_test,y_test),batch_size=10,verbose=0,epochs=1)
    
    costs[epoch]=hist.history['loss'][0]
    accuracies[epoch]=hist.history['val_accuracy'][0]
    
    print(accuracies[epoch])
    wght_vc=model_nm.get_weights()
    wght_1=wght_vc[0]
    s=wght_vc[2]
    wght_2=wght_vc[3]
    
    n_con_1=np.count_nonzero(wght_1)
    n_change=round(n_con_1*con_delt)
    w_vals=wght[np.nonzero(wght_1)]
    w_abs=np.sort(np.abs(w_vals))
    w_thresh=w_abs[n_change]
    wght_1[np.abs(wght_1)<w_thresh]=0
           
    n_con_2=np.count_nonzero(wght_2)
    n_change=round(n_con_2*con_delt)
    w_vals=wght[np.nonzero(wght_2)]
    w_abs=np.sort(np.abs(w_vals))
    w_thresh=w_abs[n_change]
    wght_2[np.abs(wght_2)<w_thresh]=0 
                      
    if epoch<epochs: # Add random new weights if not last run
        n_con_12=np.count_nonzero(wght_1)
        n_change=n_con_1-n_con_12                  
        pos_locs=np.nonzero(wght==0)
        pos_rws=pos_locs[0]
        pos_cls=pos_locs[1]                
        new_ws=np.random.randn(n_change)
        new_inds=np.random.choice(np.size(pos_rws),n_change,replace=False)                
        wght_1[pos_rws[new_inds],pos_cls[new_inds]]=new_ws
        
        wght_ker_inds=np.nonzero(wght_1)
        wght_ker_vals=np.ones(np.count_nonzero(wght_1))
        wght_ker_1=np.zeros([np.size(wght_1,0),np.size(wght_1,1)])
        wght_ker_1[wght_ker_inds]=wght_ker_vals
        
        veight_1=wght_1/s
        n_aff_pst=np.count_nonzero(veight_1,0)
        n_aff_pst[n_aff_pst==0]=1
        mrph_nm_pst=np.tile(n_aff_pst,[784,1])
        weight_1=s*veight_1/mrph_nm_pst
        weight_1[np.isnan(weight_1)]=0
        wght_vc[0]=weight_1   
        
        #####################
        n_con_22=np.count_nonzero(wght_2)
        n_change=n_con_2-n_con_22
        n_aff_pre=np.count_nonzero(wght_2,0)
        mrph_nm_pre=np.tile(n_aff_pre,[100,1])
                 
        pos_locs=np.nonzero(wght_2==0)
        pos_rws=pos_locs[0]
        pos_cls=pos_locs[1]                
        new_ws=np.random.randn(n_change)
        new_inds=np.random.choice(np.size(pos_rws),n_change,replace=False)                
        wght_2[pos_rws[new_inds],pos_cls[new_inds]]=new_ws
        
        wght_ker_inds=np.nonzero(wght_2)
        wght_ker_vals=np.ones(np.count_nonzero(wght_2))
        wght_ker_2=np.zeros([np.size(wght_2,0),np.size(wght_2,1)])
        wght_ker_2[wght_ker_inds]=wght_ker_vals
        
        veight_2=wght_2*mrph_nm_pre/s
        n_aff_pst=np.count_nonzero(veight_2,0)
        n_aff_pst[n_aff_pst==0]=1
        mrph_nm_pst=np.tile(n_aff_pst,[100,1])
        weight_2=s*veight_2/mrph_nm_pst
        weight_2[np.isnan(weight_2)]=0
        wght_vc[3]=weight_2   
        
        
        ###################
    
        model_nm = Sequential()
        model_nm.add(Flatten(input_shape=(28,28,1)))
        model_nm.add(Morph_layer(units,kernel_constraint=MaskWeights(wght_ker_1),activation='sigmoid'))
        model_nm.add(Morph_layer(units,kernel_constraint=MaskWeights(wght_ker_2),activation='sigmoid'))
        model_nm.add(Dense(10, activation='softmax'))

        model_nm.set_weights(wght_vc)
        
```
