## Supplementary material for "Dendritic normalisation improves learning in sparsely connected artificial neural networks": Code and data.: network_nm_L2.py

```
#!/usr/bin/env python3
# -*- coding: utf-8 -*-
"""
Created on Wed Nov 27 00:37:07 2019

@author: Alex
"""


from __future__ import absolute_import
from __future__ import division
from __future__ import print_function

import numpy as np

import copy
import types as python_types
import warnings

from keras import backend as K
from keras import activations
from keras import initializers
from keras import regularizers
from keras import constraints
from keras.engine.base_layer import InputSpec
from keras.engine.base_layer import Layer
from keras.utils.generic_utils import func_dump
from keras.utils.generic_utils import func_load
from keras.utils.generic_utils import deserialize_keras_object
from keras.utils.generic_utils import has_arg
from keras.legacy import interfaces

import tensorflow as tf
from keras.models import Sequential
from keras.layers import Dense, Activation, Flatten, Conv2D, MaxPooling2D
from keras import optimizers


class L2_layer(Layer):
    @interfaces.legacy_dense_support
    def __init__(self, units,
                 activation=None,
                 use_bias=True,
                 kernel_initializer='glorot_uniform',
                 bias_initializer='zeros',
                 kernel_regularizer=None,
                 bias_regularizer=None,
                 activity_regularizer=None,
                 kernel_constraint=None,
                 bias_constraint=None,
                 **kwargs):
        if 'input_shape' not in kwargs and 'input_dim' in kwargs:
            kwargs['input_shape'] = (kwargs.pop('input_dim'),)
        super(L2_layer, self).__init__(**kwargs)
        self.units = units
        self.activation = activations.get(activation)
        self.use_bias = use_bias
        self.kernel_initializer = initializers.get(kernel_initializer)
        self.bias_initializer = initializers.get(bias_initializer)
        self.kernel_regularizer = regularizers.get(kernel_regularizer)
        self.bias_regularizer = regularizers.get(bias_regularizer)
        self.activity_regularizer = regularizers.get(activity_regularizer)
        self.kernel_constraint = constraints.get(kernel_constraint)
        self.bias_constraint = constraints.get(bias_constraint)
        self.input_spec = InputSpec(min_ndim=2)
        self.supports_masking = True

    def build(self, input_shape):
        assert len(input_shape) >= 2
        input_dim = input_shape[-1]

        self.kernel = self.add_weight(shape=(input_dim, self.units),
                                      initializer=self.kernel_initializer,
                                      name='kernel',
                                      regularizer=self.kernel_regularizer,
                                      constraint=self.kernel_constraint)
        if self.use_bias:
            self.bias = self.add_weight(shape=(self.units,),
                                        initializer=self.bias_initializer,
                                        name='bias',
                                        regularizer=self.bias_regularizer,
                                        constraint=self.bias_constraint)
        else:
            self.bias = None
        self.g=self.add_weight(shape=(self.units,),
                                        initializer=self.kernel_initializer,
                                        name='g',
                                        regularizer=self.kernel_regularizer,
                                        constraint=self.kernel_constraint)  
        
        self.input_spec = InputSpec(min_ndim=2, axes={-1: input_dim})
        self.built = True

    def call(self, inputs):
        output = K.dot(inputs, self.kernel)
        if self.use_bias:
            output = K.bias_add(output, self.bias, data_format='channels_last')
        if self.activation is not None:
            output = self.activation(output)
        return output

    def compute_output_shape(self, input_shape):
        assert input_shape and len(input_shape) >= 2
        assert input_shape[-1]
        output_shape = list(input_shape)
        output_shape[-1] = self.units
        return tuple(output_shape)

    def get_config(self):
        config = {
            'units': self.units,
            'activation': activations.serialize(self.activation),
            'use_bias': self.use_bias,
            'kernel_initializer': initializers.serialize(self.kernel_initializer),
            'bias_initializer': initializers.serialize(self.bias_initializer),
            'kernel_regularizer': regularizers.serialize(self.kernel_regularizer),
            'bias_regularizer': regularizers.serialize(self.bias_regularizer),
            'activity_regularizer':
                regularizers.serialize(self.activity_regularizer),
            'kernel_constraint': constraints.serialize(self.kernel_constraint),
            'bias_constraint': constraints.serialize(self.bias_constraint)
        }
        base_config = super(L2_layer, self).get_config()
        return dict(list(base_config.items()) + list(config.items()))
#%%

from keras.optimizers import Optimizer
from keras.legacy import interfaces

class L2_SGD(Optimizer):     
    def __init__(self, learning_rate=0.05, **kwargs):
        learning_rate = kwargs.pop('lr', learning_rate)
        super(L2_SGD, self).__init__(**kwargs)
        with K.name_scope(self.__class__.__name__):
            self.iterations = K.variable(0, dtype='int64', name='iterations')
            self.learning_rate = K.variable(learning_rate, name='learning_rate')
        
    @interfaces.legacy_get_updates_support
    @K.symbolic
    def get_updates(self, loss, params):
        grads=[]
        grads.append(self.get_gradients(loss, params[0]))
        grads.append(self.get_gradients(loss, params[1]))
        grads.append(0)
        grads.append(self.get_gradients(loss, params[3]))
        grads.append(self.get_gradients(loss, params[4]))
        self.updates = [K.update_add(self.iterations, 1)]

        weights=params
        grmn=grads[0]
        wght=weights[0]
        g=weights[2]
        n_aff_L2=K.sqrt(K.sum(K.square(wght),axis=0))
        #n_aff_L2=K.expand_dims(n_aff_L2,axis=1)
        #L2_nm=K.transpose(K.tile(n_aff_L2,784))
        g_delt=K.batch_dot(K.transpose(grmn[0]),K.transpose(K.cast(wght, dtype='float32')),axes=1)/K.expand_dims(n_aff_L2,axis=1)
        term_1=g/n_aff_L2
        term_2=g*K.squeeze(g_delt,axis=1)/K.square(n_aff_L2)
        n_grad=term_1*grmn-term_2*wght
        
        #grads[0]=K.squeeze(n_grad,axis=0)
        #grads[2]=K.squeeze(g_delt,axis=1)
        grads[0]=n_grad
        grads[2]=K.transpose(g_delt)

        lr = self.learning_rate       
        for p, g in zip(params, grads):
            v = - lr * g           
            new_p = p + v
            new_p=K.squeeze(new_p,axis=0)
            #if getattr(p, 'constraint', None) is not None:
             #   new_p = p.constraint(new_p)
            self.updates.append(K.update(p, new_p))
        return self.updates

    def get_config(self):
        config = {'learning_rate': float(K.get_value(self.learning_rate))}
        base_config = super(L2_SGD, self).get_config()
        return dict(list(base_config.items()) + list(config.items()))

    
#%%

class Constraint(object):

    def __call__(self, w):
        return w

    def get_config(self):
        return {}

class MaskWeights(Constraint):

    def __init__(self, mask):
        self.mask = mask
        self.mask = K.cast(self.mask, K.floatx())

    def __call__(self, w):
        w *= self.mask
        return w

    def get_config(self):
        return {'mask': self.mask}


con_prob=0.2
con_delt=0.15
g_init=25

units=100
input_dim=784

tot_sz=units*input_dim
iwght=np.zeros([tot_sz,1])
iker=np.zeros([tot_sz,1])
n_allowed=int(np.round(con_prob*tot_sz))
ind_allowed=np.random.choice(tot_sz,n_allowed,replace=False)
vals_allowed=np.random.randn(n_allowed,1)
ker_allowed=np.ones((n_allowed,1))
iwght[ind_allowed]=vals_allowed
iker[ind_allowed]=ker_allowed

veight=np.reshape(iwght,[input_dim,units])
non_sparse=np.reshape(iker,[input_dim,units])

n_aff_L2=np.sqrt(np.sum(np.square(veight),0))
L2_nm=np.tile(n_aff_L2,[784,1])
weight=g_init*veight/L2_nm
weight[np.isnan(weight)]=0

fin_wght_vals=np.random.randn(10*units,1)
fin_wghts=np.reshape(fin_wght_vals,[units,10])

model_nm = Sequential()
model_nm.add(Flatten(input_shape=(28,28,1)))
model_nm.add(L2_layer(units,kernel_constraint=MaskWeights(non_sparse),activation='sigmoid'))
model_nm.add(Dense(10, activation='softmax'))


wghtv=model_nm.get_weights()
wghtv[0]=weight
wghtv[2]=g_init*np.ones(units)
wghtv[3]=fin_wghts
model_nm.set_weights(wghtv)

(x_train, y_train), (x_test, y_test) = tf.keras.datasets.fashion_mnist.load_data()
x_train = x_train.astype('float32') / 255
x_test = x_test.astype('float32') / 255
x_train = x_train.reshape(-1,28, 28, 1)
x_test = x_test.reshape(-1,28, 28, 1)


#%%
epochs=5
eta=0.05
accuracies=np.zeros(epochs)
costs=np.zeros(epochs)
for epoch in range(0,epochs):
    model_nm.compile(loss='sparse_categorical_crossentropy', optimizer=L2_SGD(0.05), metrics=['accuracy'])
    hist=model_nm.fit(x=x_train,y=y_train,validation_data=(x_test,y_test),batch_size=10,verbose=0,epochs=1)
    
    costs[epoch]=hist.history['loss'][0]
    accuracies[epoch]=hist.history['val_accuracy'][0]
    
    print(accuracies[epoch])
    wght_vc=model_nm.get_weights()
    wght=wght_vc[0]
    g_vec=wght_vc[2]
    
    n_con=np.count_nonzero(wght)
    n_change=round(n_con*con_delt)
    w_vals=wght[np.nonzero(wght)]
    w_abs=np.sort(np.abs(w_vals))
    w_thresh=w_abs[n_change]
    wght[np.abs(wght)<w_thresh]=0
                                       
    if epoch<epochs: # Add random new weights if not last run
        n_con_2=np.count_nonzero(wght)
        n_change=n_con-n_con_2
                    
        pos_locs=np.nonzero(wght==0)
        pos_rws=pos_locs[0]
        pos_cls=pos_locs[1]
                
        new_ws=np.random.randn(n_change)
        new_inds=np.random.choice(np.size(pos_rws),n_change,replace=False)
                
        wght[pos_rws[new_inds],pos_cls[new_inds]]=new_ws
        
        wght_ker_inds=np.nonzero(wght)
        wght_ker_vals=np.ones(np.count_nonzero(wght))
        wght_ker=np.zeros([np.size(wght,0),np.size(wght,1)])
        wght_ker[wght_ker_inds]=wght_ker_vals
        
        veight=wght/g_vec        
        n_aff_L2=np.sqrt(np.sum(np.square(veight),0))
        L2_nm=np.tile(n_aff_L2,[784,1])
        weight=g_vec*veight/L2_nm
        weight[np.isnan(weight)]=0
        wght_vc[0]=weight     
    
        model_nm = Sequential()
        model_nm.add(Flatten(input_shape=(28,28,1)))
        model_nm.add(L2_layer(units,kernel_constraint=MaskWeights(wght_ker),activation='sigmoid'))
        model_nm.add(Dense(10, activation='softmax'))
        
        model_nm.set_weights(wght_vc)
        
```
