## Supplementary material for "Dendritic normalisation improves learning in sparsely connected artificial neural networks": Code and data.: network_sp_2lyr.py

```
#!/usr/bin/env python3
# -*- coding: utf-8 -*-
"""
Created on Wed Nov 27 00:37:07 2019

import @author: Alex
"""

from keras.models import Sequential
from keras.layers import Dense, Activation, Flatten, Conv2D, MaxPooling2D
from keras import optimizers

from keras import backend as K


class Constraint(object):

    def __call__(self, w):
        return w

    def get_config(self):
        return {}

class MaskWeights(Constraint):

    def __init__(self, mask):
        self.mask = mask
        self.mask = K.cast(self.mask, K.floatx())

    def __call__(self, w):
        w *= self.mask
        return w

    def get_config(self):
        return {'mask': self.mask}


con_prob=0.2
con_delt=0.15

units=100
input_dim=784

tot_sz=units*input_dim
iwght=np.zeros([tot_sz,1])
iker=np.zeros([tot_sz,1])
n_allowed=int(np.round(con_prob*tot_sz))
ind_allowed=np.random.choice(tot_sz,n_allowed,replace=False)
vals_allowed=np.random.randn(n_allowed,1)
ker_allowed=np.ones((n_allowed,1))
iwght[ind_allowed]=vals_allowed
iker[ind_allowed]=ker_allowed
weight_in=np.reshape(iwght,[input_dim,units])
non_sparse_1=np.reshape(iker,[input_dim,units])

tot_sz=units*units
iwght=np.zeros([tot_sz,1])
iker=np.zeros([tot_sz,1])
n_allowed=int(np.round(con_prob*tot_sz))
ind_allowed=np.random.choice(tot_sz,n_allowed,replace=False)
vals_allowed=np.random.randn(n_allowed,1)
ker_allowed=np.ones((n_allowed,1))
iwght[ind_allowed]=vals_allowed
iker[ind_allowed]=ker_allowed
weight_mid=np.reshape(iwght,[units,units])
non_sparse_2=np.reshape(iker,[units,units])


fin_wght_vals=np.random.randn(10*units,1)
fin_wghts=np.reshape(fin_wght_vals,[units,10])

model_sp = Sequential()
model_sp.add(Flatten(input_shape=(28,28,1)))
model_sp.add(Dense(units,kernel_constraint=MaskWeights(non_sparse_1),activation='sigmoid'))
model_sp.add(Dense(units,kernel_constraint=MaskWeights(non_sparse_2),activation='sigmoid'))
model_sp.add(Dense(10, activation='softmax'))

wghtv=model_sp.get_weights()
wghtv[0]=weight_in
wghtv[2]=weight_mid
wghtv[4]=fin_wghts
model_sp.set_weights(wghtv)

(x_train, y_train), (x_test, y_test) = tf.keras.datasets.fashion_mnist.load_data()
x_train = x_train.astype('float32') / 255
x_test = x_test.astype('float32') / 255
x_train = x_train.reshape(-1,28, 28, 1)   #Reshape for CNN -  should work!!
x_test = x_test.reshape(-1,28, 28, 1)

#np.nonzero(fash_training[ind][1])[0][0]
#fash_training_imag=np.zeros([len(fash_training),784])
#fash_training_lab=np.zeros([len(fash_training),1])
#for ind in range(len(fash_training)):
#    fash_training_imag[ind,:]=np.reshape(fash_training[ind][0],784)
#    fash_training_lab[ind]=np.nonzero(fash_training[ind][1])[0][0]
#    
#fash_test_imag=np.zeros([len(fash_test),784])
#fash_test_lab=np.zeros([len(fash_test),1])
#for ind in range(len(fash_test)):
#    fash_test_imag[ind,:]=np.reshape(fash_test[ind][0],784)
#    fash_test_lab[ind]=fash_test[ind][1]

#%%
epochs=50
eta=0.05
accuracies=np.zeros(epochs)
costs=np.zeros(epochs)
for epoch in range(0,epochs):
    sgd = optimizers.SGD(eta, momentum=0,nesterov=False)
    model_sp.compile(loss='sparse_categorical_crossentropy', optimizer=sgd, metrics=['accuracy'])
    hist=model_sp.fit(x=x_train,y=y_train,validation_data=(x_test,y_test),batch_size=10,verbose=0,epochs=1)
    
    costs[epoch]=hist.history['loss'][0]
    accuracies[epoch]=hist.history['val_accuracy'][0]
    
    print(accuracies[epoch])
    wght_vc=model_sp.get_weights()
    wght_1=wght_vc[0]
    wght_2=wght_vc[2]
    
    n_con_1=np.count_nonzero(wght_1)
    n_change_1=round(n_con_1*con_delt)
    w_vals=wght_1[np.nonzero(wght_1)]
    w_abs=np.sort(np.abs(w_vals))
    w_thresh=w_abs[n_change_1]
    wght_1[np.abs(wght_1)<w_thresh]=0            
    
    n_con_2=np.count_nonzero(wght_2)
    n_change_2=round(n_con_2*con_delt)
    w_vals=wght_2[np.nonzero(wght_2)]
    w_abs=np.sort(np.abs(w_vals))
    w_thresh=w_abs[n_change_2]
    wght_2[np.abs(wght_2)<w_thresh]=0 
                        
    if epoch<epochs: # Add random new weights if not last run
        n_con_12=np.count_nonzero(wght_1)
        n_change=n_con_1-n_con_12
                    
        pos_locs=np.nonzero(wght_1==0)
        pos_rws=pos_locs[0]
        pos_cls=pos_locs[1]
                
        new_ws=np.random.randn(n_change)
        new_inds=np.random.choice(np.size(pos_rws),n_change,replace=False)
                
        wght_1[pos_rws[new_inds],pos_cls[new_inds]]=new_ws
        wght_vc[0]=wght_1
        
        wght_ker_inds=np.nonzero(wght_1)
        wght_ker_vals=np.ones(np.count_nonzero(wght_1))
        wght_ker_1=np.zeros([np.size(wght_1,0),np.size(wght_1,1)])
        wght_ker_1[wght_ker_inds]=wght_ker_vals
        
        #############################
        n_con_22=np.count_nonzero(wght_2)
        n_change=n_con_2-n_con_22
                    
        pos_locs=np.nonzero(wght_2==0)
        pos_rws=pos_locs[0]
        pos_cls=pos_locs[1]
                
        new_ws=np.random.randn(n_change)
        new_inds=np.random.choice(np.size(pos_rws),n_change,replace=False)
                
        wght_2[pos_rws[new_inds],pos_cls[new_inds]]=new_ws
        wght_vc[2]=wght_2
        
        wght_ker_inds=np.nonzero(wght_2)
        wght_ker_vals=np.ones(np.count_nonzero(wght_2))
        wght_ker_2=np.zeros([np.size(wght_2,0),np.size(wght_2,1)])
        wght_ker_2[wght_ker_inds]=wght_ker_vals
        
    
        model_sp = Sequential()
        model_sp.add(Flatten(input_shape=(28,28,1)))
        model_sp.add(Dense(units,kernel_constraint=MaskWeights(wght_ker_1),activation='sigmoid'))
        model_sp.add(Dense(units,kernel_constraint=MaskWeights(wght_ker_2),activation='sigmoid'))
        model_sp.add(Dense(10, activation='softmax'))
        
        model_sp.set_weights(wght_vc)
        
               
 
```
